# A neural substrate of sex-dependent modulation of motivation by value

**DOI:** 10.1101/2022.07.07.499209

**Authors:** Julia Cox, Adelaide R. Minerva, Weston T. Fleming, Christopher A. Zimmerman, Cameron Hayes, Samuel Zorowitz, Akhil Bandi, Sharon Ornelas, Brenna McMannon, Nathan F. Parker, Ilana B. Witten

## Abstract

While there is emerging evidence of sex differences in decision-making behavior, the neural substrates that underlie such differences remain largely unknown. Here, we demonstrate that in mice performing a value-based decision-making task, while choices are similar between the sexes, motivation to engage in the task is modulated by action value in females more strongly than in males. Inhibition of activity in anterior cingulate cortex (ACC) neurons that project to the dorsomedial striatum (DMS) disrupts this relationship between value and motivation preferentially in females, without affecting choice in either sex. In line with these effects, in females compared to males, ACC-DMS neurons have stronger representations of negative outcomes, and more neurons are active when the value of the chosen option is low. In contrast, the representation of each choice is similar between the sexes. Thus, we identify a neural substrate that contributes to sex-specific modulation of motivation by value.

## Introduction

Animals continually select between behaviors based on previous experience with actions and their outcomes. While there is emerging evidence of sex differences in this process of value-based decision-making^1–7^, the neural underpinnings of these sex differences remain largely unknown^8–12^.

Recent studies comparing male and female decision-making suggest that females may be more sensitive to negative outcomes than males. For instance, compared to males, female rats learn to avoid shock faster^4^, are more sensitive to the risk of punishment when seeking reward^2, 13^ and both human and rodent females may be more sensitive to losses in gambling tasks^5, 14, 15^. This may be particularly important given sex differences in susceptibility to psychiatric diseases such as depression and anxiety^16^, diseases which may also relate to altered processing of outcomes^17^.

Our limited knowledge of the neural mechanisms mediating sex differences in decision-making is due to the focus of previous neuroscience studies on male subjects^8–12^. This previous work identified a distributed network of brain regions that contribute to value-based decisions, including the anterior cingulate cortex (ACC) as well as its major striatal target (the dorsomedial striatum, or DMS)^18–35^. These regions may be important not only for the value-dependent selection of actions, but also in regulating motivation to engage in reward-seeking behavior, which is also modulated by value (or expected reward) of the environment or chosen action^29, 36–44^. However, it remains unknown if and how differences in neural activity in these regions are responsible for producing behavioral differences between the sexes.

To address this gap, we trained mice in a self-initiated probabilistic reversal learning task in which they regulated both their trial-by-trial choices and their motivation to engage in the task based on recent experiences^45, 46^. We first asked whether there were differences in behavior between males and females. While choices were similar between the sexes, we found a stronger relationship between value and motivation to engage in the task in females than in males, as assayed with trial initiation latencies^36, 37^. Specifically, females displayed less motivation to engage in the task on lower value trials. We investigated the role of ACC neurons that project to the DMS (ACC-DMS neurons) in this effect, and found that inhibition of these neurons increased motivation to engage in the task and removed the dependence of motivation on value primarily in females. Consistent with these perturbations, we found stronger representation of negative (unrewarded) outcomes, as well as low-value choices, in females compared to males, while downstream connectivity of ACC-DMS neurons was similar between the sexes. Thus, we show that ACC neurons that project to the DMS have a sex-dependent role in the modulation of motivation by value.

## Results

### Value more strongly influences motivation to engage in a decision making task in females than in males

To determine whether and how value-based decisions differ in males and females, we trained mice of both sexes to perform a self-initiated, probabilistic reversal learning task^45, 46^ (**Figure 1a**). After an intertrial interval (ITI), mice initiated a trial by entering a central nose poke, which caused levers to extend on either side. One lever was rewarded with a high probability (0.7) and the other with a low probability (0.1), and after a variable number of trials, there was an unsignaled reversal of the reward probabilities, which occurred multiple times during a session. Mice indicated their choice by pressing one of the levers. If rewarded, a drop of sucrose solution was delivered to a central reward port accompanied by an auditory cue (CS+). Unrewarded outcomes were signaled with a different auditory cue (CS-, see **Methods** for details).

**Figure 1:**
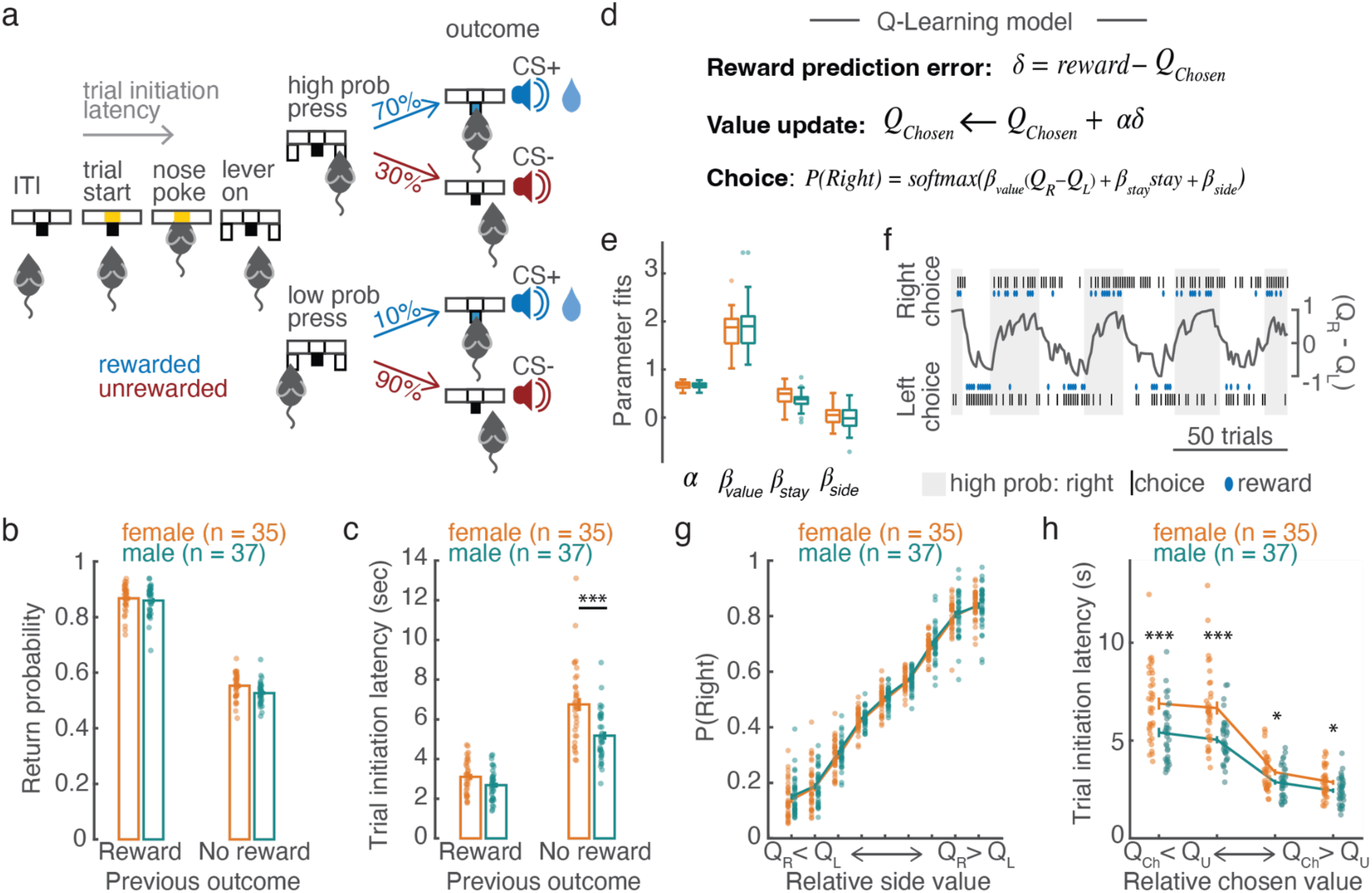
Sex-dependent modulation of motivation, but not choices, by previous outcome and value. **a)** Schematic of the self-initiated, probabilistic reversal learning task. Rewarded outcomes were signaled with a tone (“CS+”) and unrewarded outcomes with a white noise auditory stimulus (“CS-”) **b)** The probability that mice returned to the previously selected lever (return probability) was modulated by previous outcome similarly in males and females (see **Supplementary Table 1** for details of statistical model). **c)** Previous outcome affected trial initiation latencies to a greater extent in females than males (see **Supplementary Table 2** for details of statistical model). Both males and females were significantly slower to initiate trials following rewarded than unrewarded outcomes (Female: F(1,72) = 190.03, p = 6.96x10^-22^; male: F(1,72) = 93.72, p = 1.14x10^-14^, F-tests of contrasts). Additionally, females were significantly slower than males to initiate trials following unrewarded outcomes (male vs. female previously unrewarded: F(1,136.5) = 27.80, p = 5.13x10^-7^, F-tests of contrast) but not following rewarded trials (male vs. female previously rewarded: F(1,136.5) = 1.95, p = 0.16, F-tests of contrast). **d)** Q-learning model used to estimate trial-by-trial action values for the right and left lever. (reward prediction error) is the difference between reward (1 for reward, 0 for no reward) and the action value for the chosen action, *Q_Chosen_*. *Q_Chosen_* is updated with δ multiplied by the learning rate (α). The probability of making a given choice is given by the softmax equation with inverse temperature (*β_value_*), a stay parameter (*β_stay_*), to capture the tendency of mice to repeat the previous action, *stay*, which indicates the previous choice and *β_bias_* to capture side bias. **e)** Box plots of parameter estimates for males and females averaged across sessions. **f)** Example performance of the mouse and model. The shading indicates the lever rewarded with a high probability: gray for the right lever and no shading for the left lever. Black lines indicate the choice made by the mouse (right on top, left on bottom). Rewarded trials are marked with a blue oval, and the hypothesized decision variable, relative side value (*Q_R_* − *Q_L_*, where *Q_R_* and *Q_L_* are the action values for the right and left lever, respectively), is plotted in black. **g)** Choice was significantly and similarly modulated by relative side value in males and females (F-tests on mixed-effects regression: *sex*: F(1,816924) = 1.84, p = 0.18, *relative side value*: F(1,816924) = 311.1, p<0.001, *sex*relative side value*: F(1,816924) = 0.96, p = 0.46; see **Supplementary Table 4** for details of statistical model). For plotting, trials were divided into 9 quantile bins of relative side value and the probability of making a right choice was calculated for each bin for each mouse. **h)** Trial initiation latency was significantly modulated by relative chosen value (*Q_Ch_* − *Q_U_*, where *Q_Ch_* and *Q_U_* are the action values for the chosen and unchosen lever, respectively) to a greater extent in females than males (F-tests on mixed-effects regression: *sex*: F(1,51.84) = 7.82, p = 0.01, *relative chosen value*: F(3,108.79) = 77.03, p = 8.50x10^-27^, *sex:relative chosen value*: F(3,108.78) = 24.70, p = 2.89x10^-12^; see **Supplementary Table 5** for details; post-hoc 2-sided Wilcoxon rank sum tests: male vs. female bin 1: Z = 3.44, p = 5.90x10^-4^; bin 2: Z = 3.98, p = 6.98x10^-5^; bin 3: Z = 2.11, p = 0.04; bin 4: Z = 2.29, p = 0.02). For each mouse, trials were divided into quartiles of relative chosen value and trial initiation latencies were averaged for each bin. **(b,c,e,g,h)** Circles: individual mice; lines or bars averages across mice; error bars are SEM; males (green): n = 37; females (orange): n = 35; *** p < 0.001; * p < 0.05

As expected, mice were significantly more likely to return to the previously selected lever if that choice was rewarded than unrewarded (**Figure 1b**, mixed-effects regression: *previous outcome*: F(1,816592) = 1.38x10^3^, p < 0.001; see **Supplementary Table 1** for details of the statistical model and complete regression results). This effect was similar across the sexes (*previous outcome*sex*: F(1,816592)=0.01, p=0.91, **Figure 1b**).

In contrast to the similarity in return probability between the sexes, motivation to perform the task, which we quantified as trial initiation latency^36, 37^, varied by sex (**Figure 1c**, **Supplementary Figure 1**). While both male and female mice were significantly slower to re-engage in the task following unrewarded than rewarded trials, this effect was stronger in females (**Figure 1c**; mixed-effects regression: *Sex*: F(1,72) = 18.02, p = 6.43x10^-5^; *previous outcome*: F(1,72) = 276.61, p = 2.30x10^-26^; *previous outcome*sex*: F(1,72) = 9.81, p = 0.003, see **Supplementary Table 2** for details of the statistical model). Specifically, females were slower to initiate trials than males following unrewarded, but not rewarded, outcomes (Post-hoc comparisons: male vs. female unrewarded: F(1,136.5) = 27.80, p = 5.13x10^-7^, rewarded: F(1,136.5) = 1.95, p = 0.16).

Because reward is probabilistic in this task, and the reward probabilities associated with each lever change over time, animals integrate choices and outcomes across multiple trials to make a new choice. We modeled this process with a Q-learning model, which was fit hierarchically across animals and sessions, to generate trial-by-trial estimates of the action value (*Q*-values) of selecting each lever (**Figure 1d-f**, **Supplementary Figure 2, Supplementary Table 3**)^47^. The fitted parameters were similar for males and females (**Figure 1e**). As expected, the probability that mice chose the right side lever was significantly modulated by relative side value (*Q_Right_* − *Q_Left_*; **Figure 1f-g**, **Supplementary Table 4**), such that they were more likely to press the lever with the greater *Q*-value. Consistent with the similar fitted parameters for males and females (**Figure 1e**), and the similar effect of previous outcome on return probability in males and females (**Figure 1b**), there was no effect of sex on the relationship between relative side value and choice (**Figure 1g**; mixed-effects regression: *relative side value*sex*: F(8,816924) = 0.96, p = 0.46; see **Figure 1g** and **Supplementary Table 4** for details of statistical model). This suggests that males and females employ similar action selection strategies in this task.

Motivation to perform a behavior is related to the expectation of receiving reward for that behavior^29, 36–41^. In other words, trial initiation latency may be modulated by the value of the chosen action. Indeed, trial initiation latency was significantly modulated by relative chosen value (*Q_Chosen_* − *Q_Unchosen_*), such that mice were slower to initiate low relative chosen value trials (**Figure 1h**, mixed-effects regression: *relative chosen value*: F(3,108.8) = 77.03, p = 8.50x10^-27^). Furthermore, this effect was significantly modulated by sex (*relative chosen value*sex*: F(3,108.8) = 24.70, p = 2.89x10^-12^; see **Figure 1h** and **Supplementary Table 5** for details of statistical model). In particular, when females selected the lower value action, they slowed their trial initiation latency to a greater extent than males (**Figure 1h**), consistent with their greater modulation of trial initiation by previous outcome (**Figure 1c**).

Motivation is also modulated by the estimated total value of available options (*Q_Chosen_* + *Q_Unchosen_*, **Supplementary Fig. 3a**)^29, 36, 39, 40, 48^, and this modulation was also sex-dependent: mice were slower to initiate trials when the total value was low, and females were significantly slower than males on low total value trials (**Supplementary Fig 3a**). Furthermore, both total and relative chosen value significantly and sex-dependently predicted trial initiation latencies when included in the same model (**Supplementary Table 6** for details).

To assess whether fluctuations in the levels of circulating hormones could contribute to these behavioral differences, we monitored estrous cycle in a separate group of female mice and analyzed how estrous stage affected value-dependent modulation of motivation (**Supplementary Fig. 4**). There was a small effect of estrous cycle on the relationship between relative chosen value and trial initiation latency (**Supplementary Fig. 4b**). Specifically, mice were significantly slower on low relative chosen value trials during the follicular phase of the estrous cycle (proestrus and estrus when estradiol and progesterone levels are high) compared to the luteal phase (metestrus and diestrus)^49, 50^.

Task engagement was also modulated by daily fluctuations in weight (possibly related to overall levels of motivation due to variation in thirst and/or hunger across days, **Supplementary Figure 1e-f**), as well as by time in session (possibly related to satiety, **Supplementary Figure 1g**). However, the relationships between weight or time in session (trial number) and value-dependent modulation of trial initiation latency were not significantly modulated by sex, suggesting that weight and satiety affect value-dependent motivation similarly in males and females (*weight*relative chosen value*sex:* F(3,99.67) = 2.56, p = 0.06; *trial*relative chosen value*sex:* F(3,86.88) = 1.40, p = 0.25; see **Supplementary Table 5** for details of statistical model).

We next asked whether this sex difference in motivation to engage in the task was specific to mice, or if there were also gender differences in human subjects performing a similar task (142 female, 208 male, age 19-70; task schematized in **Supplementary Fig. 5a**, see **Methods** for details). We fit the hierarchical *Q* -learning model described in **Figure 1d-f** to evaluate how choice and trial initiation latency were modulated by value in men and women. Similar to the mice, parameter estimates from this model were similar in men and women (**Supplementary Fig 5b**) and there were no gender differences in the influence of relative side value on choice (mixed-effects regression: *gender*: F(1,142619) = 0.93, r*elative side value:* F(1,142619) = 99.9, p = 1.64x10^-207^, *relative side value*gender*: F(1,142619) = 0.81, p = 0.62; see **Supplementary Fig 5c,d** for details of the statistical model). On the other hand, there was a small but significant gender by value interaction (mixed-effects regression: *gender*relative chosen value*: F(4,1750) = 2.50, p = 0.04). Interestingly, this effect was modulated by age (**Supplementary Fig. 5e-g;** *relative chosen value*gender*age*: F(4,792) = 2.45, p = 0.04, see **Supplementary Fig. 5** for details of the statistical model). In particular, older men were slower to initiate trials and showed increased value modulation compared to younger men (**Supplementary Fig. 5e-g**). Thus, the behavior of younger adults was similar to the mice, in that females were slower than males to initiate low-value trials (**Supplementary Fig. 5f**).

### Inhibition of ACC-DMS neurons during outcome presentation removes the influence of relative chosen value on motivation in females

Given previous work implicating the ACC and its striatal target, the DMS, in value-dependent decision-making and motivation^18–28, 36, 51^, we examined whether this circuit could contribute to the observed sex differences in behavior. ACC-DMS neurons were targeted for inhibition by injecting a retroAAV into the DMS to express *Cre*-recombinase, and a *Cre-* dependent virus in the ACC to express the inhibitory opsin eNpHR3.0 (or control virus) in ACC-DMS neurons. On a small subset of trials, we bilaterally delivered 2 seconds of 5 mW light, triggered by the onset of the CS signaling choice outcome, because this period is likely when mice decide when to re-engage in the task (**Figure 2a-b**, **Supplementary Fig. 6**, see **Methods** for details).

**Figure 2:**
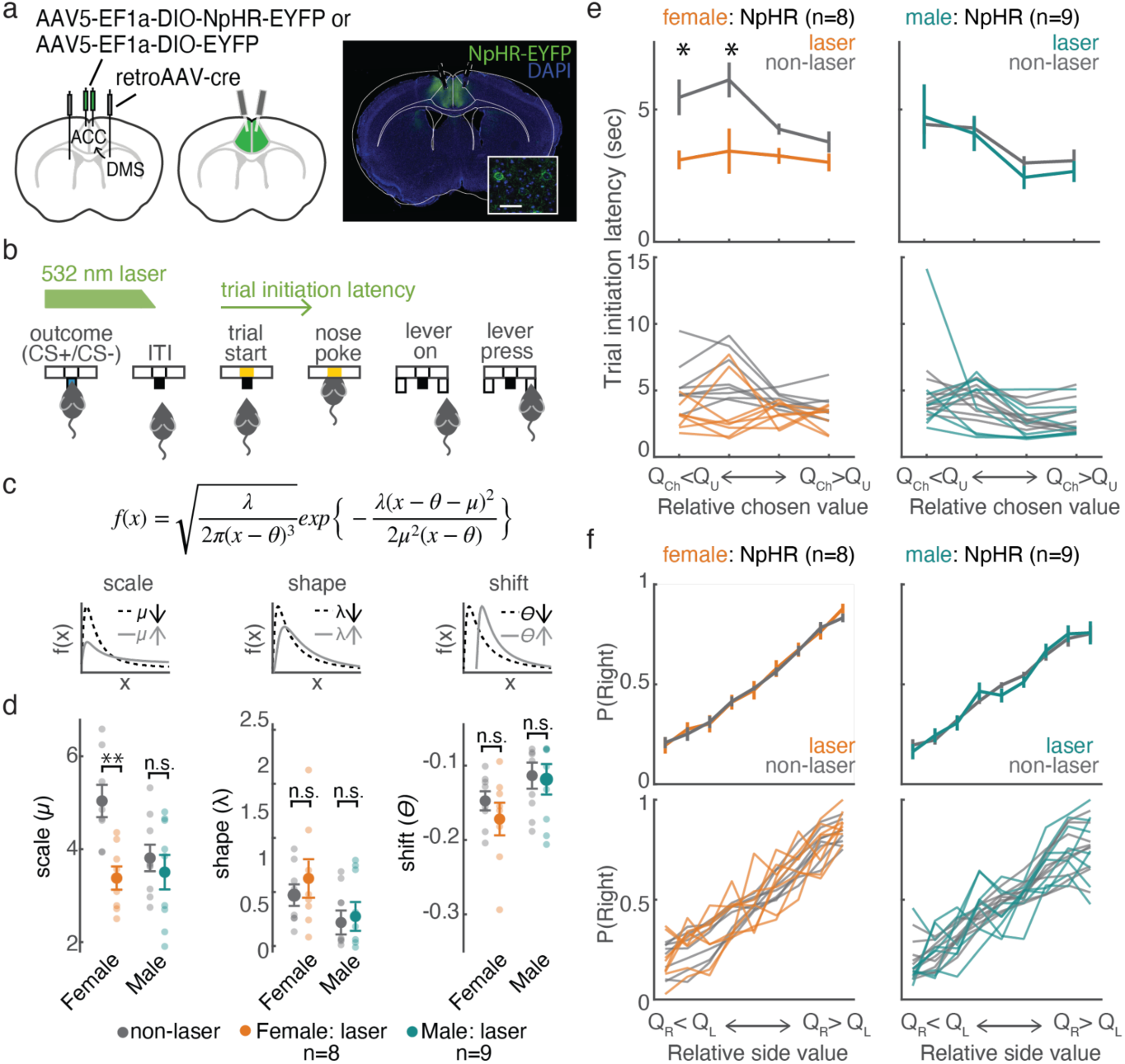
ACC-DMS inhibition increased motivation in female mice, especially on low relative chosen value trials, without affecting choice in either sex. **a.** Left: schematic of viral strategy to target ACC neurons projecting to the DMS. Right: example histology showing NpHR-EYFP expression and the lesions from optic fibers. Inset: confocal image of neurons expressing NpHR-EYFP. Scale bar = 50 *μ*m. **b.** On a subset of trials (5-7.5%), 532 nm light was delivered bilaterally to the ACC for 2 seconds, triggered by the presentation of the outcome (CS+ or CS-). Laser power was gradually ramped down during the last 200 ms. Effects of inhibition were assessed by comparing trial initiation latencies preceded by inhibition (laser trials, indicated by green arrow) and not preceded by inhibition (non-laser trials). **c.** Each mouse’s trial initiation latencies were fitted with a shifted inverse gaussian distribution separately for laser and non-laser trials. Top: the shifted inverse gaussian distribution. Bottom: illustration of the influence of each parameter on the distribution. **d.** Laser significantly affected trial initiation latencies only in females. The scale parameter (μ) was significantly lower for laser trials compared to non-laser trials in females expressing NpHR (W = 36, *p* = 0.008, 2-sided Wilcoxon signed rank test). There were no differences between estimates for the shape or shift parameter for laser and non-laser trials, and no parameters were affected in the males (2-sided Wilcoxon signed rank tests, p>0.1). Transparent circles: individual mice; opaque circles: average across mice; error bars: SEM. **e.** ACC-DMS inhibition affected trial initiation latencies in females expressing NpHR on low relative chosen value trials (**Supplementary Table 7** for details of mixed-effects regression; post-hoc comparisons between laser and control trials for males and females: female laser vs. no laser, value bin 1: W = 35, p = 0.02; value bin 2: Z = 32, p = 0.04, value bin 3: W = 32, p = 0.05; value bin 4: W = 31, p = 0.08; n = 8; male laser vs. no laser: value bin 1: W = 28, p = 0.57; value bin 2: W = 24, p = 0.91, value bin 3: W = 36, p = 0.13; value bin 4: W = 34, p = 0.20, 2-sided Wilcoxon signed rank tests). **f.** ACC-DMS inhibition had no effect on choice (see **Supplementary Table 9** for details of mixed-effects regression). (**e-f**) Average and standard error of the mean across mice (top) and data from individual mice (bottom). n = 8 females, 9 males; *Q_Ch_*: action value for the chosen lever; *Q_U_*: action value for the unchosen lever; *Q_R_*: action value for the right lever; *Q_L_*: action value for the left lever; * p < 0.05, ** p < 0.01.

Interestingly, ACC-DMS inhibition during the outcome of the previous trial significantly affected trial initiation latencies (our proxy for motivation) in females (**Figure 2c-e**). To quantify this, we fit each animal’s trial initiation latencies from trials preceded by outcome inhibition (laser trials) and control (non-laser) trials with a shifted inverse gaussian distribution (**Figure 2c-d** and **Supplementary Fig. 7a**). The scale parameter (*μ*), which reflects the density of trials in the peak versus the tail of the distribution, was significantly lower for laser trials than control trials in females expressing NpHR (2-sided Wilcoxon signed rank test, W = 36, p = 0.008; **Figure 2d**). This indicates that trial initiation latencies were faster and task engagement was higher following inhibition in female mice. The other parameters (shape, λ, and shift, θ) were not significantly affected by inhibition (2-sided Wilcoxon signed rank test, p > 0.1). Furthermore, there were no significant differences between parameters fit to laser and control trials in males expressing NpHR or in males or females expressing EYFP (**Figure 2d**, **Supplementary Figure 7a**).

We also tested whether inhibition during other trial epochs affected trial initiation latency distributions. There were no significant effects of inhibition on the estimated parameters in these conditions (**Supplementary Figure 8**). Furthermore, we did not observe a correlation between the effect of inhibition and individual sensitivity to relative chosen value during sessions without inhibition (**Supplementary Figure 9a**).

Given that relative chosen value more strongly influences motivation in females than males (**Figure 1h**), we wondered whether there was a sex-dependent effect of ACC-DMS inhibition during the outcome on the relationship between relative chosen value and trial initiation latency (**Figure 2e** and **Supplementary Figure 7b**). The effect of laser depended on the interaction between opsin and sex (**Figure 2e**; mixed effects regression on the difference in trial initiation latency following laser and control trials with opsin, sex, relative chosen value and their interactions as fixed effects and random effects of subject; *opsin*sex:* F(1,100) = 4.24 p = 0.04, see **Supplementary Table 7** for details). Specifically, ACC-DMS inactivation reduced the influence of value on trial initiation latency in females expressing NpHR when relative chosen value was low (2-sided Wilcoxon signed rank test, laser v. no laser, value bin 1: W = 35, p = 0.02; value bin 2: W = 33, p = 0.04, value bin 3: W = 32, p = 0.05; value bin 4: W = 31, p = 0.08; n = 8; **Figure 2e**; **Supplementary Figure 7b**). This effect was similar early and late in the session (**Supplementary Figure 10** and **Supplementary Table 8**). Inhibition also similarly disrupted the relationship between total value and motivation to engage in the task sex-dependently (**Supplementary Figure 3b**).

In contrast, inhibition of ACC-DMS neurons during the outcome had no effect on choice in the next trial in males or females (**Figure 2f**, **Supplementary Figure 7c**, see **Supplementary Table 9** for details of statistical model). Thus, inhibition of ACC-DMS neurons preferentially reduced the influence of relative chosen value on motivation in females, without affecting upcoming choice in either sex.

### Similar excitatory synaptic strength of ACC inputs on DMS D1 and D2 MSNs in males and females

Why might ACC-DMS inhibition differentially affect motivation in females versus males? One possibility is differences in the connection strength between ACC-DMS neurons and the direct and indirect pathways of the DMS (D1R versus D2R-expressing neurons), as these pathways are themselves thought to differentially modulate motivation^44, 52–58^. However, we found that activation of ACC-DMS terminals evoked similar EPSCs in D1R and D2R neurons in both sexes (**Figure 3d**; mixed effects regression: *MSN subtype:* F(1,25) = 0.940, p = 0.342; *Sex*: F(1,8.497) = 0.062; *MSN subtype*Sex:* F(1,25) = 2.442, p = 0.131; see **Supplementary Table 10** for details of statistical model). Furthermore, the amplitude of optogenetically-evoked excitatory postsynaptic potentials did not differ in D1R and D2R MSNs in males (**Supplementary Fig. 11**). While we cannot make definitive conclusions from a negative result, this suggests that differences in neural activity in ACC-DMS neurons, rather than differences in downstream connectivity, may mediate the observed sex differences in the effect of optogenetic inhibition.

### More ACC-DMS neurons signal negative outcomes in females than in males

We thus evaluated whether there were sex differences in neural activity in ACC-DMS neurons. We expressed the calcium indicator GCaMP6f in ACC-DMS neurons (**Figure 4a**), and imaged GCaMP6f fluorescence in behaving mice using a head-mounted microscope through a gradient refractive index (GRIN) lens or prism implanted in the ACC^59^. An example field of view and regions of interest extracted with CNMFe^60^ are shown in **Figure 4b** (implant locations in **Supplementary Figure 12**) and the trial-averaged, event-triggered fluorescence for all imaged neurons is shown in **Supplementary Figure 13a**.

**Figure 3:**
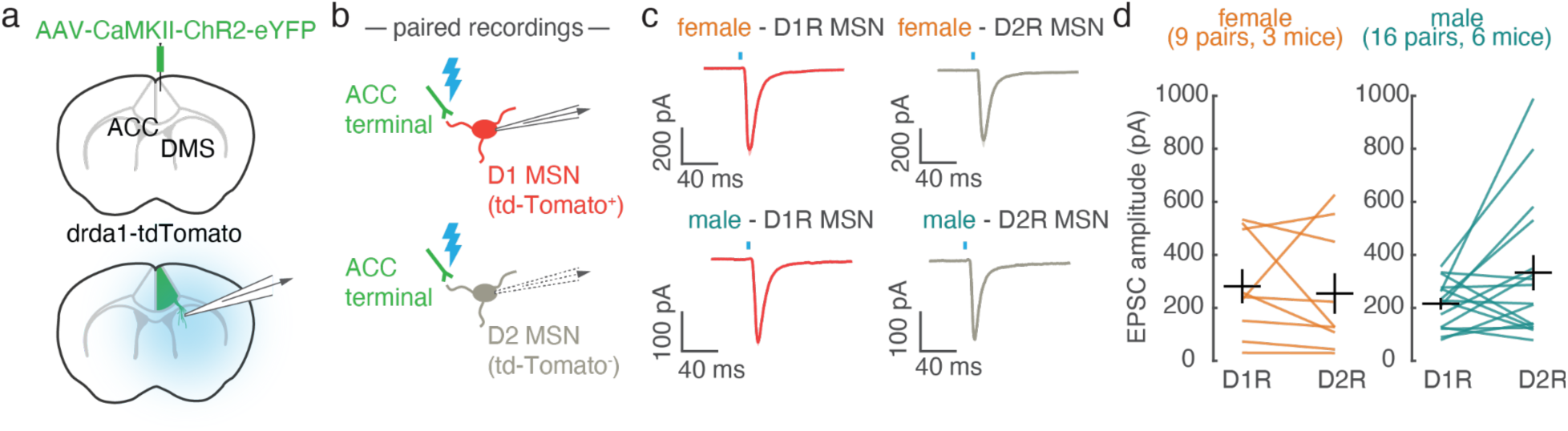
Similar excitatory synaptic strength of ACC-DMS neurons on D1R versus D2R MSNs in males and females. **a.** Schematic of viral strategy to express ChR2 in ACC neurons (top), and record postsynaptic optogenetically-evoked excitatory postsynaptic currents (EPSCs) in DMS (bottom). **b.** Schematic of paired, sequential recordings of neighboring MSNs, where MSNs were visually identified as D1R MSNs (tdTomato+) or D2R MSNs (tdTomato-). Brief light pulses elicited EPSCs from ChR2-expressing ACC terminals. **c.** Example EPSCs measured in pairs including D1R MSN (red) and D2R MSN (grey) in brain slices taken from female (top row) or male (bottom row) mice. Traces are mean responses across trials from a single cell. Shading is SEM. Blue line indicates the time of light stimulation. **d.** Summary of EPSC amplitudes in MSN pairs from female (left) and male (right) mice. A linear mixed effects regression on EPSC amplitudes showed no effect of MSN subtype, sex, or their interaction (*MSN subtype*: F(1,25) = 0.940, p = 0.342; *Sex*: F(1,8.497) = 0.062, p = 0.809; *MSN subtype*Sex:* F(1,25) = 2.442, p = 0.131; see **Supplementary Table 10** for details of mixed-effects regression).

**Figure 4:**
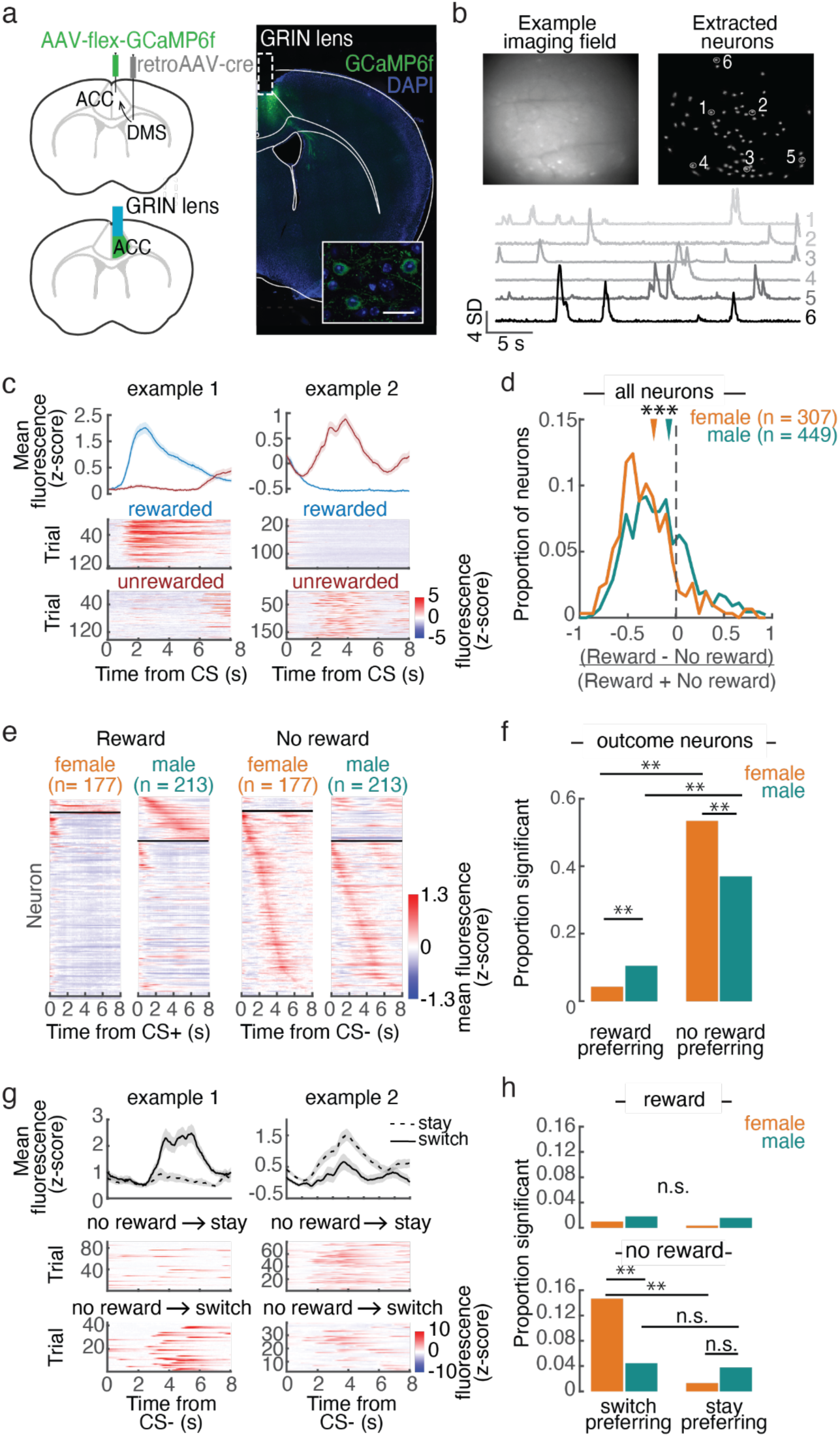
More ACC-DMS neurons respond during unrewarded outcomes in female mice compared to males. **a.** Left: schematic of the strategy to express GCaMP6f specifically in ACC-DMS neurons. Right: example histological section with the location of the GRIN lens in the ACC and expression of GCaMP6f. Inset: confocal image of expression in individual ACC-DMS neurons. Scale bar is 50 *μ*m. **b.** Mean projection of a subset of frames from an example imaging session (top left) and the location of regions of interest extracted using CNMFe^60^ (top right). Example fluorescence traces from the circled neurons (bottom). **c.** Example outcome responses of 2 ACC-DMS neurons. Trial-averaged, Z-scored fluorescence time-locked to the CS+ (red) and CS- (blue) presentation (top). The heatmaps show the CS-triggered activity for all rewarded (middle) and unrewarded trials (bottom). **d.** Distributions of an outcome-modulation index for ACC-DMS neurons (fluorescence was averaged between 0 and 8 seconds following CS presentation and the outcome modulation index was calculated as (reward fluorescence - no reward fluorescence)/(reward fluorescence + noreward fluorescence). The outcome-modulation index was significantly more negative (i.e. biased towards negative outcomes) in female than male neurons (2-sided, Wilcoxon rank sum test, Z = 5.22, p = 1.78x10^-7^). Arrows: median of each distribution. **e.** Trial - averaged, Z - scored fluorescence time-locked to CS presentation for all neurons that significantly encoded outcome (see **Methods** for details). Outcome-encoding neurons were classified as reward-preferring (above the black line) or no reward-preferring (below the black line; see **Methods** and **Supplementary Figure 8B** for details), and each subset of neurons were then separately sorted by the time of their peak fluorescence. **f.** Femaleshad more outcome-encoding and more no reward-preferring neurons than males, and males had more reward-preferring neurons than females (*χ* ^2^−test for the proportion of outcome-encoding neurons in males and females, *χ* ^2^(2, n=756) = 24.16, p = 5.67x10^-6^; paired comparisons performed with the Marascuilo procedure with alpha = 0.01). In females, 177/307 (57.7%) of neurons significantly encoded outcome and 13 of these (7.3%) were preferentially active on rewarded trials and 164 (92.7%) were preferentially active on unrewarded trials. In males, 213/449 (47.4%) of neurons significantly encoded outcome and 47 of these (22.1%) were reward-preferring and 166 (77.9%) were no reward-preferring. **g.** Example neurons that were preferentially active during unrewarded outcomes preceding trials when the mouse was going to switch to the alternative lever (left) or return the previously selected lever (right) on the next trial. Top panels: trial-averaged fluorescence. Bottom panels: fluorescence during unrewarded outcomes for all upcoming stay trials (top) or switch trials (bottom). **h.** Top: no difference in the proportion of male and female neurons significantly encoding stay versus switch during rewarded outcomes (*χ*^2^−test, *χ* ^2^(2,n=756) = 3.5, p = 0.17). Bottom: significantly more neurons encoded stay versus switch in females than males during unrewarded outcomes (*χ* ^2^−test, *χ* ^2^(2,n=756) = 27.35, p = 1.1x10^-6^). There was no sex difference in the proportion of stay-preferrring neurons (see **Methods** and **Supplementary Figure 13d** for classification of neurons), but females had significantly more switch-preferring neurons than males. Additionally, females had more switch-preferring than stay-preferring neurons, while in males there was no difference (Marascuilo procedure with alpha = 0.01 for paired comparisons). (**c,g**) shading is SEM.

**Figure 5:**
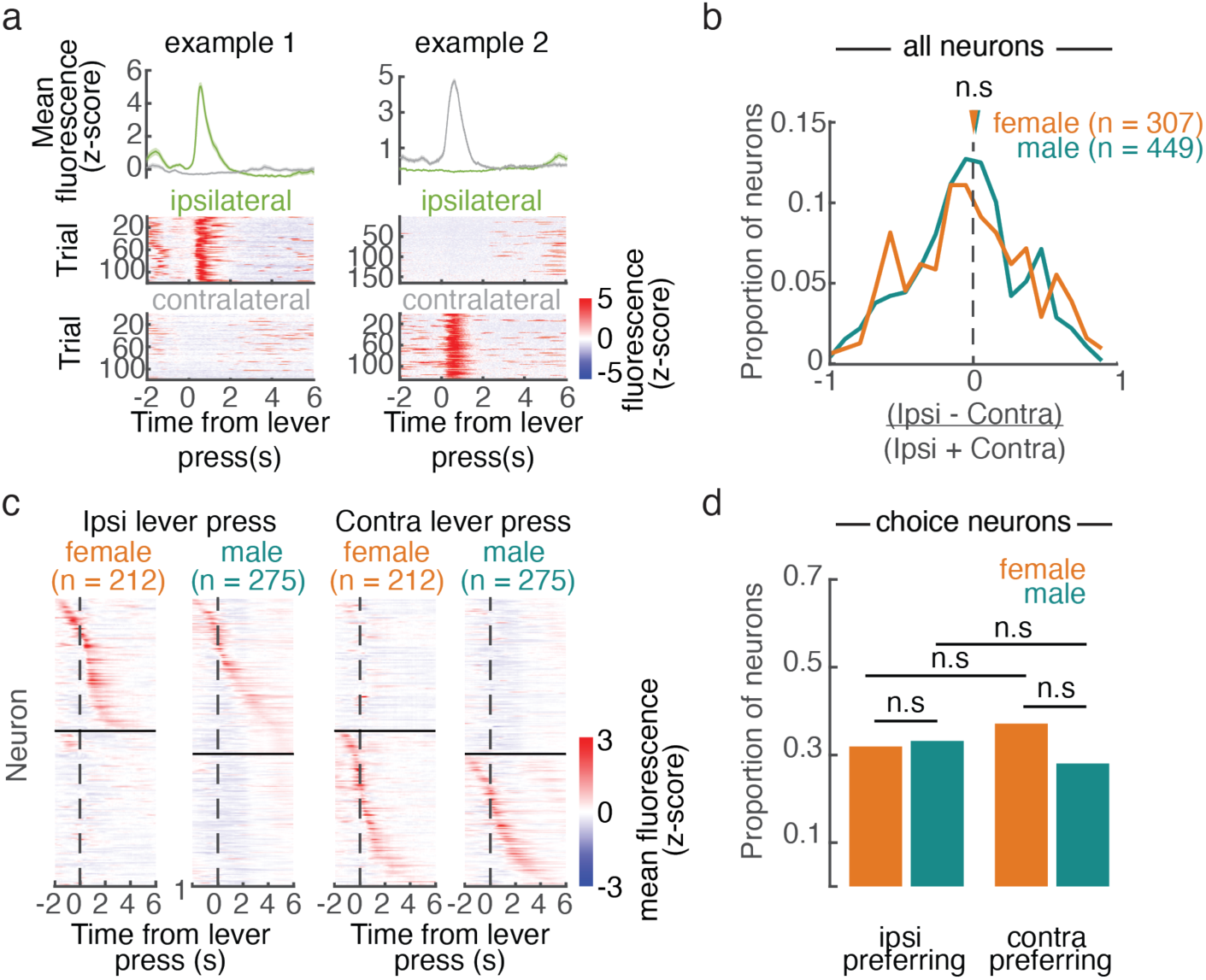
Similar correlates of contralateral versus ipsilateral actions in ACC-DMS neurons of females and males. **a.** Example neuron preferentially active during ipsilateral (left column) or contralateral (right column) trials relative to the imaged side. Top panels: trial-averaged fluorescence traces. Bottom panels: all ipsilateral trials (top) and contralateral trials (bottom). Shading is SEM. **b.** Distribution of choice modulationindex (fluorescence was averaged from 2 seconds before until 6 seconds after the lever press and the choice modulation index was calculated as (ipsilateral fluorescence contralateral fluorescence)/ (ipsilateral fluorescence + contralateral fluorescence). The choice modulation indices were not different between males and females (2-sided, Wilcoxon rank sum test, Z=1.10, p = 0.27). **c.** Trial-averaged Z-scored fluorescence for ipsilateral and contralateral lever press events for neurons significantly modulated by choice (212/307, 69.1% in females and 275/449, 61.25%). Neurons above the black line were classified as ipsilateral choice-preferring and those below the black line were contralateral choice-preferring (see **Supplementary Figure 13c** and **Methods** for details of classification). **d.** Proportion of significant ipsi- and contra-preferring neurons in males and females. There were no differences in the proportion of ipsilateral- and contralateral preferring neurons within or between the sexes (*χ* ^2^-test, *χ* ^2^ (2, n=756) = 7.94, p = 0.02, Marascuilo procedure with alpha = 0.05).

Because males and females differed in how previous outcome modulated motivation to engage in the task (**Figure 1c**) and inhibition of ACC-DMS neurons during the outcome sex-dependently affected motivation (**Figure 2**), we first investigated sex differences in the neural correlates of outcome in ACC-DMS neurons (example outcome responses in **Figure 4c**). In both males and females, ACC-DMS neurons were more active following unrewarded outcomes than rewarded outcomes (**Figure 4d**)^61–63^. Interestingly, this bias towards negative outcome encoding was more pronounced in females compared to males (2-sided, Wilcoxon rank sum test comparing all imaged male and female neurons, Z = 5.22, p = 1.78x10^-7^, n = 307 female neurons, n = 449 male neurons; **Figure 4d**).

Similar results emerged from considering only the neurons with significantly different activity on rewarded and unrewarded trials (**Figure 4e-f**). We classified neurons as outcome-encoding based on whether including the type of outcome (rewarded vs. unrewarded) improved the fit of a linear encoding model for each neuron, compared to a null distribution generated from fitting the same models to randomly shifted fluorescence data (see **Methods** for details)^64, 65^. In females compared to males, a significantly larger proportion of outcome-encoding neurons were preferentially active on unrewarded trials, while a significantly smaller proportion were preferentially active on rewarded trials (**Figure 4e-f**; *χ* ^2^−test comparing the proportions of outcome-encoding neurons in males and females, *χ* ^2^ (2, n=756) = 24.16, p = 5.67x10^-6^; the Marascuilo procedure with alpha=0.01 was used for comparisons between proportions of reward- and no reward-preferring outcome-encoding neurons in females and males; see **Supplementary Figure 13b** for definition of reward- and no reward-preferring neurons). We did not observe a relationship between individual variability in value modulation of trial initiation latencies and bias towards negative outcome encoding (**Supplementary Figure 9b**). Thus, following unrewarded outcomes, when females are slower to re-engage with the task than males (**Figure 1c**), more of the ACC-DMS population was active in females than in males. This, together with the inhibition results (**Figure 2**), suggests that increased activation of the ACC-DMS population during negative outcomes may decrease motivation to perform the task.

Previous studies report that activity in the ACC is correlated with changes in strategy (e.g. switching to the alternate choice)^66–68^. Therefore, we wondered whether outcome responses differed based on whether the mouse would return to the previously selected lever or switch to the alternative lever on the next trial (**Figure 4g-h**). Interestingly, during unrewarded outcomes, in females, but not in males, more neurons were preferentially active preceding switch trials, and females had significantly more switch-preferring neurons than males (**Figure 4h**, *χ* ^2^− test, *χ* ^2^ (2,n=756) = 27.35, p = 1.1x10^-6^, comparing the proportion of stay/switch neurons in males and females on unrewarded trials; Marascuilo procedure with alpha = 0.01 for post-hoc comparisons, see **Supplementary 13d** for definition of stay- and switch-preferring neurons). Thus, not only did females have more neurons encoding negative outcome (**Figure 4d,f**), but negative-outcome activity was modulated by upcoming strategy to a greater extent than in males. This increased activity preceding switch trials may relate to females’ greater modulation of trial initiation latency dependent on upcoming stay versus switch compared to males (**Supplementary Fig 1h**).

### Choice is similarly represented in male and female ACC-DMS neurons

In addition to outcome encoding, choice encoding was also prominent in the ACC-DMS population (example neurons active during a particular choice in **Figure 5a**). However, in contrast to outcome, the activity of ACC-DMS neurons was similarly selective for ipsilateral versus contralateral lever presses in males and females (**Figure 5b,** 2-sided, Wilcoxon rank sum test, Z=1.10, p = 0.27, n = 307 female neurons, n = 449 male neurons). Consistently, when considering only significant choice-encoding neurons (see **Methods** and **Supplementary Figure 12c** for details), there were no sex differences in the proportion of either ipsi- or contralateral preferring neurons, nor differences within the sexes (**Figure 5D**; *χ* ^2^− test, *χ* ^2^ (2,n=756) = 7.9, p = 0.02; no comparisons significant based on Marascuilo procedure with alpha = 0.05).

Thus, we found evidence that in the ACC-DMS population, neural correlates of outcome, but not choice, significantly differ by sex, such that these neurons signal negative outcomes to a greater extent in females than in males. Interestingly, this parallels our behavioral findings that choice did not differ by sex (**Figure 1b,g**), as well as our optogenetic findings that inhibition of this projection affected trial initiation latencies (motivation), but not choice (**Figure 2**).

### Relative chosen value, but not relative side value, is differentially represented in female and male ACC-DMS neurons

Given the preferential encoding of negative outcomes in females (**Figure 4**), and the value-dependent effect of ACC-DMS inhibition during the outcome period on motivation (**Figure 2e**), we may expect related sex differences in value representations during the outcome period. Indeed, we found that during the outcome period, chosen and total value encoding was more prominent in females than males (**Figure 6** and **Supplementary Figure 3c**). Significant value-encoding neurons were identified with bilinear encoding models fit to each neuron’s fluorescence, which allows transient responses related to task events to be modulated multiplicatively by value. Significance was assessed relative to a null distribution produced by randomly shifting the neural data (**Supplementary Figure 14a-c**, see **Methods** for details).

**Figure 6:**
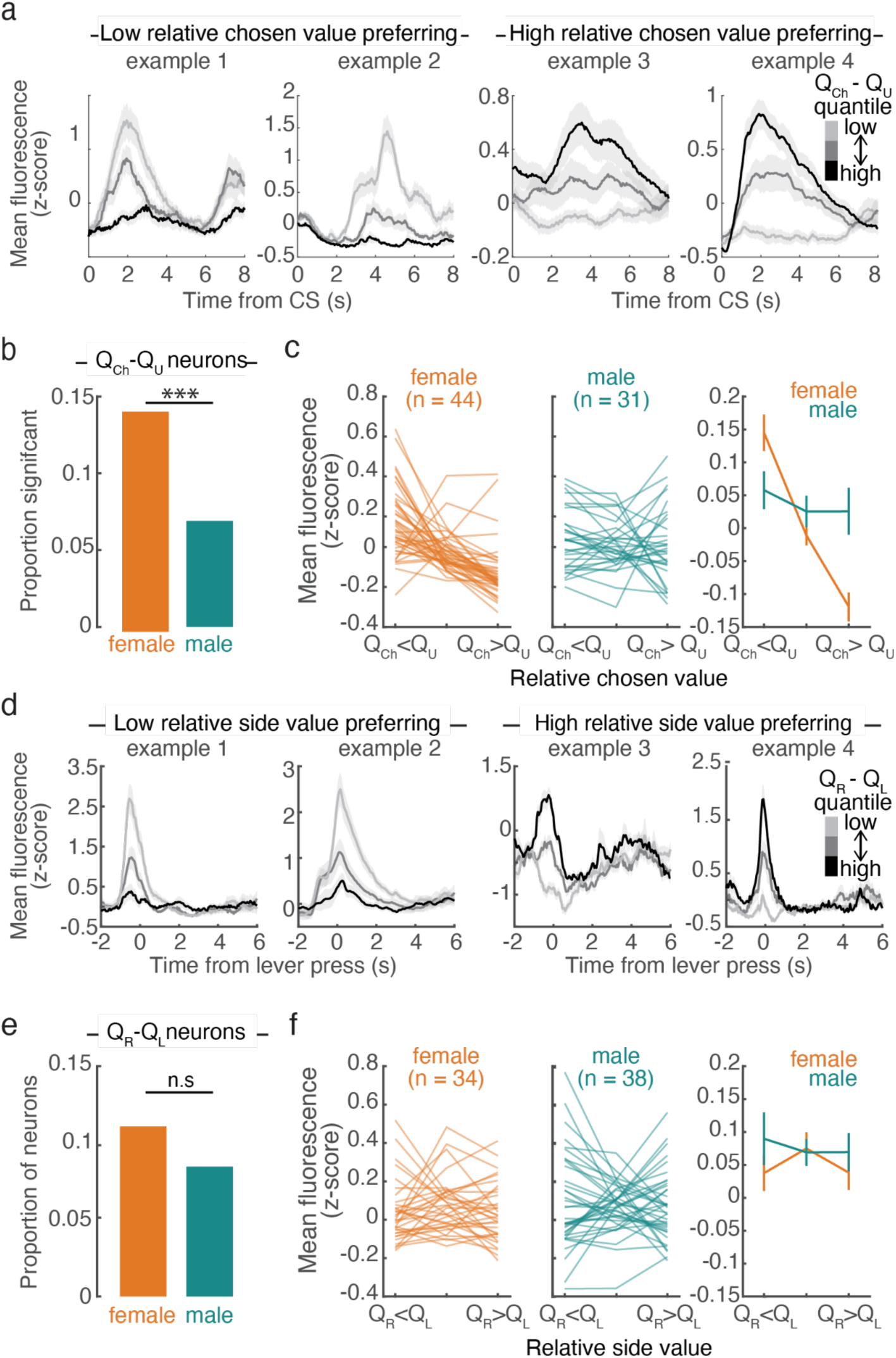
More relative chosen value encoding, but similar relative side value encoding, in females compared to males. **a.** Example fluorescence time-locked to the CS, plotted by quantile bin of relative chosen value (*Q_Ch_* − *Q_U_* where *Q_Ch_* and *Q_U_* are the action values for the chosen and unchosenlever, respectively). Examples 1 and 2 were more active during the outcome period preceding low relative chosen value trials and examples 3 and 4 were more active preceding high relative chosen value trials. **b.** Femaleshada significantly larger proportion of neurons encoding relative chosen value during the outcome than males (*χ* ^2^-test, *χ* ^2^ (1, n=756) = 11.26, p=0.0008). **c.** Average fluorescence during the outcome period versus relative chosen value quantile bin for n e u r o n s s i g n i f i c a n t l y encoding relative chosen value during the outcome for females (left) and males (center). Mean and SEM across significant neurons (right, n = 44/307 female neurons, n = 31/449 male neurons). **d.** Example relative side value (*Q_R_* − *Q_L_*, where *Q_R_* and *Q_L_* are the action values for the right and left lever, respectively) encoding neurons. Mean lever-press triggered fluorescence by relative side value (quantile bins). Examples 1 and 2 were preferentially active during low relative side value lever presses and examples 3 and 4 were preferentially active during high relative side value lever presses. **e.** There was no significant difference in the proportion of neurons significantly encoding relative side value in males versus females (*χ* ^2^-test, *χ* ^2^(1, n=756) = 1.44, p = 0.23). **F.** Average lever press activity versus relative side value quantile bin for neurons significantly encoding relative side value during the lever press for females (left) and males (center). Mean and SEM across significant neurons (right, n = 34/307 female neurons, n = 38/449 male neurons).

A higher fraction of ACC-DMS neurons in females than males significantly encoded relative chosen value during the outcome (**Figure 6a-c**, 14.3% in females (44/307) and 6.9% in males (31/449); *χ* ^2^ -test, *χ* ^2^ (1, n=756) = 11.26, p=0.0008; **Supplementary Figure 14d**). In females, these neurons were most active preceding low relative chosen value trials (**Figure 6c**), which is consistent with our observation of more negative outcome encoding in females (**Figure 4**), and the greatest effect of inhibition of these neurons on low chosen value trials (**Figure 2e**). Similarly, the proportion of total value-encoding neurons was significantly larger in females than males during the outcome period (*χ* ^2^-test, *χ* ^2^(1,N=756) = 13.58, p = 2.3x10^-4^; **Supplementary Figure 3c**).

We next asked whether ACC-DMS neurons encoded relative side value, which is predictive of choice (**Figure 1g**). A small proportion of neurons (8.5% in males (38/449) and 11.1% in females (34/307)) had lever press-related activity that was significantly modulated by relative side value. Consistent with the similar choice encoding in males and females (**Figure 5**), this proportion was not significantly different between males and females (*χ* ^2^-test, *χ* ^2^(1, n=756) = 1.44, p = 0.23; **Figure 6e-f** and **Supplementary Figure 14e**).

## Discussion

While males and females make similar choices in a value-based decision-making task, we found differences in how action values relate to motivation to engage in the task (**Figure 1**). In addition, we identified a sex-dependent role for the projection from the ACC to the DMS in this value-dependent regulation of motivation. Specifically, females were less motivated than males to engage in the task when relative chosen value was low (**Figure 1**). Inactivation of ACC-DMS neurons during the outcome led to an increase in motivation on low relative chosen value trials in females, without an effect on choice in either sex (**Figure 2**). Consistent with these behavioral results, we found increased representation of unrewarded outcomes and low relative chosen value in ACC-DMS neurons in female mice compared to males (**Figure 4, 6a-c**), without sex differences in the representation of choice (**Figure 5, 6d-f**). Thus, we identified value-dependent differences in motivation but not choices between the sexes, as well as a neural substrate that contributes to these sex differences in motivation without affecting choices.

### Previous behavioral evidence for sex differences in value-based decision-making

Although neuroscience research has focused primarily on male subjects^8,9,11, 12^, recent studies have begun to explore sex differences in cognitive behaviors, including value-based decision making^1–6, 69–72^. These studies find that while males and females perform decision-making tasks with similar degrees of accuracy, they can employ different strategies to learn or execute these tasks^1,3,6, 15, 69^. Our results are consistent with this idea. Males and females make similar choices and achieve similar rates of reward per trial when performing the reversal learning task, but differ in how recent experience affects their motivation to perform the task. Specifically, females are more likely to disengage from the task when they are less likely to be rewarded. This may reflect a greater tendency to explore alternative behaviors when engaging in the task is less profitable. Compatible with this idea, the ACC is thought to be important for regulating exploratory behavior^68, 73–75^. Furthermore, sex differences in exploration have been reported in a reinforcement learning task^7^ and exploratory choices in a reversal learning task have been linked to the effect of sex hormones on D2R MSNs in the DMS in females^76^.

Another consistent sex difference in cognitive behavior is the finding that females are more sensitive to negative or aversive outcomes than males^2, 4, 5, 13–15, 71, 77^. Even though there is no explicit punishment in the present study, our results are consistent with the interpretation that females are more sensitive to negative outcomes (in our case, unrewarded trials). In particular, females are less motivated than males following unrewarded but not rewarded outcomes and are more sensitive to low-value trials, when their actions are unlikely to lead to reward (**Figure 1c,h**).

### New insight into the neural basis of sex difference in value-based decision-making

Despite recent interest in the study of sex differences in cognitive behavior, the neural mechanisms underlying these differences remain largely unknown. While there is evidence for structural and functional differences in various brain regions of males and females, including the ACC and striatum^6, 70, 78–83^, these studies were conducted without behavior or used methods that were unsuited to temporally resolve neural correlates of behavior.

Thus, the novelty of our study is linking activity of a projection-defined neural population in males and females to sex differences in behavior. This allowed us to discover sex differences in neural correlates of the task, as well as to identify a causal and sex-dependent role of the ACC projection to the DMS in motivation but not choice. Our findings may relate to a study of impulse inhibition in humans, which showed that despite similar accuracy in a go/go-no task, women had slower reaction times than men and differing activation patterns in the ACC^84^.

Together our results suggest that increased activation of ACC-DMS neurons promotes disengagement from reward-seeking behavior when the probability of reward is low. The greater endogenous activity in this projection in females following unrewarded outcomes (**Figure 4**) and low value trials (**Figure 6a-c**) may explain why the effect of inhibiting these neurons was larger in females than in males (**Figure 2**). An important open question is whether a different population of neurons mediates this behavior in males, or if in different behavioral contexts males display greater value-dependent modulation of motivation and greater activation of ACC-DMS neurons. Another possibility is that ACC-DMS neurons contribute to value-dependent modulation of motivation in both males and females during the reversal learning task, but that other neural circuits compensate for the effects of ACC-DMS inhibition more effectively in males. Another open question is what mechanisms underlie the sex differences in the role of ACC-DMS neurons in the regulation of motivation. There are numerous ways that sex influences neural function and behavior. There are widespread genetic differences between males and females as well as effects of gonadal hormones during development or puberty^10, 85, 86^. Additionally, sex could indirectly lead to differences because males and females interact differently with their environments throughout their lives^86, 87^. Differences can also depend on circulating hormones in adulthood. Although several studies found no effect of estrous cycle on value-based choices in females^15, 71^, manipulating gonadal hormones does affect decision making in males^88^ and females^89, 90^. Here we found a small effect of the estrous cycle on motivation (**Supplementary Figure 3**), consistent with a previous study in rats^91^. Sex differences can also be contextual in that they only emerge in certain conditions^87^, such as after exposure to stressors^1^. Sex differences in response to stress are partially related to sex differences in basal and stress-evoked corticosterone levels^92^, although basal corticosterone levels in PFC do not appear to differ in male and female rats^93^. Although there were no overt stressors in the reversal learning task, it is possible that stress associated with water deprivation or handling differently affected males and females in our experiments contributing to differences in their behavior.

Ultimately, many of these factors likely combine and interact to dictate motivation in individual mice^87, 94^. Moreover, factors other than sex influenced motivation in the present study (**Supplementary Figure 1e-g**). Further work is needed to disentangle the complicated interactions between the myriad factors that influence motivation such as thirst, hunger, body weight, physiological responses to water deprivation and sucrose, and how they may be affected by sex.

### Relationship to previous studies on the role of ACC and DMS in value-based decision making

Our work was motivated by studies implicating the ACC and DMS in decision-making. The ACC is thought to be important for updating behavior based on recent outcomes: ACC activity modulates post-error slowing of response times^95, 96^, affects persistence of behavioral strategy^68^, modulates switching away from a behavior that is no longer rewarding^66, 97^ and correlates with the decision of when to act ^98^. Projections from cortex to the striatum are thought to carry information necessary for learning and performance of value-based behavior^28, 31, 52, 99–103^. One of the major cortical inputs to the DMS is from the ACC^26, 104–106^. This projection is thought to facilitate appropriately timed motor responses during cognitive tasks^107, 108^, and is important for cost-benefit evaluation^27^. Together, these results suggest that the ACC (and its projection to the DMS) is important for controlling when to engage a particular behavior or strategy based in part on the expected value of available options. This is consistent with our finding that, at least in females, ACC-DMS neurons regulate motivation to perform a decision-making task based on the value of the chosen action (**Figure 2**).

What information is transmitted from the ACC to the DMS had not been previously elucidated, however. Consistent with recordings from the ACC^,18, 34, 42, 61, 109–117^ and DMS^36, 53, 118– 124^, we identified neural correlates of the value of available options (**Figure 6**), the animal’s choices (**Figure 5**) and the outcome of those choices (**Figure 4**) in ACC-DMS neurons. A caveat for these analyses is the possibility that we were unable to perfectly isolate individual neurons due to the lack of z-plane sectioning during single-photon imaging. However, we have no reason to believe this differently affects males and females, so it is unlikely to change our conclusions.

Perhaps most interestingly, our recordings and perturbations of ACC-DMS neurons add to this previous literature by revealing an unexpected dissociation between the neural substrates of choice and motivation. Specifically, both neural correlates and perturbations revealed sex differences related to motivation and not choice.

Because many studies of decision making do not measure motivation to engage in the task, it is not clear to what extent the neural regulation of motivation and choice is dissociated across the brain. However, manipulation of the medial PFC ^29, 46, 125^ affects both choice and latencies, while inhibition of the basolateral amygdala^126^ can affect choice without affecting response latencies. The DMS itself has been implicated in motivation^52, 127, 128^ and choice^36^ in goal-directed behavior^30, 129^. Thus, our results suggest that while ACC input may not be necessary for the DMS’s role in choice, it may be important for the decision of whether and when to engage in the behavior in the first place.

## Supplementary figures

**Supplementary Figure 1:**
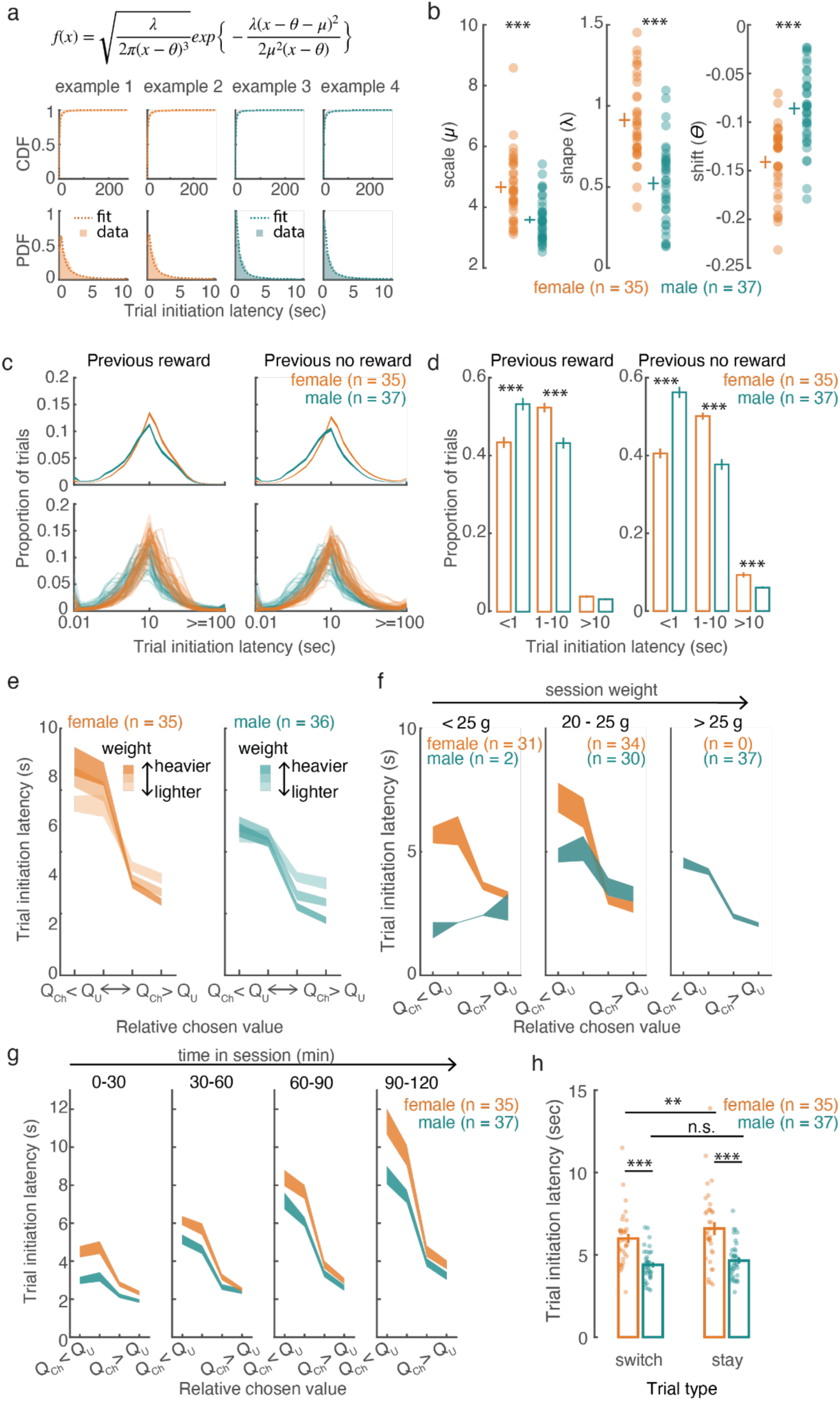
Sex differences in motivation to perform a value-based decision making task in mice. **a.** To quantify sex differences in the distribution of trial initiation latencies, we fit each animal’s data with a shifted inverse gaussian distribution. Top: probability distribution function for the shifted inverse gaussian. Middle: Cumulative distribution of trial initiation latencies (shaded line) and fit (dashed line) for example females (orange) and males (green). Bottom: Probability distribution function of trial initiation latencies (shading) and fit (dashed line) for example females (orange) and males (green) for 1-10 seconds. **b.** Parameter estimates for each mouse. Translucent circles are individual animals, crosses are mean and SEM across males or females. Comparisons between males and females were performed with 2-sided Wilcoxon rank sum tests. μ: Z = -4.44, p = 9.04x10^-6^; λ: Z = -5.55, p = 4.65x10^-8^; θ: Z= 5.18, p = 2.19x10^-7^. **c.** Histograms of trial initiation latencies binned in log-space for males and females following rewarded and unrewarded trials. Top panels are the mean and SEM across animals, and the bottom panels show the histograms separately for each animal. Trials preceded by reward or no reward are plotted separately. **d.** Trial initiation latencies binned as short (<1 s), medium (1-10 s) and long (>10 s) for previously rewarded and unrewarded trials for males and females. Males had more short trials than females (2-sided Wilcoxon rank sum tests, reward: Z = -3.90, p = 9.7x10^-5^; unrewarded: Z = -5.99, p = 2.1x10^-9^). Females had more medium trials than males (2-sided Wilcoxon rank sum tests, rewarded: Z = 4.20, p = 2.6x10^-5^; unrewarded: Z = 5.62, p = 1.9x10^-8^). Females had more long trials only following unrewarded outcomes (2-sided Wilcoxon rank sum tests, rewarded: Z = 1.37, p = 0.17; unrewarded: Z = 4.19, p = 2.78x10^-5^). **e.** Daily fluctuations in weight affected value modulation of trial initiation latencies similarly in males and females (mixed effects regression of latency with sex, weight, relative chosen value, trial number and their interactions as fixed effects; see **Supplementary Table 5** for details. *weight*: F(1,56.35) = 8.67, p = 0.004; *sex:weight:* F(1,56.35) =12.84, p =7.10x10^-4^; *relative_chosen_value:weight*: F(3,135)=28.71, p = 2.00x10^-14^; *sex:relative_chosen_value:weight*: F(3,135) = 1.63, p = 0.18. For each mouse, sessions were binned in terciles of weight and trials were divided into quantile bins of relative chosen value. Trial initiation latencies were then averaged for each bin. n=35 females and 36 males. One male maintained constant weight throughout the experiment and was excluded from this plot. **f.** Sessions were divided based on weight, and trial initiation latencies were binned in 4 quantile bins of relative chosen value and averaged for each animal. **g.** Average trial initiation latency versus relative chosen value in 30 min bins for males and females. Trials were binned based on time in session as well as in quantiles of relative chosen value and averaged for each mouse. Trial number significantly affected trial initiation latencies but there was no effect of trial on the interaction between relative chosen value and sex (see **Supplementary Figure 5** for details; *trial:* F(1,66.30) = 390.61, p = 1.71x10^-29^; *sex:trial*: F(1,66.30) = 4.17, p = 0.05; *sex:trial:relative_chosen_value*: F(3,135) = 1.63, p = 0.18). **h.** Following unrewarded trials, only females modulated their trial initiation latencies based on the upcoming stay versus switch decision. There was a significant effect of sex, stay/switch and a significant interaction between sex and stay/ switch, reflecting greater modulation of trial initiation latency by whether or not the upcoming trial was a stay or switch trial in females compared to males (mixed-effects regression: *latency ∼ sex + stay + sex:stay + (1+stay|subject)*; Sex: F(1,346950) = 653.13, p = 6.34x10^-144^; Stay: F(1,37.39) = 8.63, p = 0.006; Sex:Stay: F(1,18461) = 8.26, p = 0.004). Females were significantly slower than males to initiate stay and switch trials (2-sided Wilcoxon rank sum tests. switch male vs. female: Z = 3.97, p = 7.3x10^-5^; stay male vs. female: Z = 4.69, p = 2.77x10^-6^). In females, trial initiation latencies for switch trials were slower than stay trials. There was no difference between trial types in males (2-sided Wilcoxon signed rank tests. Female: Z = 3.18, p = 0.002; Male: Z = 0.96, p = 0.34). ** p < 0.01, *** p < 0.001; n = 37 males, 35 females. Error bars or shading are standard error of the mean.

**Supplementary Figure 2:**
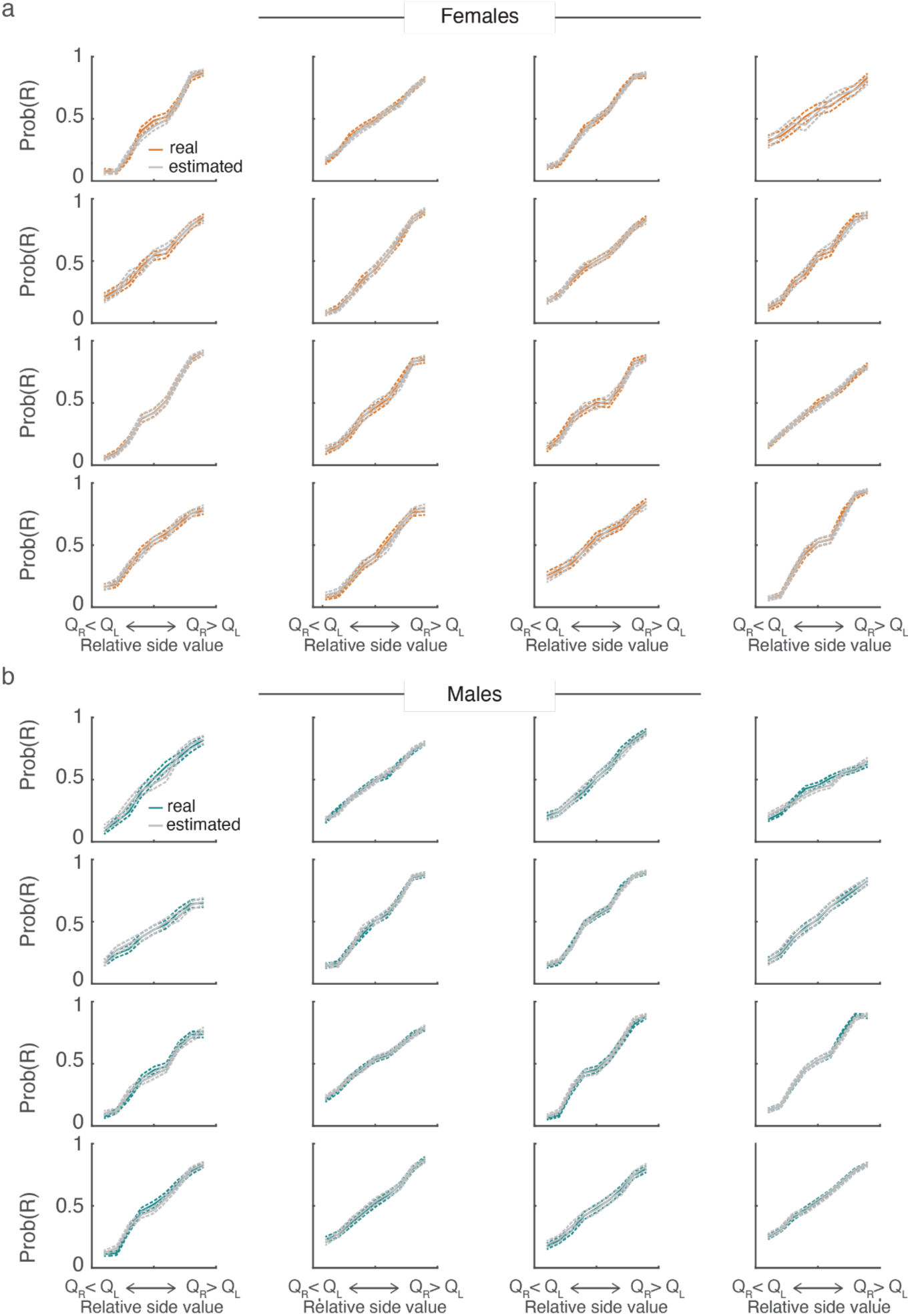
Example psychometric curves showing the relationship between relative side value and choice for real data and data estimated from the Q-learning model. **a)** The probability of making a right choice was calculated in quantile bins of relative side value for the mouse and model for all trials and plotted for randomly selected mice. Error bars are 95% binomial confidence intervals. **b)** Same as **a**, for male mice.

**Supplementary Figure 3:**
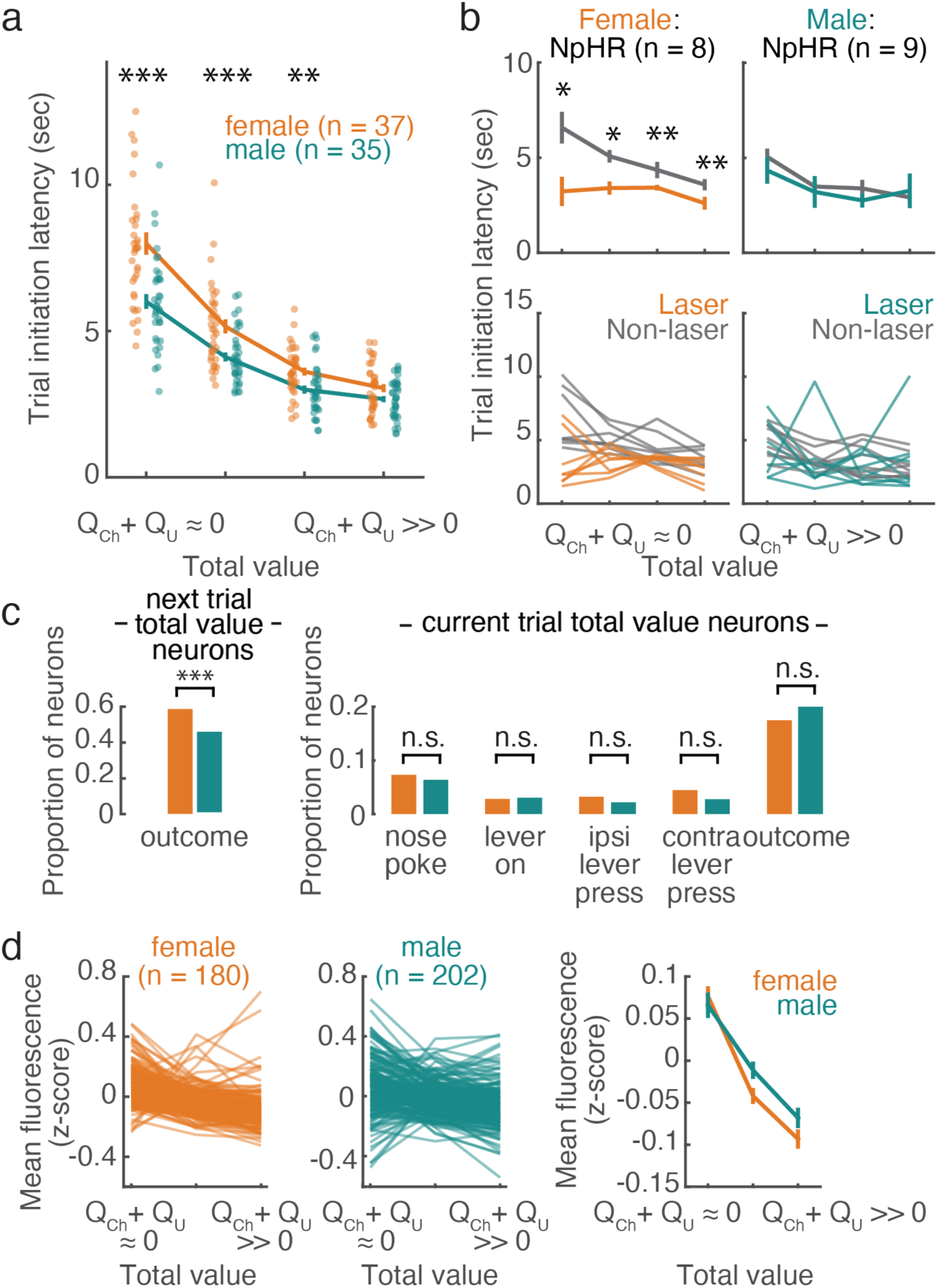
Trial initiation latency was modulated by total value. **a)** Trial-by-trial total value (*Q_Ch_* + *Q_U_*, where *Q_Ch_* and *Q_U_* are the action values for the chosen and unchosen lever, respectively) was divided into quartiles for each animal, and trial initiation latencies were averaged in quartile bins. To compare how total value influenced trial initiation latency in males and females we fit the following mixed effects regression: *trial_initiation_latency ∼ Sex + Total_value + Trial + Weight + Sex:Total_value + Trial:Total_value + Trial:Sex + Weight:Total_value + Weight:Sex + Weight:Sex:Total_value + Trial:Sex:Total_value + (1 + Total_value + Weight + Trial + Weight:Total_value + Trial:Total_value|subject) + (1 + Total_value + Trial + Total_value:Trial|session:subject)*. Model was fit with effects coding and significance was determined with F-tests using the Satterthwaite method to estimate degrees of freedom. Total value and the interaction between sex and total value significantly affected trial initiation latency (*Sex:* F(1,61.56) = 3.95, p = 0.05; *Trial:* F(1,66.21) = 383.55, p < 0.001; *Total_value:* F(3,109.58) = 59.42, p < 0.001; *Weight:* F(1,58.75) = 5.92, p = 0.02; *Trial:Sex:* F(1, 66.21) = 2.14, p = 0.15; *Trial:Total_value:* F(3,91.4) = 55.30, p < 0.001; *Sex:Total_value:* F(3, 109.58) = 15.99, p < 0.001; *Sex:Weight:* F(1, 58.75) = 13.03, p<0.001; *Total_Value:Weight:* F(3, 142.71) = 15.66, p < 0.001; *Trial:Sex:Total_value:* F(3, 91.4) = 0.22, p = 0.88; *Weight:Sex:Total_value:* F(3, 142.71) = 1.94, p = 0.13). Specifically, females were slower than males to initiate low total value trials (2-sided Wilcoxon rank sum tests between males and females: bin 1: W = 1638, p = 4.99x10^-5^; bin 2: W = 1582, p = 4.99x10^-4^; bin 3: W = 1556, p = 0.002; bin 4: W = 1433 p = 0.08). n = 37 females, 35 males. **b)** Inhibition of ACC-DMS neurons decreased the influence of total value on trial initiation latency in females. To determine how the effect of inhibition on trial initiation latencies was influenced by sex and opsin, we divided trials into quartile bins of total value and averaged trial initiation latencies for laser and non-laser trials in each bin. We then fit a linear mixed-effects model to the difference between laser and non-laser trials with sex, total value, opsin and their interactions as fixed effects and random intercepts for each subject. The effect of inhibition depended on sex and opsin (*sex*opsin*: F(1,29) = 4.86, p = 0.04, *sex:* F(1,29) = 14.52, p = 6.68x10^-4^; *opsin:* F(1,29) = 3.02, p = 0.09; *total value*: F(1,29) = 4.10, p = 0.02, *sex*total value quantile:* F(3,29) = 1.42, p = 0.26; *opsin*total value quantile:* F(3,29) = 0.41, p = 0.75, *sex*opsin*total value quantile*: F(3,29) = 0.55, p = 0.65). Specifically, females expressing NpHR were significantly faster to initiate trials following inhibition compared to control trials (2-sided Wilcoxon rank sum tests, bin 1: W=35, p = 0.02; bin 2: W=35, p = 0.02; bin 3: W=36, p = 0.008, bin 4: W=36, p = 0.008). In males, trial initiation latencies for laser and non-laser trials did not differ in any bin (2-sided Wilcoxon signed rank tests, p>0.2 for all comparisons). n = 8 females, 9 males. **c)** Significantly more neurons encoded upcoming total value during the outcome in females than males (*χ* ^2^-test, *χ* ^2^ (1,N=756) = 13.58, p = 2.3x10^-4^). Proportions of neurons encoding current trial total value did not differ between males and females for any task event (*χ* ^2^-tests, all p > 0.3). **d)** Outcome activity was averaged across time (8 seconds from presentation of the outcome) and averaged in 3 quantile bins of total value. The left plots show the value modulation for all total value encoding neurons in females and males and the right-most plot shows the mean and SEM across neurons for males and females. (**a,b**) Circles are data from individual mice and lines show across animal averages and error bars standard error of the mean. * p<0.05, ** p<0.01, ***p<0.001

**Supplementary Figure 4:**
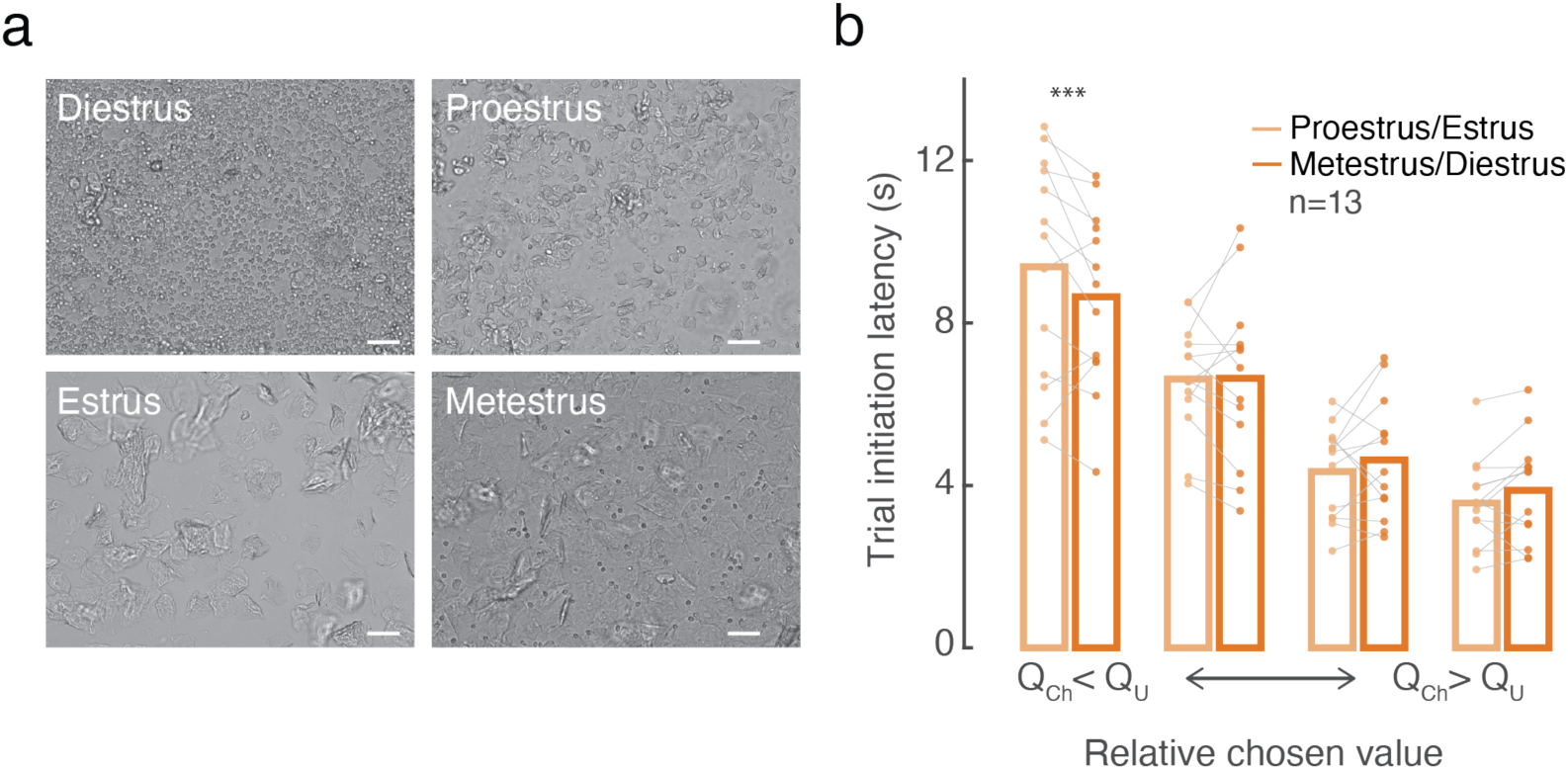
Estrous cycle modulates trial initiation latencies. **a.** Representative images of vaginal cells across the estrous cycle. Scale bar is 50*μ*m. **c.** Estrous cycle significantly modulated the relationship between relative chosen value and trial initiation latency. (Mixed effects regression, significance assessed with F tests using the Satterthwaite method to estimate degrees of freedom: *trial initiation latency ∼ relative_chosen_value + estrous_stage + relative_chosen_value:estrous_stage + (1+relative_chosen_value|subject)*). *Relative_chosen_value*: F(3,16.56) = 90.91, p = 1.71x10^-10^; *Estrous_stage*: F(1,178080) = 10.77, p = 4.51x10^-7^; *Relative_chosen_value:Estrous_stage*: F(9,19140) = 2.98, p = 0.002. Post-hoc comparisons of trial initiation latencies during proestrus/estrus and metestrus/ diestrus for each value bin were performed with F-tests for each contrast using the Satterthwaite method to estimate degrees of freedom (bin 1: F(1,63731) = 11.88, p = 5.68x10^-4^; bin 2: F(1,33322) = 0.37, p = 0.55; bin 3: F(1,10564) = 1.91, p = 0.17; bin 4: F(1,4098) = 2.49, p = 0.11). *** p < 0.001.

**Supplementary Figure 5:**
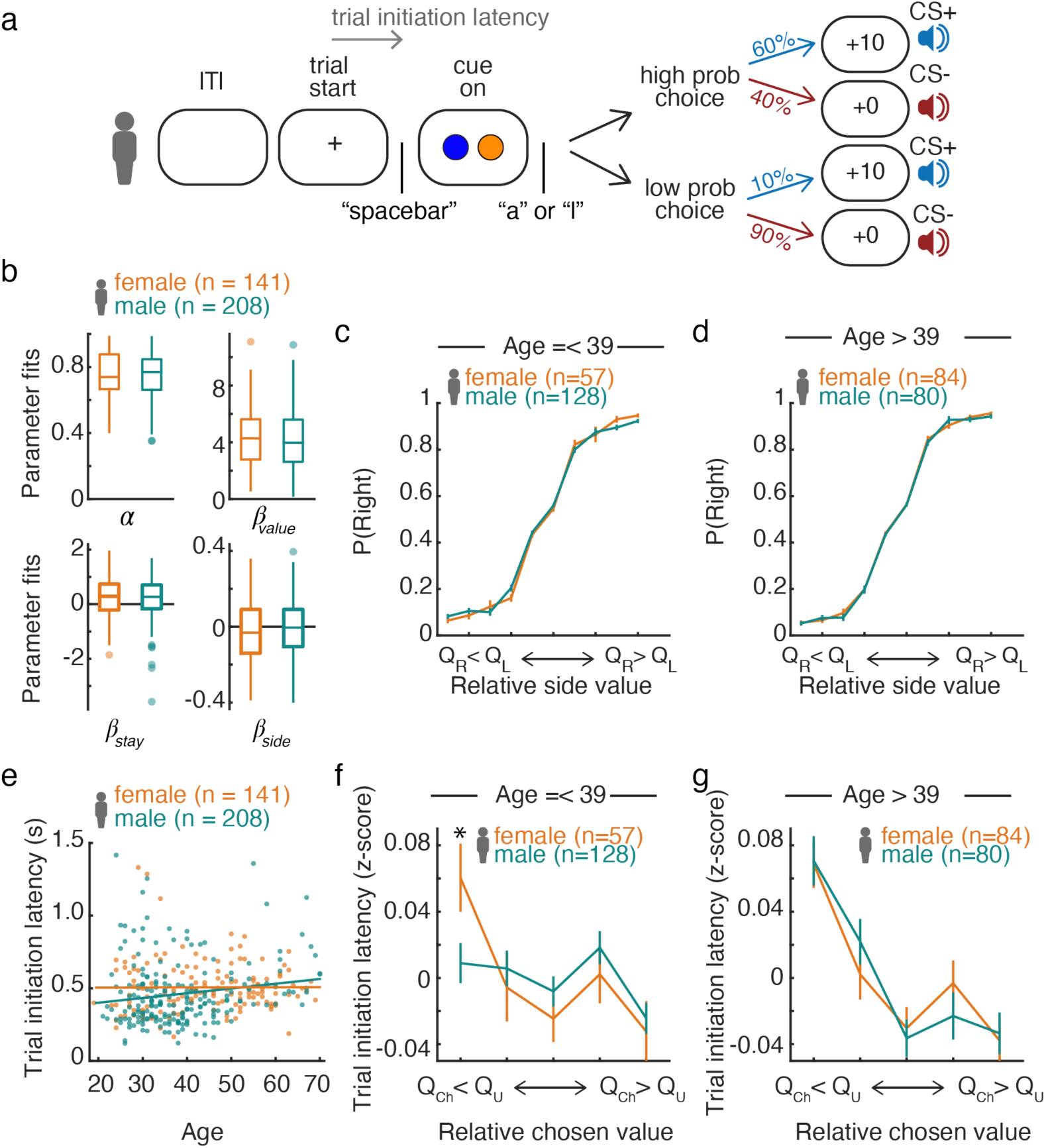
Gender differences in human subjects performing the self-initiated probabilistic reversal learning task. **a)** Schematic of the online task. After an intertrial interval (ITI), trial start was cued with the appearance of a plus sign (“+”) on the screen. Subjects initiated a trial by pressing the spacebar which led to the presentation of 2 colored circles on either side of the screen. Subjects indicated their choice by pressing the “a” key for a left choice and the “l” key for a right choice. The outcome was then presented, indicating the number of points received for that choice (rewarded: +10, unrewarded +0) accompanied by auditory cues indicating reward (bell-like sound) or no reward (buzzer). High probability choices were rewarded 60% of the time and low probability choices 10% of the time and the identity of the high and low choice alternated as in the mouse task (see **Methods** for details). **b)** Estimates of Q-learning parameters fit with the same hierarchical model used for the mice. **c-d)** Choice was similar between men and women. There was a significant main effect of relative side value, but no effect of gender on how relative side value modulated the probability of choosing the right option (mixed-effects, logistic regression: *choice ∼ relative_side_value_quantile + gender + age + sex: relative_side_value_quantile + sex: age + relative_side_value_quantile: age + sex*relative_side_value_quantile:age + (1 + relative_side_value_quantile|subject)*. *relative side value*: F(1,142180) = 100.5, p = 8.58x10^-209^; *gender*: F(1,142180) = 1.20, p = 0.27; *age*: F(1,142180) = 0.98, p = 0.32; *relative_side_value:gender*: F(1,142180) = 0.72, p = 0.71; *gender:age*: F(1,142180) = 1.26, p = 0.26; *relative_side_value:age*: F(10,142180) = 0.73, p = 0.70; *relative_side_value:gender:age*: F(1,142180) = 0.62, p = 0.80, gender was a categorical variable and age was z-scored. Significance of coefficients was assessed with F-tests). **c)** Trials were divided into 11 quantile bins of relative side value and the probability of making a right choice was averaged by bin for each subject younger than or equal to the median age (between 19 and 39 years old). Error bars are standard error of the mean. **d)** Same as **c,** except for subjects 40-70 years old. **e)** Trial initiation latencies were significantly affected by age in men, but not women (linear correlation, males: r = 0.16, p = 0.02, females: r = -0.006, p = 0.95). **f-g)** The modulation of trial initiation latency by value differed in males and females, and this effect was modulated by age. (mixed-effects regression with relative chosen value quantile, gender, age and their interactions as fixed effects and random effects of subject. (*relative_chosen_value*: F(4,1754) = 3.46, p = 0.008: *gender* : F(1, 1754) = 1. 68 x 10^-4^, p = 0. 99; *age* : F(1, 1754) = 1. 62 x 10^-4^, p = 0. 99; *relative_chosen_value:gender*: F(4,1754) = 2.49, p = 0.04; *relative_chosen_value:age*: F(4,1754) = 6.71, p = 2.36x10^-5^; *gender:age*: F(1,1754) = 1.16x10^-4^, p = 0.99; *relative_chosen_value:gender:age*: F(4,1754) = 2.46, p = 0.04; n = 141 women, 209 men). **f)** Trials were divided into 5 quantile bins of relative chosen value for males and females 19-39 years old and trial initiation latencies were averaged by bin. Two-sided Wilcoxon rank sum tests showed that women were significantly slower than men to initiate trials in the lowest quantile bin of relative chosen value (bin 1: Z = -2.01, p = 0.04, bin 2: Z = –0.10, p = 0.92, bin 3: Z = 0.79, p = 0.43, bin 4: Z = 1.35, p = 0.18, bin 5: Z = 0.54, p = 0.59) **g)** Same as **F** except for subjects aged between 40 and 70. There were no significant differences in trial initiation latencies between males and females older than 39 (2-sided Wilcoxon rank sum tests, p>0.07). *Q_Ch_*: action value for the chosen lever; *Q_U_*: action value for the unchosen lever; *Q_R_*: action value for the right lever; *Q_L_*: action value for the left lever; * p < 0.05.

**Supplementary Figure 6:**
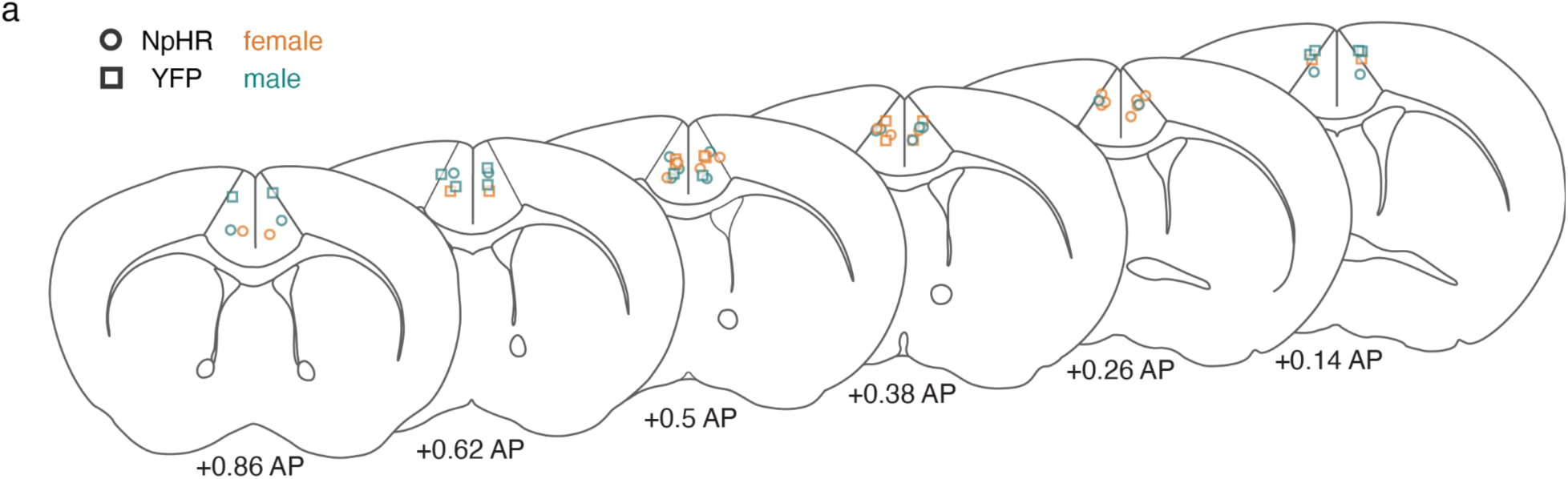
Location of fiber tips for optogenetic inhibition. **a.** Circles indicate animals expressing eNpHR and squares indicate animals expressing EYFP. Orange: females, green: males.

**Supplementary Figure 7:**
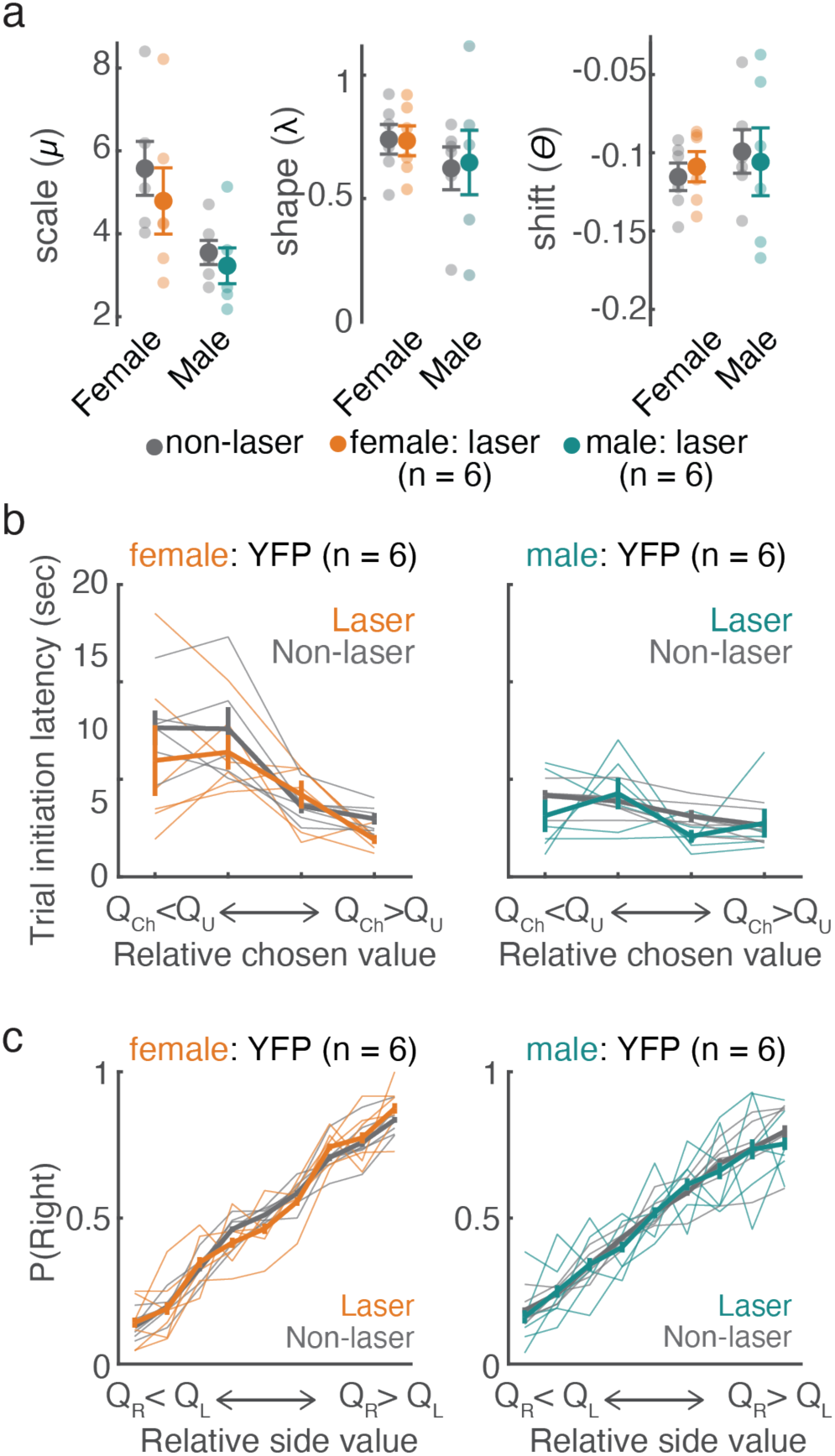
Laser during outcome presentation had no effect on trial initiation latencies or choice in mice expressing EYFP in ACC-DMS neurons. **a.** Comparisons between shifted inverse gaussian fits from laser and non-laser trials revealed no differences in EYFP-expressing males or females. (2-sided Wilcoxon signed rank tests, females: scale: W = 19, p = 0.09, shape: W = 4, p = 0.22 shift: W = 12, p = 0.84, males: shape: W = 15, p = 0.44, scale: W = 12, p = 0.84, shift: W = 15, p = 0.44, n = 6 females, 6 males). **b.** Trial initiation latencies averaged in quartile bins of relative chosen value (*Q_Ch_* − *Q_U_*, where *Q_Ch_* and *Q_U_* are the action values for the chosen and unchosen lever, respectively) for mice expressing EYFP in ACC-DMS neurons. Paired comparisons between laser and non-laser trials were performed for each bin with 2-sided Wilcoxon signed rank tests (Female laser v. no laser bin 1: W = 17, p = 0.22, bin 2: W = 19, p = 0.09, bin 3: W = 6, p = 0.44, bin 4: W = 20, p = 0.06; Male bin 1: W = 16, p = 0.31, bin 2: W = 10, p = 1, bin 3: W = 19, p = 0.09, bin 4: W = 13, p = 0.69). **c.** The probability of making a right choice was averaged in quantile bins of relative side value (*Q_R_* − *Q_L_*, where *Q_R_* and *Q_L_* are the action values for the right and left lever, respectively) for each animal and averaged.

**Supplementary Figure 8:**
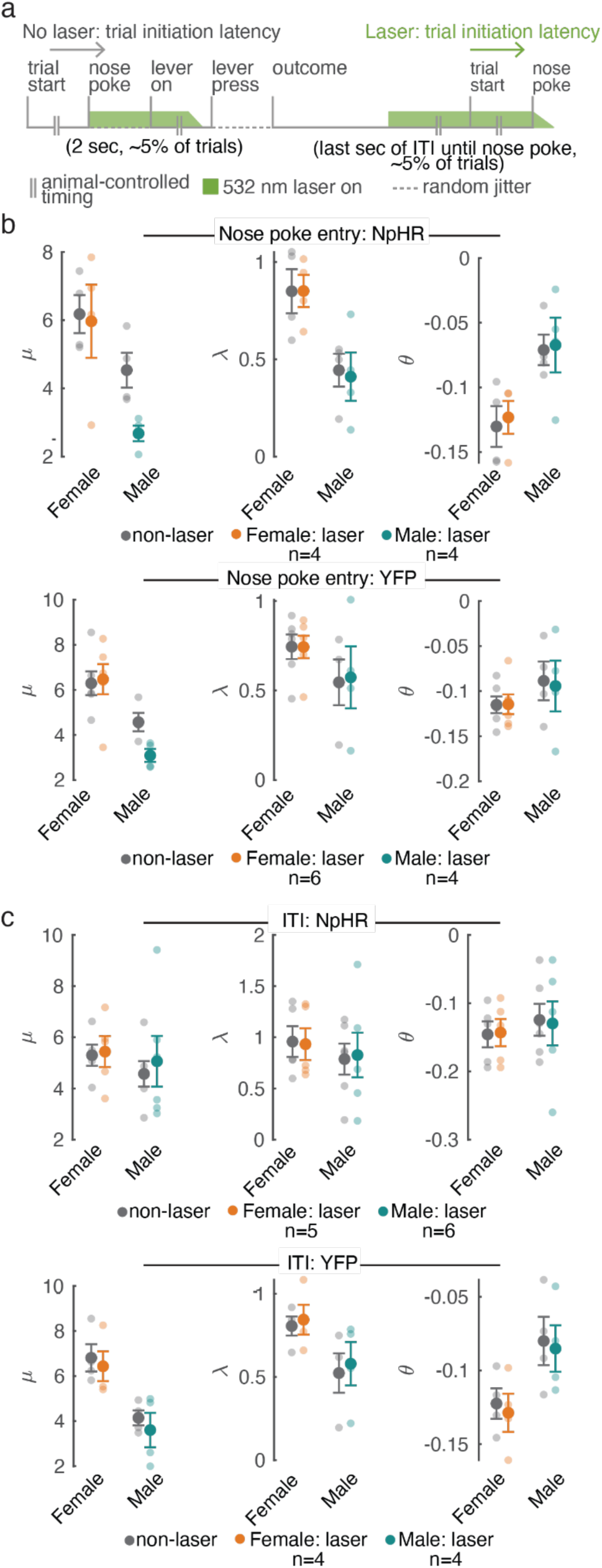
ACC-DMS inhibition during the nose poke or intertrial interval. **a.** In subsets of mice, 5 mW of 532 nm light was delivered bilaterally to the ACC for 2 s triggered by the mouse entering the nose poke to initiate the trial. Additionally, in subsets of mice 532 nm light was delivered starting 2 seconds after the presentation of the outcome and persisting until the mouse initiated the next trial, with a jittered offset between 0.05 and 1 second so that the timing of the offset of the laser was less contingent on the mouse’s behavior. **b**. Estimated parameters from nose poke laser and non-laser trials fit with a shifted inverse gaussian distribution. Data from mice expressing NpHR in ACC-DMS neurons are plotted in the top panels and mice expressing EYFP are plotted in the lower panels. Comparisons between laser and control parameters were performed with 2-sided Wilcoxon signed rank tests (n = 4 female NpHR, 4 male NpHr, 6 female EYFP, 4 male EYFP, p>0.1 for all comparisons). There was a non-specific effect of laser on the scale parameter in males (both EYFP- and NpHR-expressing mice), but this did not reach significance (2-sided Wilcoxon signed rank tests: male eNpHR: W = 10, p = 0.12; male EYFP: W = 10, p = 0.12). **c.** Same as **b** for inhibition during the ITI. There were no significant effects on trial initiation latencies from inhibition during the late ITI. Comparisons between laser and control parameters were performed with 2-sided Wilcoxon signed rank tests (n = 5 female NpHR, 6 male NpHr, 4 female EYFP, 4 male EYFP, p > 0.1 for all comparisons). Translucent circles show the fits from each animal and the opaque circles are averaged across animals.

**Supplementary Figure 9:**
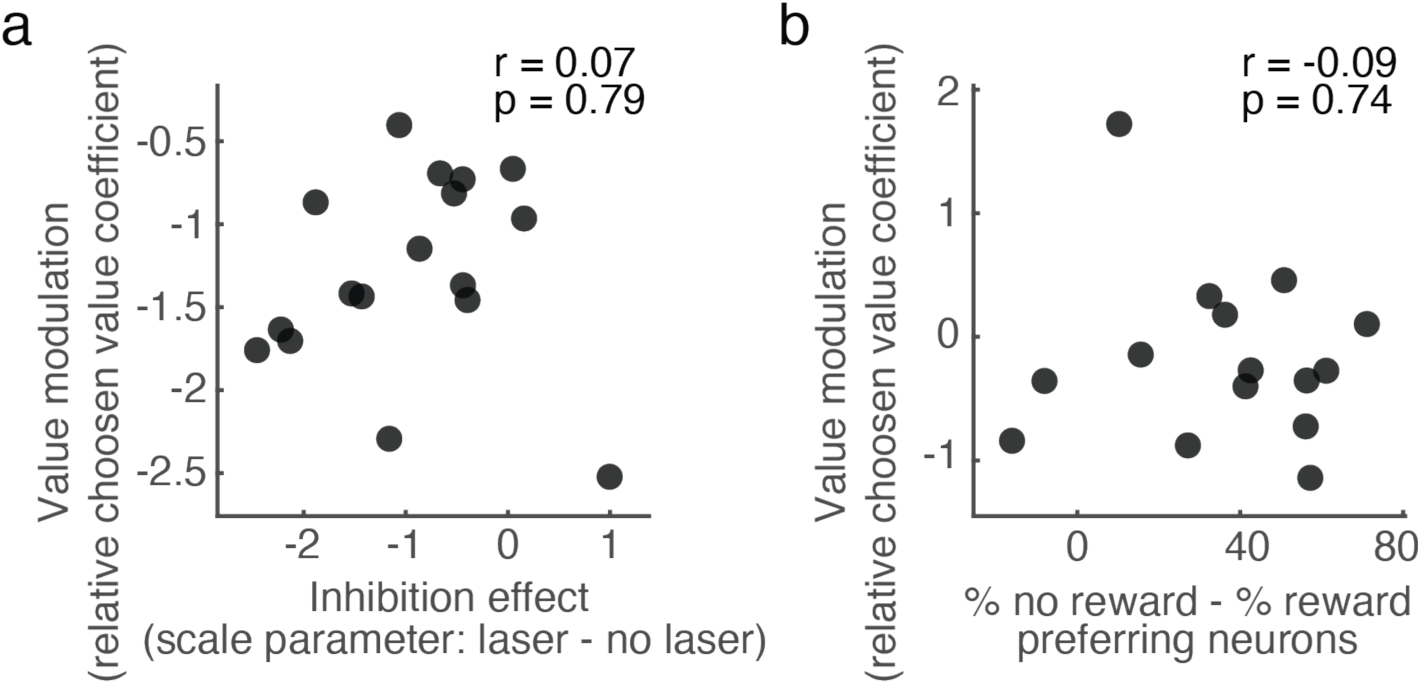
Individual variability in optogenetic and imaging results. **a.** Inhibition effect size did not correlate with value modulation of trial initiation latencies during control sessions (linear correlation, inhibition effect vs. value modulation: r = 0.07, p = 0.79, p = 17 mice). For each mouse, the inhibition effect was quantified as the difference between the scale parameter of the inverse gaussian distribution fit separately for laser and control trials (**Figure 2d**). To estimate modulation of trial initiation latencies by relative chosen value for each mouse, we fit the following mixed-effects regression to the trial initiation latencies for each mouse separately: *latency ∼ relative_chosen_value + (1+relative_chosen_value|session)*. The value coefficient reflects how much trial initiation latencies were modulated by relative chosen value. **c.** The bias towards negative outcome encoding did not significantly correlate with value modulation of trial initiation latencies (linear correlation, %no reward-%reward encoding neurons vs. value modulation of trial initiation latencies: r = -0.09, p = 0.74, n = 15 mice). For each mouse, bias towards negative outcome encoding was defined as the difference in the proportion of no-reward preferring outcome encoding neurons and the proportion of reward-preferring outcome encoding neurons. To estimate modulation of trial initiation latencies by relative chosen value for each mouse, we fit the following mixed-effects regression to the trial initiation latencies for each mouse separately: *latency ∼ relative_chosen_value + (1+relative_chosen_value|session)*. The value coefficient reflects how much trial initiation latencies were modulated by relative chosen value.

**Supplementary Figure 10:**
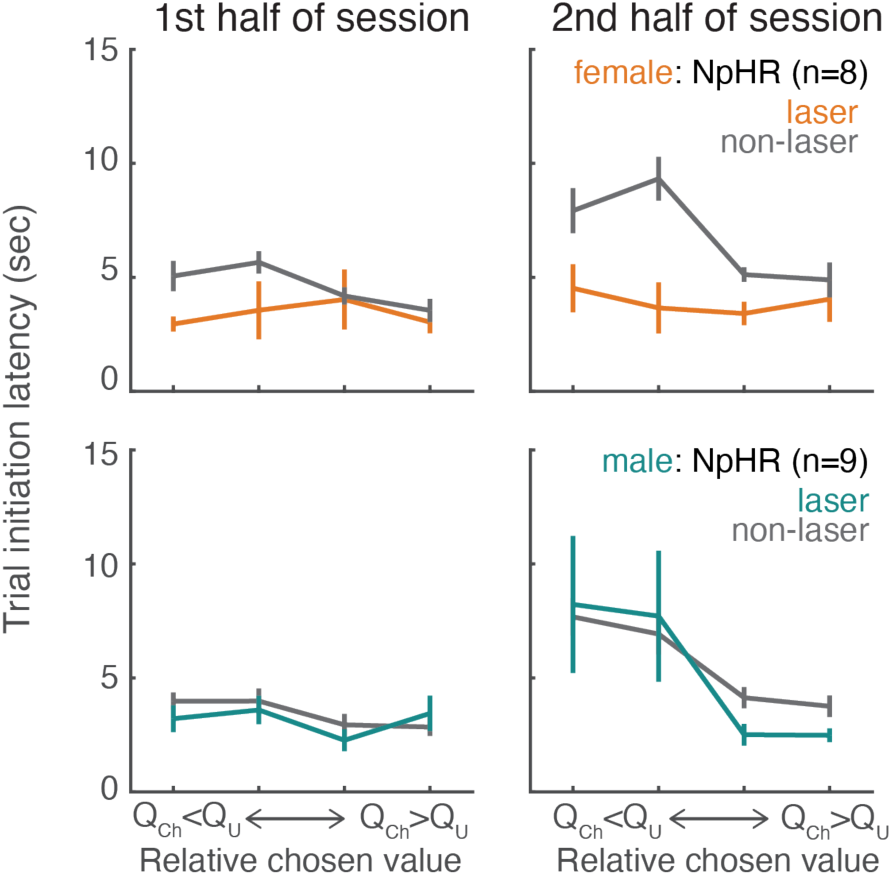
Inhibition affected trial initiation latencies similarly in the first and second half of the session. Trials were divided based on whether they were initiated during the first or second hour of the session and divided into 4 quantile bins of relative chosen value and averaged for each mouse. To quantify the effect of ACC-DMS inhibition in males and females across the session we used a mixed-effects regression of the difference in trial initiation latencies on laser and control trials with sex, opsin, relative chosen value quantile bin and time bin (first or second half of the session) as fixed effects and random intercepts and effects of value for each subject (See **Supplementary Table 8** for details). There was a significant effect of sex:opsin (F(1,47.68) = 5.09, p = 0.03) and a significant effect of sex:opsin:relative_chosen_value (F(3,52.07) = 3.52, p = 0.02) but the interactions with time were not significant (sex:opsin:time_bin: F(1,174) = 0.81, p = 0.37, sex:opsin:relative_chosen_value:time_bin: F(3,174) = 2.26, p = 0.08). To further characterize these effects, we estimated latencies separately in males and females during the first and second half of the session with laser, relative chosen value and laser:relative chosen value as fixed effects and random effects of subject. There was a significant effect of laser in females for both the first and second half of the session (first half: *laser*: F(1,18682) = 6.60, p = 0.01; *relative_chosen_value*: F(3,18682) = 1.11, p = 0.34; *laser:relative_chosen_value*: F(3,18682) = 1.58, p = 0.19. Second half: *laser*: F(1,15522) = 8.78, p = 0.003; *relative_chosen_value*: F(3,15522) = 1.30, p = 0.27; *laser:relative_chosen_value*: F(3,15522) = 1.28, p = 0.28). In contrast, there was no effect of laser in males in either half of the session (first half: *laser*: F(1,21160) = 0.924, p = 0.34; *relative_chosen_value*: F(3,21160) = 2.23, p = 0.08; *laser:relative_chosen_value*: F(3,21160) = 0.79, p = 0.50. Second half: *laser*: F(1,17580) = 0.57, p = 0.45; *relative_chosen_value*: F(3,17580) = 5.29, p = 0.001; *laser:relative_chosen_value*: F(3,17580) = 0.20, p = 0.90).

**Supplementary Figure 11:**
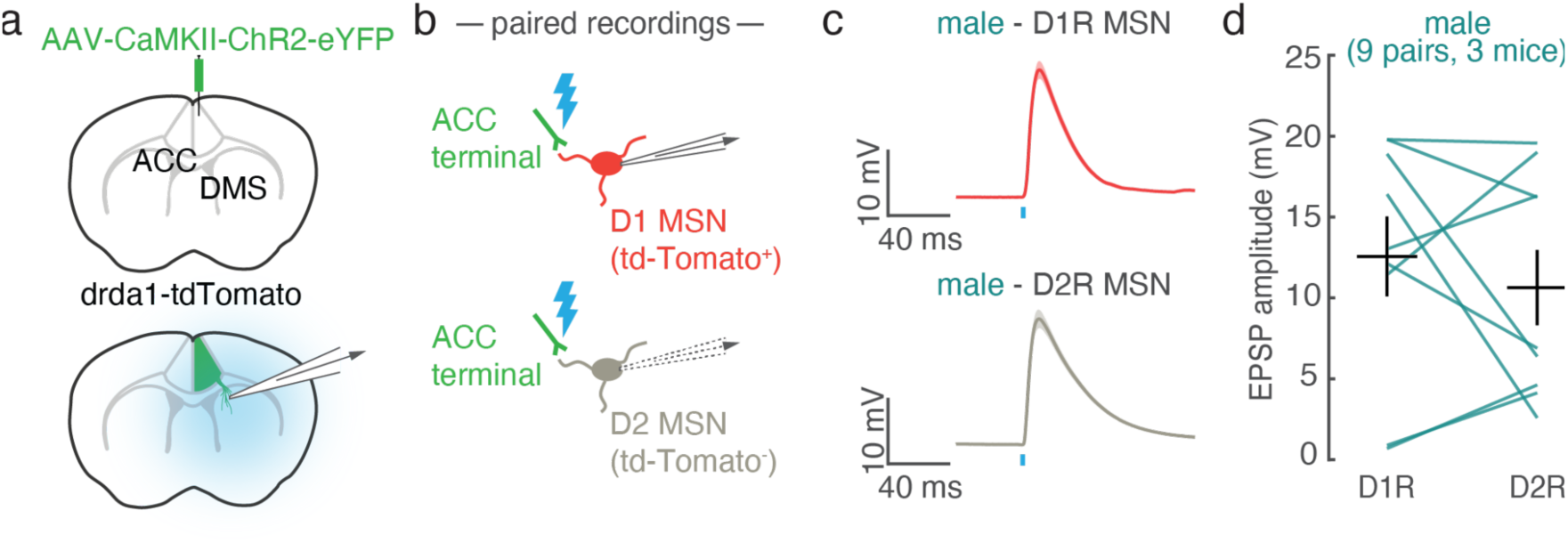
Similar excitatory postsynaptic potentials evoked by ACC-DMS stimulation in D1R and D2R MSNs in males. **a.** Schematic of viral strategy to express ChR2 in ACC neurons (top), and record optogenetically-evoked excitatory postsynaptic potentials (EPSPs) in DMS (bottom). **b.** Schematic of paired, sequential recordings of neighboring MSNs, where MSNs were visually identified as D1R MSNs (tdTomato+) or D2R MSNs (tdTomato-). Brief light pulses elicited EPSPs from ChR2-expressing ACC terminals. **c.** Example EPSPs measured in pairs from a D1R MSN (red) and D2R MSN (grey). Traces are mean responses across trials from a single cell. Shading is SEM. Blue line indicates the time of light stimulation. **d.** Summary of EPSP amplitudes. Each line is data from a pair of MSNs. A mixed effects regression revealed no effect of cell-type on EPSP. The model was *EPSP ∼ msn_type + (1|subject) + (1|subject:pair)*. Significance was assessed with an F-test using the Satterthwaite method to estimate degrees of freedom; *msn_type*: F(1,9) = 0.67, p = 0.43. N = 3 mice, 9 pairs.

**Supplementary Figure 12:**
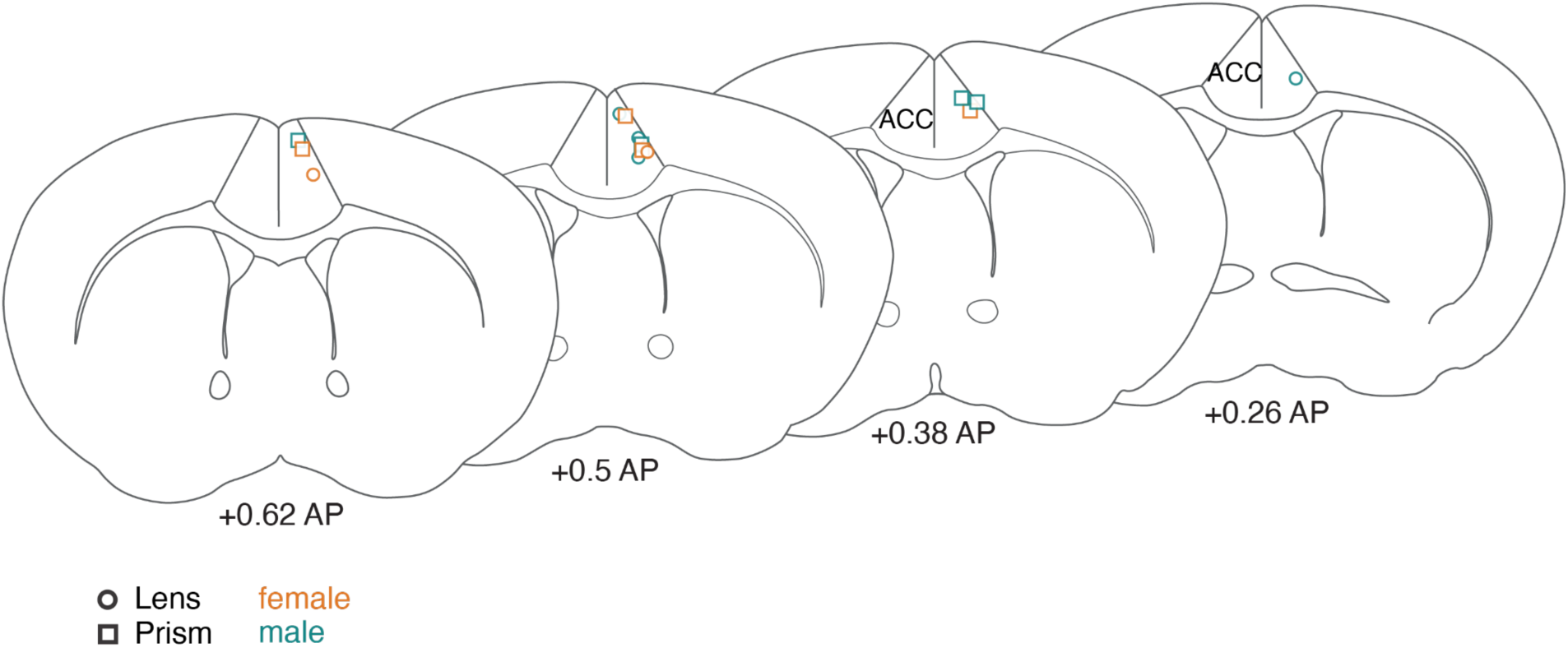
Center of imaging fields in ACC. Color indicates whether the subject was male (green) or female (orange). Circles indicate lens implants and squares are prism implants.

**Supplementary Figure 13:**
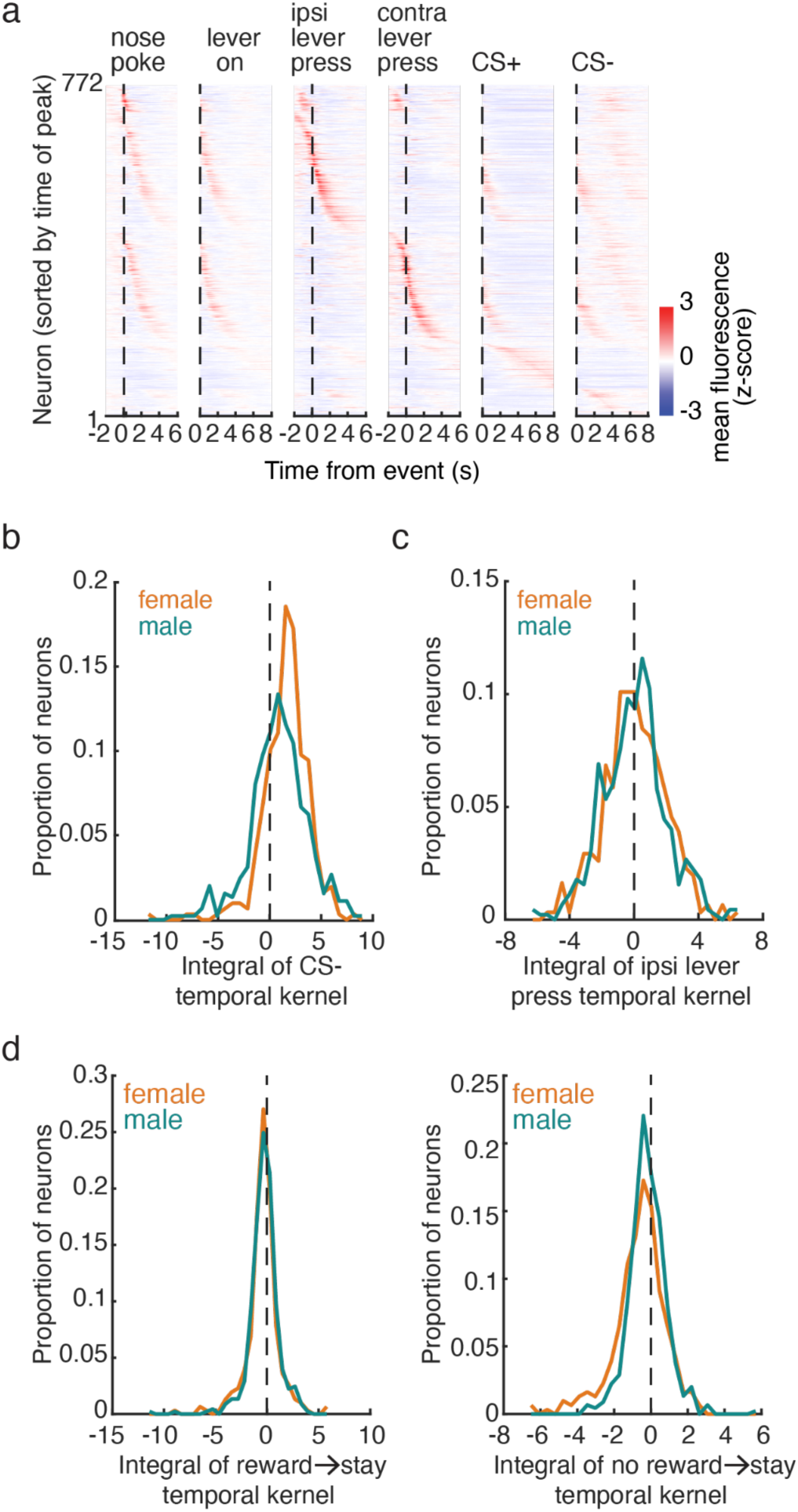
Task-related activity of ACC-DMS neurons. **a)** Trial-averaged, time-locked fluorescence to each task event for all imaged neurons. Neurons are sorted by the time of their peak fluorescence. **b)** Histograms showing the distribution of all imaged neurons, of the approximate numerical integral of the response kernel for the “no reward” temporal kernel from the regression used to assess significant outcome encoding. The histograms are plotted separately for male and female neurons. The sign of the integral of the “no reward” response kernel was used to classify neurons as reward-preferring (negative) or no reward-preferring (positive). **c)** Same as **b** for the “ipsilateral lever press” event in the regression used to assess significant choice encoding. The sign of the integral was used to classify neurons as contra-preferring (negative) or ipsi-preferring (positive). **d)** Same as **b** for the “reward to stay” event (left) and the “no reward to stay” event (right) in the regression used to assess significant stay versus switch encoding. The sign of the integral was used to classify neurons as stay-preferring (positive) or switch-preferring (negative) for rewarded (left) and unrewarded (right) trials. n = 307 female neurons, 449 male neurons.

**Supplementary Figure 14:**
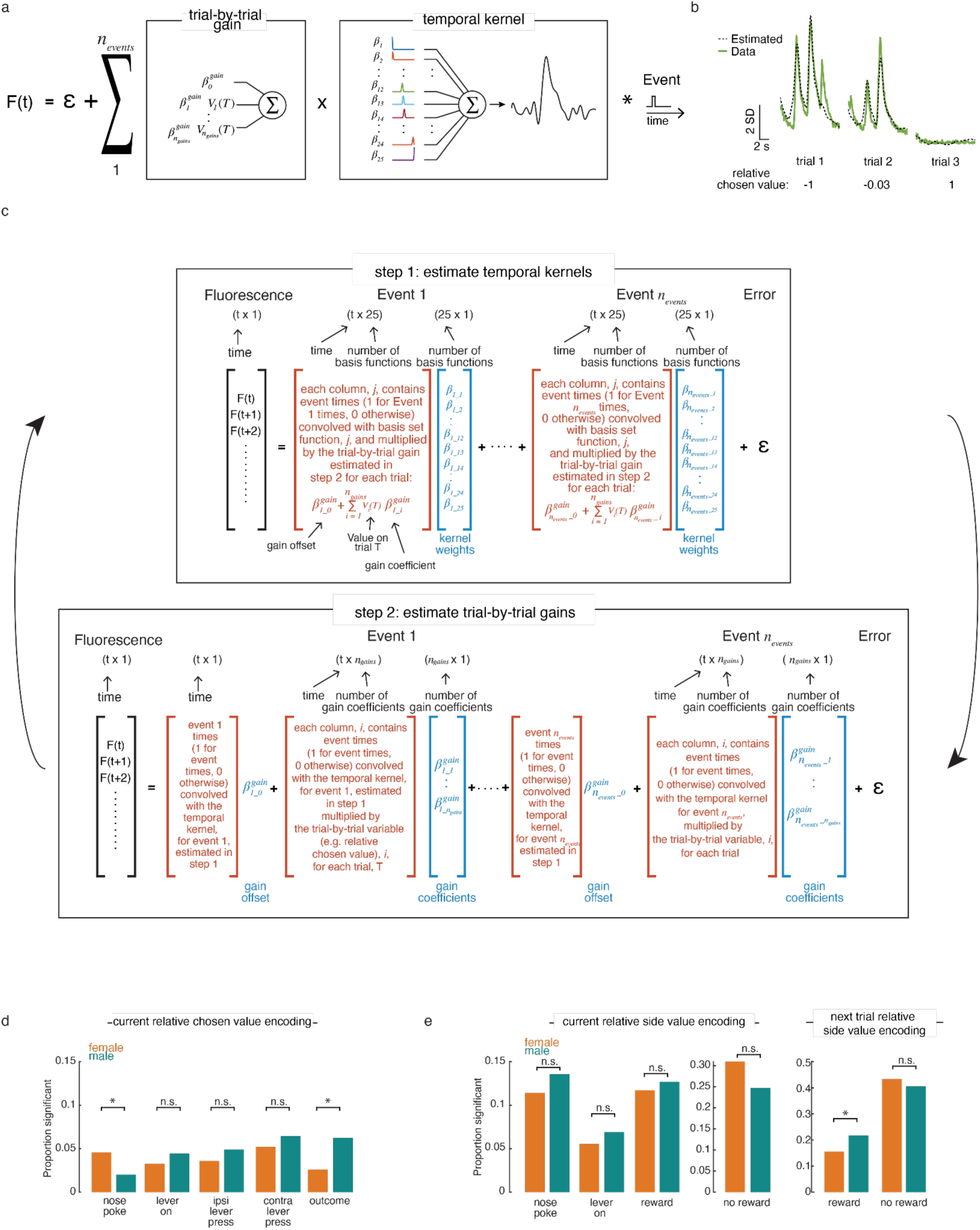
Bilinear encoding models allow value to multiplicatively modulate all event-related temporal kernels. **a.** Schematic of the encoding model used to estimate how trial-by-trial fluctuations in relative side value (*Q_R_* − *Q_L_*, where *Q_R_* and *Q_L_* are the action values for the right and left lever, respectively) or relative chosen value (*Q_Ch_* − *Q_U_*, where *Q_Ch_* and *Q_U_* are the action values for the chosen and unchosen lever, respectively) influences responses to each task event. For each neuron, GCaMP6f fluorescence was estimated as the sum of all event kernels multiplied by the gain coefficients (an offset gain as well as gains for relative chosen value (**Figure 6a-c**) or relative side value (**Figure 6d-f**)). **b.** Fluorescence (gray) and estimated fluorescence (green) from the same neuron during the outcome period on 3 trials with different upcoming relative chosen values (labeled on bottom). **c.** Schematic of the 2-step iterative fitting procedure (see **Methods** for more details). **d.** The proportion of neurons significantly encoding current relative chosen value during all task events for males and females. Significantly more neurons encoded current trial relative chosen values in females compared to males during the nose poke event (*χ* ^2^-test, *χ* ^2^ (1,N = 756) = 4.04, p = 0.04) and significantly more neurons encoded current trial relative chosen value in males compared to females during the outcome event (*χ* ^2^-test, *χ* ^2^(1,N = 756) = 5.30, p = 0.02). There were no sex differences in the encoding of relative chosen value for any other task events (*χ* ^2^-test, p>0.05). **e**. The proportion of neurons significantly encoding current relative side value during the nose poke, lever presentation, and outcome periods and the upcoming relative side value during rewarded and unrewarded outcomes. Significantly more neurons encoded upcoming relative side value in males compared to females during rewarded outcomes (*χ* ^2^-test, *χ* ^2^(1,N = 756) = 5.35, p = 0.02).

## Supplementary Tables

**Supplementary Table 1:** Mixed-effects logistic regression of return probability (**Figure 1b**): *return_probability ∼ Sex + Previous_outcome + Trial + Sex:Prevous_outcome + Sex:Trial + Previous_outcome:Trial + Sex:Previous_outcome:Trial + (1 + Previous_outcome+Trial + Previous_outcome:Trial|subject)*. The model was fit with effects coding. Sex: female = -1, Male = 1, Previous outcome: reward = -1, no reward = 1. Trial number was Z-scored. Significance of each coefficient was assessed with an F-test. n = 35 females, 37 males.

| | $\beta \pm$ standard error | F-stat (1, 816952) | p-value | 95% CI [upper, lower] | |
| --- | --- | --- | --- | --- | --- |
| Intercept | 1.024±0.03 | 1101.81 | 1.93E-241 | 0.96 | 1.08 |
| Previous_outcome | -0.879±0.02 | 1378.12 | 2.14E-301 | -0.93 | -0.83 |
| Trial | 0.110±0.01 | 80.66 | 2.68E-19 | 0.09 | 0.13 |
| Sex | -0.046±0.03 | 2.22 | 0.14 | -0.11 | 0.01 |
| Previous_outcome:Trial | -0.071±0.01 | 130.69 | 2.91E-30 | -0.08 | -0.06 |
| Previous_outcome:Sex | -0.003±0.02 | 0.01 | 0.91 | -0.05 | 0.04 |
| Trial:Sex | -0.030±0.01 | 5.90 | 0.02 | -0.05 | -0.01 |
| Previous_outcome:Trial:Sex | 0.004±0.01 | 0.44 | 0.51 | -0.01 | 0.02 |

**Supplementary Table 2:** Mixed-effects regression of trial initiation latency (**Figure 1c**): *trial_initiation_latency ∼ Sex + Previous_outcome + Sex:Previous_outcome + (1+Previous_outcome| subject)*. The model was fit with effects coding. Sex: female = -1, Male = 1, Previous outcome: reward = -1, no reward = 1. Significance of each coefficient was assessed with F-tests using the Satterthwaite method to estimate degrees of freedom. n = 35 females, 37 males.

| | $\beta \pm$ standard error | F-stat | DF1 | DF2 | p-value | 95% CI [upper, lower] | |
| --- | --- | --- | --- | --- | --- | --- | --- |
| Intercept | 3.99±0.10 | 1448.73 | 1 | 71.60 | 2.99E-49 | 3.78 | 4.19 |
| Previous_outcome | 1.30±0.08 | 247.10 | 1 | 70.30 | 1.04E-24 | 1.13 | 1.46 |
| Sex | -0.51±0.10 | 23.38 | 1 | 71.60 | 7.37E-06 | -0.71 | -0.30 |
| Sex:Previous_outcome | -0.31±0.08 | 13.98 | 1 | 70.30 | 3.74E-04 | -0.47 | -0.15 |

**Supplementary Table 3:**
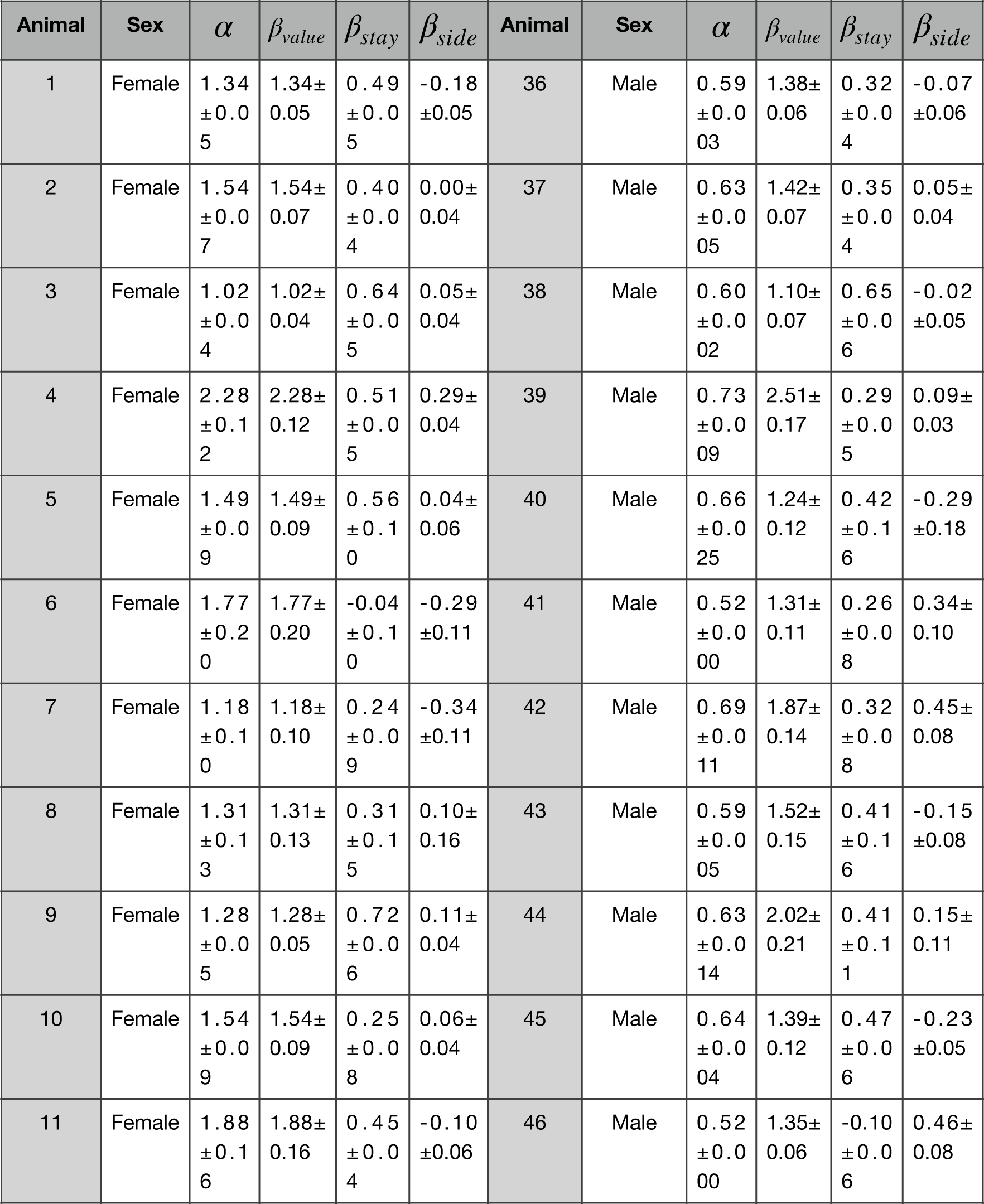
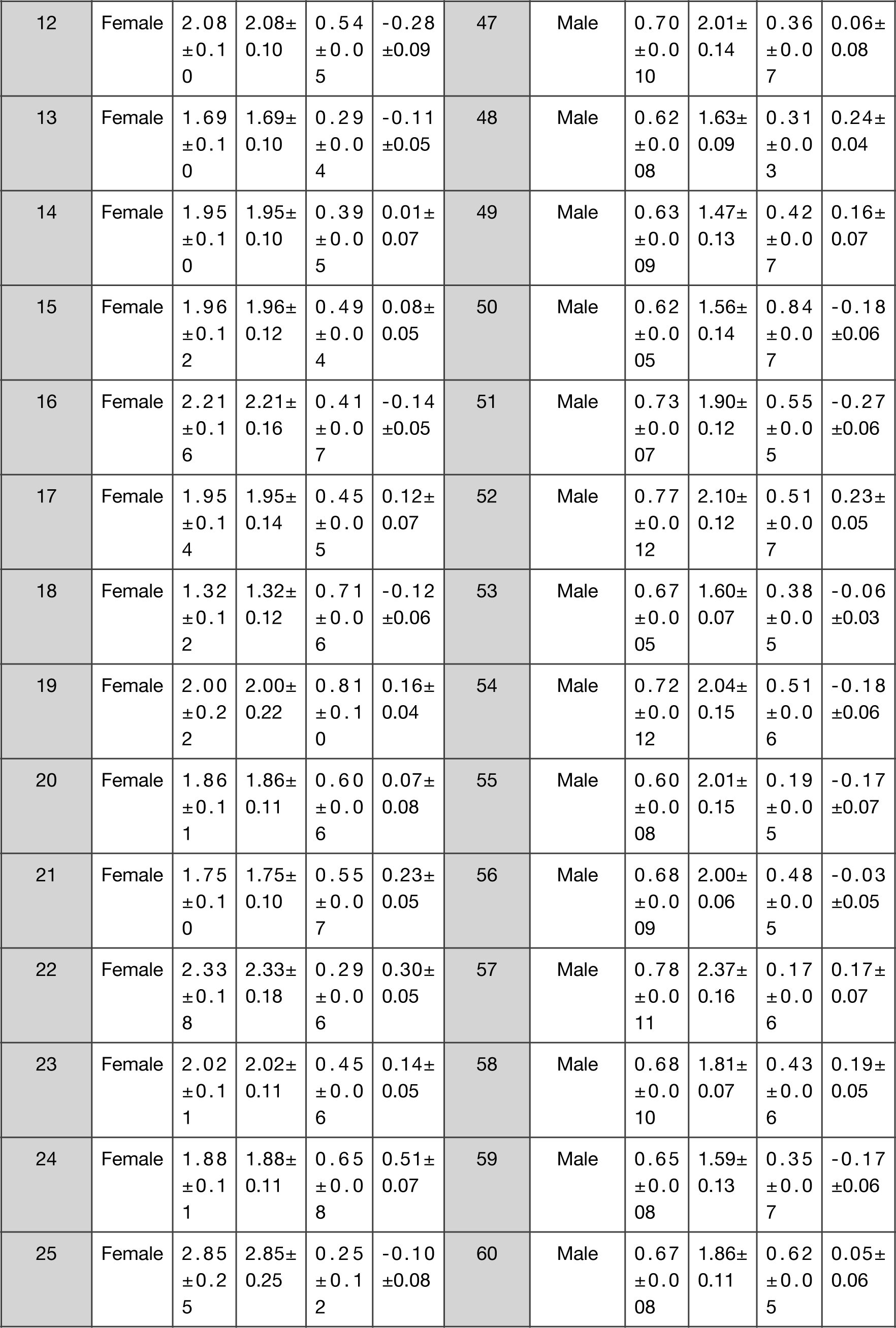
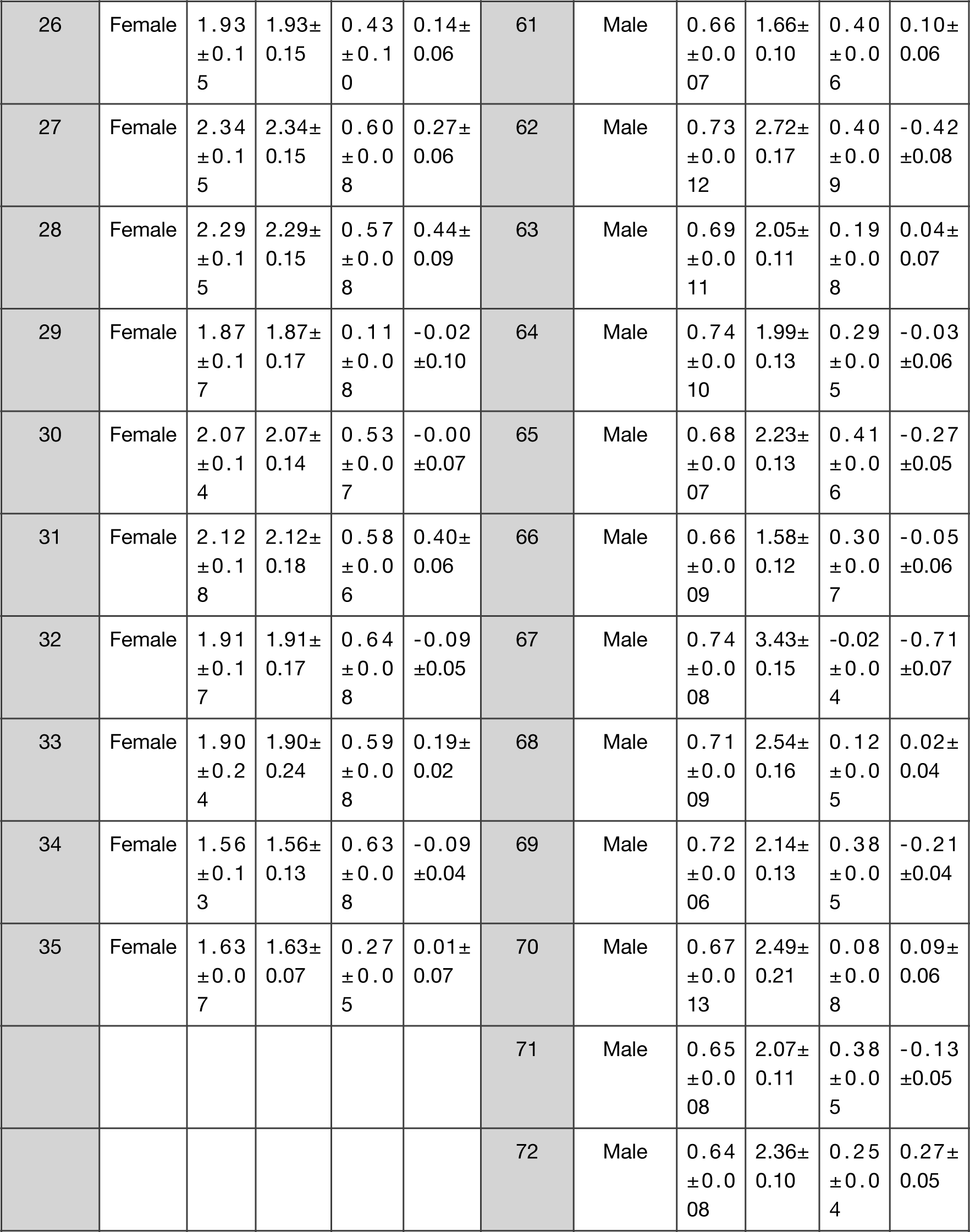
Fitted Q-learning parameters (**Figure 1e**). Parameter estimates are reported as mean±standard error of the mean across sessions for each mouse. ɑ is the learning rate and determines how much the *Q*-value for the chosen action is updated on each trial. *β_value_* is the inverse temperature which determines the degree to which the relative side value influences choice, *β_stay_* captures the tendency of the mice to repeat their previous choice and *β_side_* captures side bias.

**Supplementary Table 4:** Mixed-effects, logistic regression of choice (**Figure 1g**): *choice ∼ Sex + Relative_side_value + Trial + Sex:Relative_side_value + Sex:Trial + Relative_side_value:Trial + Sex:Relative_side_value:trial + (1+Relative_side_value+Trial + Relative_side_value:Trial|subject).* Trials were divided into 9 quantile bins of relative side value for each animal. Trial number was Z-scored and sex and relative side value bin were categorical variables. Model was fit with effects coding. (Female = -1, Male = 1). n=35 females, 37 males.

| | $\beta \pm \text{standard error}$ | T-statistic<br>(1,816924) | p-value | 95% CI [upper, lower] | |
| --- | --- | --- | --- | --- | --- |
| Intercept | 0.030±0.02 | 1.29 | 0.20 | -0.02 | 0.08 |
| Trial | 0.018±0.01 | 2.03 | 0.04 | 0.00 | 0.03 |
| Sex | -0.031±0.02 | -1.36 | 0.17 | -0.08 | 0.01 |
| Relative_side_value bin_1 | 1.871±0.05 | 34.42 | 2.28E-259 | 1.76 | 1.98 |
| Relative_side_value bin_2 | 1.530±0.05 | 29.92 | 1.38E-196 | 1.43 | 1.63 |
| Relative_side_value bin_3 | 0.829±0.02 | 37.39 | 1.02E-305 | 0.79 | 0.87 |
| Relative_side_value bin_4 | 0.258±0.01 | 18.09 | 3.72E-73 | 0.23 | 0.29 |
| Relative_side_value bin_5 | -0.046±0.01 | -3.83 | 1.26E-04 | -0.07 | -0.02 |
| Relative_side_value bin_6 | -0.309±0.02 | -16.71 | 1.17E-62 | -0.35 | -0.27 |
| Relative_side_value bin_7 | -0.857±0.02 | -36.26 | 1.21E-287 | -0.90 | -0.81 |
| Relative_side_value bin_8 | -1.504±0.05 | -30.03 | 5.69E-198 | -1.60 | -1.41 |
| Trial:Sex | -0.009±0.01 | -1.02 | 0.31 | -0.03 | 0.01 |
| Trial:Relative_side_value bin_1 | 0.202±0.02 | 9.67 | 4.04E-22 | 0.16 | 0.24 |
| Trial:Relative_side_value bin_2 | 0.178±0.02 | 10.80 | 3.45E-27 | 0.15 | 0.21 |
| Trial:Relative_side_value bin_3 | 0.061±0.01 | 5.98 | 2.23E-09 | 0.04 | 0.08 |
| Trial:Relative_side_value bin_4 | 0.002±0.01 | 0.17 | 0.86 | -0.02 | 0.02 |
| Trial:Relative_side_value bin_5 | -0.007±0.01 | -0.77 | 0.44 | -0.02 | 0.01 |
| Trial:Relative_side_value bin_6 | -0.044±0.01 | -5.19 | 2.14E-07 | -0.06 | -0.03 |
| Trial:Relative_side_value bin_7 | -0.088±0.01 | -8.84 | 9.66E-19 | -0.11 | -0.07 |
| Trial:Relative_side_value bin_8 | -0.166±0.02 | -9.21 | 3.4E-20 | -0.20 | -0.13 |
| Sex:Relative_side_value bin_1 | -0.049±0.05 | -0.91 | 0.36 | -0.16 | 0.06 |
| Sex:Relative_side_value bin_2 | -0.018±0.05 | -0.36 | 0.72 | -0.12 | 0.08 |
| Sex:Relative_side_value bin_3 | -0.013±0.02 | -0.57 | 0.57 | -0.06 | 0.03 |
| **Sex:Relative_side_value bin_4** | 0.010±0.01 | 0.69 | 0.49 | -0.02 | 0.04 |
| **Sex:Relative_side_value bin_5** | 0.016±0.01 | 1.33 | 0.18 | -0.01 | 0.04 |
| **Sex:Relative_side_value bin_6** | 0.029±0.02 | 1.55 | 0.12 | -0.01 | 0.06 |
| **Sex:Relative_side_value bin_7** | -0.004±0.02 | -0.19 | 0.85 | -0.05 | 0.04 |
| **Sex:Relative_side_value bin_8** | 0.003±0.05 | 0.05 | 0.96 | -0.10 | 0.10 |
| **Trial:Sex:Relative_side_value bin_1** | -0.005±0.02 | -0.24 | 0.81 | -0.05 | 0.04 |
| **Trial:Sex:Relative_side_value bin_2** | -0.035±0.02 | -2.15 | 0.03 | -0.07 | 0.00 |
| **Trial:Sex:Relative_side_value bin_3** | -0.005±0.01 | -0.50 | 0.62 | -0.03 | 0.02 |
| **Trial:Sex:Relative_side_value bin_4** | -0.001±0.01 | -0.14 | 0.89 | -0.02 | 0.02 |
| **Trial:Sex:Relative_side_value bin_5** | 0.008±0.01 | 0.89 | 0.38 | -0.01 | 0.02 |
| **Trial:Sex:Relative_side_value bin_6** | 0.003±0.01 | 0.35 | 0.73 | -0.01 | 0.02 |
| **Trial:Sex:Relative_side_value bin_7** | 0.005±0.01 | 0.53 | 0.60 | -0.01 | 0.02 |
| **Trial:Sex:Relative_side_value bin_8** | 0.016±0.02 | 0.89 | 0.37 | -0.02 | 0.05 |

**Supplementary Table 5:** Mixed-effects regression of trial initiation latency (**Figure 1h**): *trial_initiation_latency ∼ Sex + Relative_chosen_value + Weight + Trial + Sex:Relative_chosen_value + Relative_chosen_value:Weight + Relative_chosen_value:Trial + Sex:Weight + Sex:Trial + Sex:Relative_chosen_value:Trial + Sex:Relative_chosen_value:Weight + (1+Relative_chosen_value + Trial + Weight + Relative_chosen_value:Trial + Relative_chosen_value:Weight|Subject) + (1+Relative_chosen_value + Trial + Relative_chosen_value:Trial|Session:Subject)*. Session by session weights and trial were Z-scored. Model was fit with effects coding (Female = -1, Male = 1). n = 35 females, 37 males.

| | $\beta \pm$ standard error | T-statistic (1, 784158) | p-value | 95% CI [upper, lower] | |
| --- | --- | --- | --- | --- | --- |
| **Intercept** | 4.533±0.15 | 29.76 | 1.55E-194 | 4.23 | 4.83 |
| **Trial** | 0.980±0.05 | 19.76 | 6.39E-87 | 0.88 | 1.08 |
| **Sex** | -0.426±0.15 | -2.80 | 0.01 | -0.72 | -0.13 |
| **Relative_chosen_value bin_1** | 2.047±0.17 | 12.05 | 1.85E-33 | 1.71 | 2.38 |
| **Relative_chosen_value bin_2** | 1.364±0.11 | 12.00 | 3.41E-33 | 1.14 | 1.59 |
| **Relative_chosen_value bin_3** | -1.452±0.12 | -12.34 | 5.44E-35 | -1.68 | -1.22 |
| **Weight** | -0.295±0.10 | -2.95 | 0.003 | -0.49 | -0.10 |
| **Trial:Sex** | -0.101±0.05 | -2.04 | 0.04 | -0.20 | 0.00 |
| **Trial:Relative_chosen_value bin_1** | 0.828±0.06 | 13.14 | 2E-39 | 0.70 | 0.95 |
| **Trial:Relative_chosen_value bin_2** | 0.331±0.06 | 5.79 | 7.24E-09 | 0.22 | 0.44 |
| **Trial:Relative_chosen_value bin_3** | -0.525±0.05 | -10.53 | 6.32E-26 | -0.62 | -0.43 |
| **Sex:Relative_chosen_value bin_1** | -1.060±0.17 | -6.24 | 4.28E-10 | -1.39 | -0.73 |
| **Sex:Relative_chosen_value bin_2** | -0.840±0.11 | -7.39 | 1.45E-13 | -1.06 | -0.62 |
| **Sex:Relative_chosen_value bin_3** | 0.827±0.12 | 7.03 | 2.08E-12 | 0.60 | 1.06 |
| **Sex:Weight** | -0.358±0.10 | -3.58 | 3.4E-04 | -0.55 | -0.16 |
| **Relative_chosen_value bin_1:Weight** | 0.776±0.11 | 7.27 | 3.63E-13 | 0.57 | 0.99 |
| **Relative_chosen_value bin_2:Weight** | 0.551±0.09 | 6.02 | 1.78E-09 | 0.37 | 0.73 |
| **Relative_chosen_value bin_3:Weight** | -0.587±0.08 | -7.07 | 1.59E-12 | -0.75 | -0.42 |
| **Trial:Sex:Relative_chosen_value bin_1** | -0.110±0.06 | -1.74 | 0.08 | -0.23 | 0.01 |
| **Trial:Sex:Relative_chosen_value bin_2** | -0.036±0.06 | -0.63 | 0.53 | -0.15 | 0.08 |
| **Trial:Sex:Relative_chosen_value bin_3** | 0.088±0.05 | 1.76 | 0.08 | -0.01 | 0.19 |
| **Trial:Sex:Relative_chosen_value bin_1:Weight** | -0.192±0.11 | -1.79 | 0.07 | -0.40 | 0.02 |
| **Trial:Sex:Relative_chosen_value bin_2:Weight** | -0.101±0.09 | -1.10 | 0.27 | -0.28 | 0.08 |
| **Trial:Sex:Relative_chosen_value bin_3:Weight** | 0.102±0.08 | 1.23 | 0.22 | -0.06 | 0.26 |

**Supplementary Table 6:** In addition to modulation by relative chosen value, total value also significantly affected trial initiation latencies in a sex-dependent manner. This was assessed with a linear mixed-effects regression: *trial_initiation_latency ∼ Sex + Trial + Relative_chosen_value + Total_value + Sex*Relative_chosen_value + Sex*Total_value + (1|subject).* Trial, relative total value and relative chosen value were Z-scored and sex was a categorical variable (female = -1, male = 1). Significance of each term was assessed with F-tests using the Satterthwaite method to estimate degrees of freedom. N = 35 females, 37 males.

| | $\beta \pm \text{standard error}$ | F-stat | Numerat<br>or<br>degrees<br>of<br>freedom | Denominator<br>degrees of<br>freedom | p-value | 95% CI [upper,<br>lower] | |
| --- | --- | --- | --- | --- | --- | --- | --- |
| **Intercept** | 4.129±0.12 | 1225.56 | 1 | 71.80 | 7.248E-47 | 3.90 | 4.36 |
| **Trial** | 0.833±0.02 | 2104.44 | 1 | 790903.15 | 0 | 0.80 | 0.87 |
| **Sex** | -0.537±0.12 | 20.73 | 1 | 71.74 | 2.108E-05 | -0.77 | -0.31 |
| **Total_value** | -1.035±0.06 | 298.72 | 1 | 69.75 | 6.51E-27 | -1.15 | -0.92 |
| **Relative_chosen_value** | -0.856±0.04 | 444.51 | 1 | 67.85 | 1.685E-31 | -0.94 | -0.78 |
| **Sex:Total_value** | 0.264±0.06 | 19.41 | 1 | 69.75 | 3.727E-05 | 0.15 | 0.38 |
| **Sex:Relative_chosen_value** | 0.127±0.04 | 9.82 | 1 | 67.83 | 0.003 | 0.05 | 0.21 |

**Supplementary Table 7:**
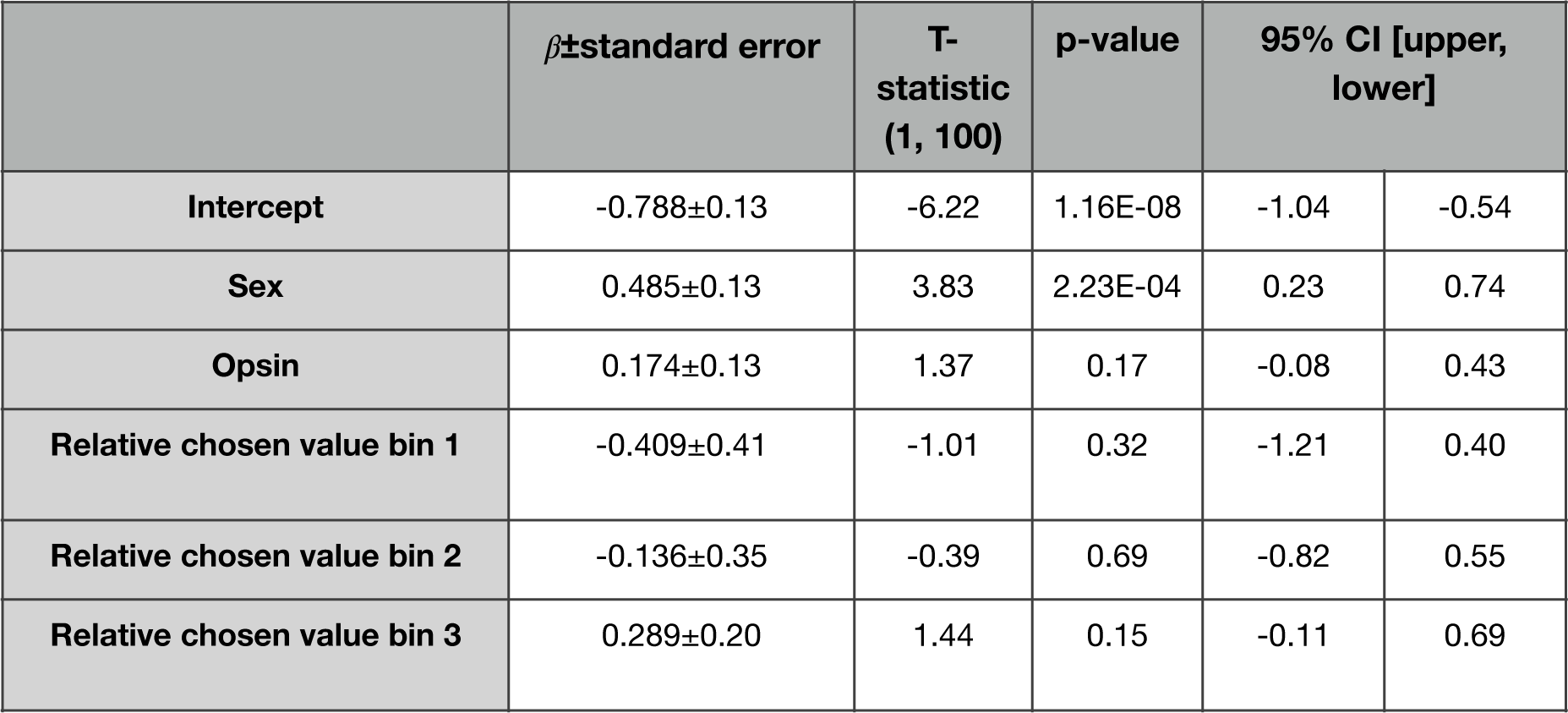
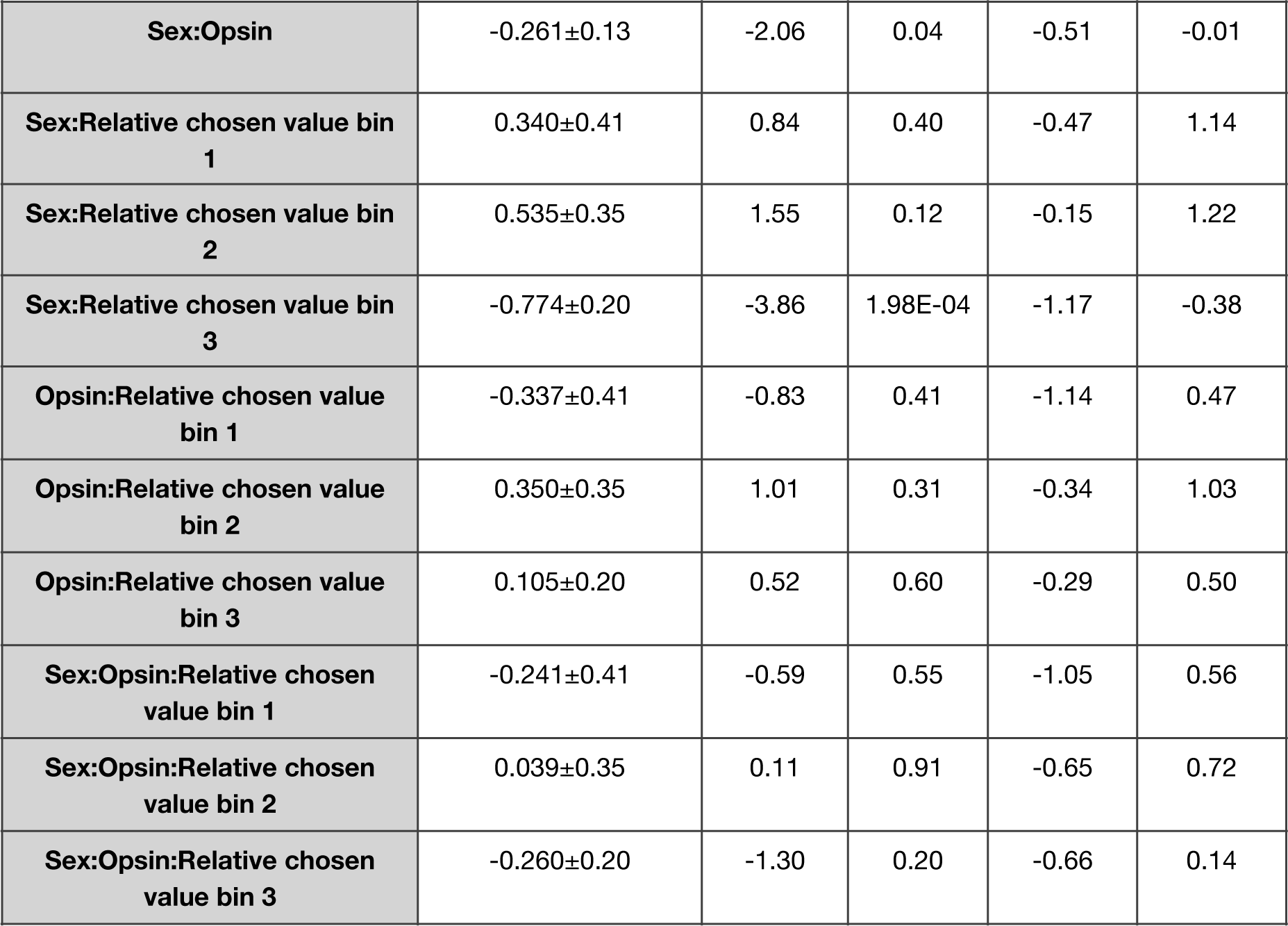
A linear mixed-effects regression was used to predict the difference in trial initiation latencies between laser and no-laser trials in 4 quantiles of relative chosen value (**Figure 2e**). *Trial_initiation_latency (laser-no laser) ∼ Sex + Opsin + Relative_chosen_value + Sex:Opsin + Sex:Relative_chosen_value + Opsin:Relative_chosen_value + Sex:Opsin:Relative_chosen_value + (1+Relative_chosen_value|Subject).* Model was fit with effects coding. (Female = -1, male = 1; NpHR = -1, EYFP = 1). n = 8 female NpHR, 9 male NpHR, 6 female EYFP, 6 male EYFP.

| | $\beta \pm \text{standard error}$ | T-statistic<br>(1, 100) | p-value | 95% CI [upper,<br>lower] | |
| --- | --- | --- | --- | --- | --- |
| **Intercept** | -0.788±0.13 | -6.22 | 1.16E-08 | -1.04 | -0.54 |
| **Sex** | 0.485±0.13 | 3.83 | 2.23E-04 | 0.23 | 0.74 |
| **Opsin** | 0.174±0.13 | 1.37 | 0.17 | -0.08 | 0.43 |
| **Relative chosen value bin 1** | -0.409±0.41 | -1.01 | 0.32 | -1.21 | 0.40 |
| **Relative chosen value bin 2** | -0.136±0.35 | -0.39 | 0.69 | -0.82 | 0.55 |
| **Relative chosen value bin 3** | 0.289±0.20 | 1.44 | 0.15 | -0.11 | 0.69 |
| **Sex:Opsin** | -0.261±0.13 | -2.06 | 0.04 | -0.51 | -0.01 |
| **Sex:Relative chosen value bin 1** | 0.340±0.41 | 0.84 | 0.40 | -0.47 | 1.14 |
| **Sex:Relative chosen value bin 2** | 0.535±0.35 | 1.55 | 0.12 | -0.15 | 1.22 |
| **Sex:Relative chosen value bin 3** | -0.774±0.20 | -3.86 | 1.98E-04 | -1.17 | -0.38 |
| **Opsin:Relative chosen value bin 1** | -0.337±0.41 | -0.83 | 0.41 | -1.14 | 0.47 |
| **Opsin:Relative chosen value bin 2** | 0.350±0.35 | 1.01 | 0.31 | -0.34 | 1.03 |
| **Opsin:Relative chosen value bin 3** | 0.105±0.20 | 0.52 | 0.60 | -0.29 | 0.50 |
| **Sex:Opsin:Relative chosen value bin 1** | -0.241±0.41 | -0.59 | 0.55 | -1.05 | 0.56 |
| **Sex:Opsin:Relative chosen value bin 2** | 0.039±0.35 | 0.11 | 0.91 | -0.65 | 0.72 |
| **Sex:Opsin:Relative chosen value bin 3** | -0.260±0.20 | -1.30 | 0.20 | -0.66 | 0.14 |

**Supplementary Table 8:** ACC-DMS inhibition affected trial initiation latencies similarly in the first and second half of the session. The full model was: *trial initiation latency (no laser-laser) ∼ Sex + Relative_chosen_value + Opsin + Time_bin + Sex:Relative_chosen_*value *+ Sex:Opsin + Sex:Time_bin + Relative_chosen_value: Opsin + Relative_chosen_value: t ime_bin + Opsin: Tme_bin + Sex:Relative_chosen_value:Opsin + Sex:Relative_chosen_value:Time_bin + Sex:Opsin:Time_bin + Relative_chosen_value:Opsin:Time_bin + Sex:Relative_chosen_value:Time_bin:Opsin + (1+relative_chosen_value|subject).* Model was fit with effects coding (Female = -1, male = 1, NpHR = -1, EYFP = 1, First half = 1, second half = -1). n = 8 female NpHR, 9 male NpHR, 6 female EYFP, 6 male EYFP.

| | $\beta \pm$ standard error | T-statistic (1, 200) | p-value | 95% CI [upper, lower] | |
| --- | --- | --- | --- | --- | --- |
| **Intercept** | -0.954±0.23 | -4.13 | 5.24E-05 | -1.41 | -0.50 |
| **Sex** | 0.335±0.23 | 1.45 | 0.15 | -0.12 | 0.79 |
| **Opsin** | 0.252±0.23 | 1.09 | 0.28 | -0.20 | 0.71 |
| **Relative chosen value bin 1** | -0.407±0.53 | -0.77 | 0.44 | -1.45 | 0.64 |
| **Relative chosen value bin 2** | -0.132±0.49 | -0.27 | 0.79 | -1.10 | 0.84 |
| **Relative chosen value bin 3** | 0.108±0.37 | 0.29 | 0.77 | -0.62 | 0.84 |
| **Time bin** | 0.390±0.21 | 1.90 | 0.06 | -0.02 | 0.80 |
| **Sex:Opsin** | -0.521±0.23 | -2.26 | 0.03 | -0.98 | -0.07 |
| **Sex:Relative chosen value bin 1** | 1.090±0.53 | 2.06 | 0.04 | 0.05 | 2.13 |
| **Sex:Relative chosen value bin 2** | -0.290±0.49 | -0.59 | 0.56 | -1.26 | 0.68 |
| **Sex:Relative chosen value bin 3** | -0.822±0.37 | -2.22 | 0.03 | -1.55 | -0.09 |
| **Opsin:Relative chosen value bin 1** | -0.180±0.53 | -0.34 | 0.73 | -1.22 | 0.86 |
| **Opsin:Relative chosen value bin 2** | 0.504±0.49 | 1.03 | 0.31 | -0.46 | 1.47 |
| **Opsin:Relative chosen value bin 3** | -0.057±0.37 | -0.15 | 0.88 | -0.79 | 0.67 |
| **Sex:Time bin** | -0.216±0.21 | -1.05 | 0.29 | -0.62 | 0.19 |
| **Opsin:Time bin** | -0.051±0.21 | -0.25 | 0.80 | -0.46 | 0.35 |
| **Relative chosen value bin 1: time bin** | -0.547±0.36 | -1.53 | 0.13 | -1.25 | 0.16 |
| **Relative chosen value bin 2:time bin** | 0.160±0.36 | 0.45 | 0.65 | -0.54 | 0.86 |
| **Relative chosen value 3:time bin** | 0.193±0.36 | 0.54 | 0.59 | -0.51 | 0.90 |
| **Sex:Opsin:Relative chosen value time bin 1** | 0.624±0.53 | 1.18 | 0.24 | -0.42 | 1.67 |
| **Sex:Opsin:Relative chosen value time bin 2** | -1.474±0.49 | -3.00 | 0.00 | -2.44 | -0.51 |
| **Sex:Opsin:Relative chosen value time bin 3** | 0.143±0.37 | 0.39 | 0.70 | -0.59 | 0.87 |
| **Sex:Opsin:Time bin** | 0.185±0.21 | 0.90 | 0.37 | -0.22 | 0.59 |
| **Sex:Relative chosen value bin 1: time bin** | -0.961±0.36 | -2.70 | 0.01 | -1.66 | -0.26 |
| **Sex:Relative chosen value bin 2: time bin** | 0.007±0.36 | 0.02 | 0.98 | -0.70 | 0.71 |
| **Sex:Relative chosen value bin 3: time bin** | 0.300±0.36 | 0.84 | 0.40 | -0.40 | 1.00 |
| **Opsin:Relative chosen value bin 1:Time bin** | -0.100±0.36 | -0.28 | 0.78 | -0.80 | 0.60 |
| **Opsin:Relative chosen value bin 2:Time bin** | 0.005±0.36 | 0.01 | 0.99 | -0.70 | 0.71 |
| **Opsin:Relative chosen value bin 3:Time bin** | 0.010±0.36 | 0.03 | 0.98 | -0.69 | 0.71 |
| **Sex:Opsin:Relative chosen value bin 1:Time bin** | -0.709±0.36 | -1.99 | 0.05 | -1.41 | -0.01 |
| **Sex:Opsin:Relative chosen value bin 2:Time bin** | 0.791±0.36 | 2.22 | 0.03 | 0.09 | 1.49 |
| **Sex:Opsin:Relative chosen value bin 3:Time bin** | 0.051±0.36 | 0.14 | 0.89 | -0.65 | 0.75 |

**Supplementary Table 9:** A linear mixed-effects regression was used to predict the difference in choice between laser and no-laser trials in 9 quantiles of relative side value (**Figure 2f**). *Right_choice (laser-no laser) ∼ Sex + Opsin + Relative_side_value + Sex:Opsin + Sex:Relative_side_value + Opsin:Relative_side_value + Sex:Opsin:Relative_side_value + (1+Relative_side_value|Subject).* Model was fit with effects coding (Female = -1, male = 1; NpHR = -1, EYFP = 1). n = 8 female NpHR, 9 male NpHR, 6 female EYFP, 6 male EYFP.

| | $\beta \pm$ standard error | T-stat(1, 225) | p-values | 95% CI [upper, lower] | |
| --- | --- | --- | --- | --- | --- |
| **(Intercept)** | -0.002±0.01 | -0.29 | 0.77 | -0.01 | 0.01 |
| **Sex** | -0.004±0.01 | -0.58 | 0.56 | -0.02 | 0.01 |
| **Opsin** | -0.003±0.01 | -0.50 | 0.62 | -0.02 | 0.01 |
| **Relative side value bin 1** | -0.012±0.02 | -0.78 | 0.44 | -0.04 | 0.02 |
| **Relative side value bin 2** | 0.014±0.02 | 0.94 | 0.35 | -0.02 | 0.04 |
| **Relative side value bin 3** | 0.002±0.01 | 0.16 | 0.87 | -0.02 | 0.03 |
| **Relative side value bin 4** | -0.007±0.01 | -0.44 | 0.66 | -0.04 | 0.02 |
| **Relative side value bin 5** | -0.026±0.02 | -1.58 | 0.11 | -0.06 | 0.01 |
| **Relative side value bin 6** | -0.003±0.01 | -0.19 | 0.85 | -0.03 | 0.03 |
| **Relative side value bin 7** | 0.011±0.02 | 0.72 | 0.47 | -0.02 | 0.04 |
| **Relative side value bin 8** | 0.006±0.01 | 0.41 | 0.69 | -0.02 | 0.04 |
| **Sex:Opsin** | -0.001±0.01 | -0.16 | 0.87 | -0.01 | 0.01 |
| **Sex:Relative side value bin 1** | -0.010±0.02 | -0.66 | 0.51 | -0.04 | 0.02 |
| **Sex:Relative side value bin 2** | 0.007±0.02 | 0.49 | 0.63 | -0.02 | 0.04 |
| **Sex:Relative side value bin 3** | -0.002±0.01 | -0.16 | 0.87 | -0.03 | 0.02 |
| **Sex:Relative side value bin 4** | 0.020±0.01 | 1.37 | 0.17 | -0.01 | 0.05 |
| **Sex:Relative side value bin 5** | 0.005±0.02 | 0.29 | 0.77 | -0.03 | 0.04 |
| **Sex:Relative side value bin 6** | 0.006±0.01 | 0.39 | 0.69 | -0.02 | 0.03 |
| **Sex:Relative side value bin 7** | -0.008±0.02 | -0.52 | 0.60 | -0.04 | 0.02 |
| **Sex:Relative side value bin 8** | 0.010±0.01 | 0.64 | 0.52 | -0.02 | 0.04 |
| **Opsin:Relative side value bin 1** | 0.013±0.02 | 0.81 | 0.42 | -0.02 | 0.04 |
| **Opsin:Relative side value bin 2** | -0.008±0.02 | -0.54 | 0.59 | -0.04 | 0.02 |
| **Opsin:Relative side value bin 3** | 0.015±0.01 | 1.18 | 0.24 | -0.01 | 0.04 |
| **Opsin:Relative side value bin 4** | -0.028±0.01 | -1.87 | 0.06 | -0.06 | 0.00 |
| **Opsin:Relative side value bin 5** | 0.006±0.02 | 0.38 | 0.71 | -0.03 | 0.04 |
| **Opsin:Relative side value bin 6** | 0.008±0.01 | 0.54 | 0.59 | -0.02 | 0.04 |
| **Opsin:Relative side value bin 7** | -0.001±0.02 | -0.03 | 0.97 | -0.03 | 0.03 |
| **Opsin:Relative side value bin 8** | 0.005±0.01 | 0.34 | 0.73 | -0.02 | 0.03 |
| **Sex:Opsin:Relative side value bin 1** | -0.003±0.02 | -0.22 | 0.83 | -0.03 | 0.03 |
| **Sex:Opsin:Relative side value bin**<br>**2** | 0.005±0.02 | 0.36 | 0.72 | -0.02 | 0.04 |
| **Sex:Opsin:Relative side value bin**<br>**3** | -0.002±0.01 | -0.15 | 0.88 | -0.03 | 0.02 |
| **Sex:Opsin:Relative side value bin**<br>**4** | -0.009±0.01 | -0.59 | 0.55 | -0.04 | 0.02 |
| **Sex:Opsin:Relative side value bin**<br>**5** | 0.023±0.02 | 1.40 | 0.16 | -0.01 | 0.05 |
| **Sex:Opsin:Relative side value bin**<br>**6** | 0.027±0.01 | 1.93 | 0.06 | 0.00 | 0.06 |
| **Sex:Opsin:Relative side value bin**<br>**7** | -0.018±0.02 | -1.19 | 0.23 | -0.05 | 0.01 |
| **Sex:Opsin:Relative side value bin**<br>**8** | -0.014±0.01 | -0.97 | 0.33 | -0.04 | 0.02 |

**Supplementary Table 10:** A linear mixed-effects regression was used to predict optogenetically-evoked EPSCs in DMS: *current ∼ Sex + MSN_type + Sex*MSN_type + (1|Subject) + (1|Pair:Subject)* (**Figure 3D**). All regressors were categorical variables and the model was fit with effects coding. Significance of fixed effects was assessed with F-tests using the Satterthwaite method to estimate degrees of freedom. n = 3 females, 9 pairs, 6 males, 16 pairs.

| | $\beta \pm$ standard error | F-stat | DF_1 | DF_2 | p-value | 95% CI [upper, lower] | |
| --- | --- | --- | --- | --- | --- | --- | --- |
| **Intercept** | -266.7±44.62 | 35.728 | 1 | 8.5 | 0.0003 | -356.52 | -176.89 |
| **Sex** | -11.15 ±44.62 | 0.062 | 1 | 8.5 | 0.81 | -100.96 | 78.664 |
| **MSN type** | 22.459±23.17 | 0.9398 | 1 | 25 | 0.34 | -24.17 | 69.09 |
| **Sex\*MSN type** | 36.204±23.16<br>7 | 2.4421 | 1 | 25 | 0.13 | -10.43 | 82.84 |

## Methods

### Mice

All procedures were approved by the Institutional Animal Care and Use Committee at Princeton University and were performed in accordance with the Guide for the Care and Use of Laboratory Animals. One hundred and ten mice were used for these experiments: 58 C57BL/6J mice (strain 000664, The Jackson Laboratory; 24 males and 34 females), 16 Drd1-cre mice (EY262Gsat, MMRRC-UCD; 7 female, 9 male), 15 Drd2-cre mice (ER44Gsat, MMRRC-UCD; 8 female, 7 male),12 A2a-cre mice (KG139Gsat, MMRRC-UCD; 6 female, 6 male), and 9 Drd1a-tdTomato mice (strain 016204, The Jackson Laboratory; 3 female, 6 male). Mice were between the ages of 2 and 5 months at the beginning of all experiments. Prior to surgery, mice were housed in groups of 2-5 mice per cage. All mice used in the imaging experiments were singly-housed after surgery to prevent damage to implants from cagemates. Mice used for optogenetic and electrophysiology experiments remained group-housed after surgery. Mice were housed in a reversed light/dark cycle with lights on from 8:30 pm - 8:00 am. All experiments were performed during the dark cycle.

### Mouse behavior

Mice were trained to perform a probabilistic reversal learning task as described in^45, 46^. Beginning 3 to 5 days prior to the onset of training, mice were handled by experimenters daily and placed on water restriction. During the remainder of the experiments, water was mostly delivered during behavioral sessions and mice received 1-2 mL of water per day. Mice were assessed daily for clinical signs of dehydration, and body weight was maintained at >=80% of *ad libitum* weight. If these criteria were not met, mice were given extra water until they recovered.

Training was performed in 21 x 18 cm operant chambers (Med Associates, ENV-307W). Two retractable levers (Med Associates, ENV-312-W) were positioned on either side of a central nose poke (Med Associates, ENV-313-M). A reward port was positioned below the nose poke. Reward spout contact was recorded with either a contact lickometer (Med Associates, ENV-250) or a capacitive sensor (https://playground.arduino.cc/Main/CapacitiveSensor/). A speaker was located on the opposite wall for the delivery of ∼80 dB auditory cues signaling rewarded (CS+; 5 kHz pure tone) and unrewarded (CS-; white noise) outcomes. The task was controlled with custom MedPC IV (Med Associates) scripts.

Mice were trained in 3 stages. The following trial structure was consistent across training stages. A trial began with illumination of the central nose poke. When the mouse entered the nose poke, the left and right levers extended with a delay drawn from a uniform distribution between 0.05 and 1 second in 50 ms intervals. During the first 2 stages of training there was no time limit for pressing the lever. In the final version of the task, mice had 15 s to press one of the levers. Once the mouse pressed a lever, both were retracted and after a delay drawn from the same distribution as above, the outcome was presented. On rewarded trials, 6 *μ*L of a 10% sucrose solution was delivered to the central reward port in conjunction with the reward cue which played for 0.5 s. On unrewarded trials, 0.5 s of the no reward cue was presented. The start of reward consumption was defined as the first contact the mouse made with the reward port following reward delivery and the end of reward consumption was defined as when mice disengaged from the reward port for >250 ms. After the end of reward consumption or unrewarded outcome presentation, a 2.5 s intertrial interval (ITI) began, which ended with illumination of the central nose poke, indicating that the mouse was able to initiate the next trial.

On the first day of training, the central nose poke was baited with ¼ of a crushed fruit loop. Once the mouse entered the nose poke, the levers remained extended until one was pressed. Reward was delivered following all lever presses during a 1 hour session. Next, to train mice to perform the nose poke-lever press sequence and to prevent development of a side-bias, either lever press was rewarded 100% of the time, unless that lever had been pressed more than 5 times in a row. If a particular lever was pressed more than 5 times in a row, reward would not be delivered for that lever press until the mouse pressed the alternate lever. After the alternate lever press, both levers were again rewarded 100% of the time. After mice received >100 rewards in a single hour-long session, they were moved to the final version of the task. This stage lasted for 5.0 ± 3.3 sessions (mean ± standard deviation across mice, n = 100 mice).

In the final version of the probabilistic reversal learning task, one lever was rewarded with a probability of 0.7 (high-probability lever) and the other with a probability of 0.1 (low-probability lever). The identity of the high probability lever reversed after 10 high-probability presses plus a random number of trials drawn from the geometric distribution:

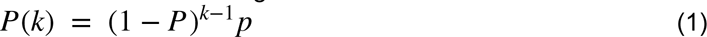

Where *P*(*k*) is the probability of a block transition after *k* trials and is the probability of switching for each trial, which was set to 0.4. Blocks were 18.49 ± 4.9 trials long (mean ± standard deviation) and there were 16.19 ± 2.00 blocks per 1 hour session and 26.5 ± 5.03 blocks per two hour session (mean ± standard deviation). In this stage, mice were required to press one of the levers within 15 s of their presentation, otherwise it was considered an abandoned trial and the levers were retracted and the ITI began (0.15 ± 0.13% of trials, n=100 mice). These trials were excluded from analysis. In order to increase the number of inactivation trials without extended virus expression times, after initial training, mice used for the optogenetic experiment were run in 2 hour sessions with a piece of their chow in the operant chamber.

### Human behavior

#### Subjects

Subjects (158 female, 235 male, 1 declined to report, 1 non-binary, 19-70 years old, 1 subject did not report their age) were recruited in December, 2020 and January-February, 2022 with Amazon Mechanical Turk with the aid of CloudResearch services^130^. Participants were eligible if they resided in the United States and passed CloudResearch’s data quality filters. Subjects were asked “What is your gender?” and could respond “Male”, “Female”, “Rather not say” or in a blank where participants could self-report their gender identity. Only subjects who reported their gender as male or female and passed performance criteria were included in analysis. Subjects were excluded if they completed fewer than 200 trials, did not discriminate between rewarded and unrewarded outcomes (the difference between stay probability following rewarded and unrewarded trials was less than 0.1) and/or they had high amounts of bias (greater than 40% bias). 209 males and 141 females passed these performance criteria and were included for analysis. Trial initiation latencies were z-scored within subject to account for variability in computer and internet connection speeds. Subjects were between 19 and 70 years of age (mean ± standard deviation: females 43.91 ± 12.46 years; males 38.66 ± 11.21 years). All subjects reported speaking English “well” or “very well.” At the end of the experiment, subjects were compensated $3.50 for their participation, which took 25 minutes on average. All subjects accepted the informed consent approved by the Institutional Review Board of Princeton University (#7392) prior to beginning of the task.

#### Task

Before beginning the session, subjects were presented with instructions for how to initiate trials and make choices and were told that on each trial they would receive either 10 points or 0 points for their choice. They were instructed that their goal was to maximize the total number of points received but they were not given any instruction about the reward probabilities or that these probabilities reversed throughout the session. Subjects then performed 4 practice trials before onset of the session.

Each trial began with the presentation of a cross (+) in the center of the screen indicating that they could initiate the trial by pressing the spacebar. If they did not initiate the trial within 10 seconds a warning message was displayed informing them that if they continued to time out, the experiment would end and they would be removed from the study. Subjects failed to initiate 0.11±0.33% (mean±standard deviation, n=350) of trials. Trial initiation was followed by a random, variable delay between 0.05 and 1s after which 2 circles were displayed on the right and left of the screen. Subjects indicated their choice with an ‘a’ or ‘l’ key press. On each trial, one circle was rewarded with a probability of 0.6 and the other with a probability of 0.1. Subjects had 15 seconds to make a choice, otherwise the trial ended and the ITI began (0.01 ± 0.06% of trials, mean±standard deviation). These trials were excluded from analysis. After subjects made a choice, the outcome was presented following a random delay between 0.05 and 1s. On rewarded trials “+10 points” was displayed on the screen and a 0.5s bell sound was played. On unrewarded trials “+0 points” was displayed on the screen and 0.5s buzzer sound was played. This was followed by a 1.5s ITI, after which the cross was displayed on the screen again, indicating that subjects could initiate the next trial. Subjects were not cued as to which option had the higher probability of reward and the reward probabilities reversed probabilistically according to the same rule as in the mouse task described above.

Because the study was conducted online, to help encourage subjects to focus on the task, the session was divided into 4, 5 minute periods. Between each period, subjects were allowed to take an untimed break and choose when to restart the session. At the beginning of each block the high probability choice was randomly assigned meaning that trials from the previous block were not informative as to the current location of the high probability choice.

### Stereotaxic surgery

Mice were anesthetized with 1.5-2% isofluorane and placed in a stereotaxic frame (David Kopf Instruments). Mice were given 2 doses of Meloxicam (10 mg/kg of body weight, the first before surgery and the second ∼24 hours later) as well as 1 dose of Baytril before surgery (5 mg/kg of body weight). After asepsis, the skull was exposed and cleaned of connective tissue with a bone scraper (Fine Science Tools). Craniotomies were drilled over the ACC (0.5mm anterior, 0.4 mm lateral to bregma) and the DMS (0.75mm anterior, 1.5mm lateral to bregma)^131^.

For optogenetic experiments (**Figure 2**), to specifically target neurons in the ACC that project to the DMS, we used a microsyringe (Micro4, World Precision Instruments) to inject 500 nL of retroAAV-Cre (retroAAV-EF1a-Cre-WPRE-hGHpA, Princeton Neuroscience Institute Vector Core, ∼2.40E14 parts/mL) into the DMS at 2.55mm ventral, 0.5 mm anterior and 0.4 mm lateral to bregma bilaterally. Additionally, we injected 500 nL of AAV5-EF1a-DIO-eNPHR3.0-EYFP-WPRE-hGH (Princeton Neuroscience Institute Vector Core, ∼1.6E12 parts/mL) or AAV5-EF1a-DIO-EYFP-hGHpA (Princeton Neuroscience Institute Vector Core, ∼1.5E12 parts/mL) bilaterally into the ACC at a depth of 1.5mm, 0.3mm lateral and 0.5 mm anterior from bregma.

We then implanted ferrules attached to optical fibers (300 μm core diameter, 0.37 NA, > 70% transmission) bilaterally at a 10° angle at 0.58mm lateral, 0.5mm anterior and 1mm ventral to bregma with metabond followed by dental cement darkened with carbon powder. The locations of the implants are shown in **Supplementary Figure 6**.

For ex-vivo recordings of EPSCs in the DMS (**Figure 3**), mice were injected unilaterally in the ACC with 250 nL of AAV5-CamKIIa-hChR2(H134R)-eYFP-WPRE (University of Pennsylvania Vector Core, ∼4.8E13 parts/mL) at 0.3 mm lateral, 0.5 mm anterior and 1.5 mm ventral from bregma.

For imaging experiments (**Figure 4-6**; n = 6 females, 9 males), we injected 500 nL of retroAAV-EF1a-Cre-WPRE-hGHpA (Princeton Neuroscience Institute Vector Core, ∼2.40E+14 parts/mL) into the DMS at 2.55mm ventral to bregma. Additionally, we injected 300-500 nL of AAV5-CAG-Flex-GCamp6f-WPRE-SV40 (University of Pennsylvania Vector Core or Addgene, ∼8.75E+12 parts/mL) into the ACC at a depth of 1.5mm. After removal of the dura, we implanted either a 1 mm diameter, ∼4.3 mm length prism probe (1050-004601, Inscopix; implanted at 0.7mm anterior, 0.5 mm lateral and 1.75 ventral to bregma at least 45 minutes after injection; n = 4 females, 5 males), or a 0.5 or 1 mm diameter, ∼4 mm length lens (1050-004595, Inscopix; implanted at 0.5 mm anterior, 0.5 mm lateral and 1.5 ventral to bregma; n = 2 females, 4 males), implanted 1 week later following aspiration of a thin layer of tissue at the surface. The lens or prism probe was held in place with a layer of metabond (Parkell) followed by dental cement darkened with carbon powder. A metal headpost was attached at the back of the skull with metabond. Finally, a plastic cap was glued to the implant to protect the lens. Two to four weeks after the implantation surgery, mice were anesthetized with 1-2% isofluorane and placed in the stereotaxic holder. A baseplate (Inscopix) attached to the miniature microscope (nVista 2.0 or nVista 3.0, Inscopix) was lowered over the lens until the GCaMP6f fluorescence was visible and neurons were in focus. The baseplate was then secured to the implant with dental cement darkened with carbon powder and covered with a baseplate cover. The locations of the implants are shown in **Supplementary Figure 11**. One mouse’s implant location could not be recovered.

### Optogenetic inactivation

After recovery from implantation surgery (at least 5 days), mice were put back on water restriction and retrained on the task in 2-hour sessions and habituated to the tether. Approximately 4 weeks after injection, once performance had stabilized, ∼ 5 mW of 532 nm light (Shanghai Laser and Optics & Co) was delivered bilaterally on a random subset of trials (10-15% of trials; **Figure 2**, **Supplementary** **Figures 6-10**). Light onset was triggered by the CS presentation (5-7.5% of trials) and stayed on for 2 s terminating with a 200 ms ramp off of power to reduce post-inhibitory rebound^132, 133^. Because both rewarded and unrewarded outcomes were explicitly cued, there was no ambiguity in the result of their choice (i.e. a possibility that reward is delayed) from the onset of inhibition. In subsets of mice, light delivery was also triggered by the nose poke at trial start and stayed on for 2 s (5% of trials; n=8 males and 10 females) or during the ITI (7.5% of trials), starting 2 seconds after the outcome presentation and terminating around the time of the nose poke (n=10 males and 9 females). Laser offset was jittered by 0.05-1 s so that offset was less contingent on the animal’s action. Laser sessions alternated with non-laser sessions across days.

### Ex vivo electrophysiology

Three to 4 weeks following injection of AAV5-CamKIIa-hChR2(H134R)-eYFP-WPRE virus in Drd1a-tdTomato mice, mice were anesthetized with an intraperitoneal injection of Euthasol (0.06ml/30g). Mice were decapitated and the brain was extracted. After extraction, the brain was immersed in ice-cold NMDG ACSF (92 mM NMDG, 2.5 mM KCl, 1.25 mM NaH2PO4, 30 mM NaHCO3, 20 mM HEPES, 25 mM glucose, 2 mM thiourea, 5 mM Na-ascorbate, 3 mM Na-pyruvate, 0.5 mM CaCl2·4H2O, 10 mM MgSO4·7H2O, and 12 mM N-Acetyl-L-cysteine; pH adjusted to 7.3-7.4) for 2 minutes. Afterwards coronal slices (300 µm) containing the ACC and DMS were sectioned using a vibratome (VT1200s, Leica, Germany) and then incubated in NMDG ACSF at 34°C for approximately 14 minutes. Slices were then transferred into a holding solution of HEPES ACSF (92 mM NaCl, 2.5 mM KCl, 1.25 mM NaH2PO4, 30 mM NaHCO3, 20 mM HEPES, 25 mM glucose, 2 mM thiourea, 5 mM Na-ascorbate, 3 mM Na-pyruvate, 2 mM CaCl2·4H2O, 2 mM MgSO4·7H2O and 12 mM N-Acetyl-l-cysteine, bubbled at room temperature with 95% O2/ 5% CO2) for at least 45 mins until recordings were performed.

Whole-cell recordings (**Figure 3** and **Supplementary Figure 11**) were performed using a Multiclamp 700B (Molecular Devices, Sunnyvale, CA) using pipettes with a resistance of 4-7 MOhm. For EPSC recordings, pipettes were filled with an internal solution containing 100 mM cesium gluconate, 0.6 mM EGTA, 10 mM HEPES, 5 mM NaCl, 20 mM TEA, 4 mM Mg-ATP, 0.3 mM Na-GTP and 3 mM QX 314 with the pH adjusted to 7.2 with CsOH. For EPSP recordings, pipettes were filled with an internal solution containing 120 mM potassium gluconate, 0.2 mM EGTA, 10 mM HEPES, 5 mM NaCl, 1 mM MgCl2, 2 mM Mg-ATP and 0.3 mM NA-GTP with the pH adjusted to 7.2 with KOH. The osmolarity of internal solutions was adjusted to 289 mmol kg−1 with sucrose. During recordings, slices were perfused with a recording ACSF solution (120 mM NaCl, 3.5 mM KCl, 1.25 mM NaH2PO4, 26 mM NaHCO3, 1.3 mM MgCl2, 2 mM CaCl2 and 11 mM D-(+)-glucose, and was continuously bubbled with 95% O2/5% CO2). Infrared differential interference contrast–enhanced visual guidance was used to select neurons that were 3–4 cell layers below the surface of the slices. MSNs were identified by the presence or absence of td-Tomato using a fluorescence microscope (Scientifica SliceScope Pro 1000; LED: SPECTRA X light engine (Lumencor)). The recording solution was delivered to slices via superfusion driven by peristaltic pump (flow rate of 4-5 ml/min) and was held at room temperature. In voltage clamp experiments, the neurons were held at −70 mV, and the pipette series resistance was monitored throughout the experiments by hyperpolarizing steps of -10 mV with each sweep. If the series resistance changed by >20 % during the recording, the data were discarded. Whole-cell currents were filtered at 1 kHz and digitized and stored at 10 KHz (Clampex 10; MDS Analytical Technologies). All experiments were completed within 4 hours after slices were made to maximize cell viability and consistency.

Optogenetically evoked EPSCs or EPSPs were recorded in the presence of picrotoxin (100 µM) in the recording ACSF solution. Sequential paired recordings were performed to control for differences in ChR2 expression levels, or differences in relative position to terminal expression. Photostimulation parameters were 474 nm and 0.008-0.075 mW/mm^2^ for durations of 0.5-3 ms, at 30 s inter-stimulation intervals. Photostimulation parameters were adjusted between pairs to achieve consistent responses, but parameters were kept the same within a given pair. For sequential paired recordings, MSNs were selected from the same field of view and approximately the same depth in the slice. The order of MSN subtype alternated between pairs to control for possible effects of rundown. For most cells, 10 EPSCs/EPSPs were recorded. EPSC and EPSP amplitudes were measured as the absolute value of the peak of the baseline-subtracted EPSC or EPSP.

### Single-photon calcium imaging

After implantation of a baseplate to secure the microscope, mice were habituated for imaging experiments with a dummy microscope and tether until their task performance was similar to before surgery. Prior to imaging, mice were briefly restrained using the implanted head bar to attach the microscope to the baseplate and focus the imaging field. Calcium imaging data was acquired with nVista HD software (Inscopix) or Inscopix Data Acquisition Software (Inscopix) at 15-20 frames per second. To synchronize imaging data with behavioral events, we recorded TTL sync pulses from the microscope and TTLs generated by the MedPC software and a SuperPort output module (Med Associates) with an RZ5D BioAmp processor from Tucker-Davis Technologies (**Figure 4-6, Supplementary Figure 12-14**).

### Estrous Tracking

To determine the stage of the estrous cycle throughout behavioral testing (**Supplementary Figure 3**), vaginal lavage was conducted as previously described^134, 135^ in 14 females. At the end of behavioral testing each day, an experimenter elevated the base of the tail of the mouse and 20uL of normal saline was flushed in and out of the vaginal opening 1-2 times using a pipette. The cell sample was then placed onto a glass slide and viewed under brightfield illumination on an EVOS M5000 microscope at 20X magnification within 10 minutes, before the sample dried.

The estrous cycle stage on each day was manually determined based on the proportion of leukocytes, cornified epithelial cells, and nucleated epithelial cells as well as cyclicity according to established protocols ^134–136^. Classifications were assigned by consensus between 3 experimenters. Briefly, proestrus was defined by the presence of mostly nucleated epithelial cells with some cornified epithelial cells. Estrus was dominated by cornified epithelial cells. Metestrus was identified by the presence of mostly cornified epithelial cells and leukocytes, with fewer nucleated epithelial cells. Samples from animals in diestrus contained primarily leukocytes. Stages of the cycle were grouped into follicular phase (proestrus/estrus), when estradiol levels are high, and luteal phase (metestrus/diestrus), when estradiol levels are low. One mouse was excluded from analysis because she did not cycle during the experiment.

### Histology

Mice were deeply anesthetized with 0.1 mL of Euthasol administered via intraperitoneal injection and transcardially perfused with 1x phosphate-buffered saline (PBS) followed by 4% paraformaldehyde in PBS. Brains were removed and post-fixed overnight in 4% paraformaldehyde and transferred to 30% sucrose in PBS for cryoprotection. The forebrain was then sectioned in 40 µm thick coronal slices with a cryostat (Leica Microsystems CM3050S). Sections were mounted with 1:2500 DAPI in Fluoromount G (Southern Biotech) or Fluoromount G with DAPI (Southern Biotech). To verify virus expression and implant location, whole sections containing the ACC were imaged with a Hamamatsu NanoZoomer S60. To confirm nuclear exclusion of GCaMP6f and produce **Figure 2a** and **Figure 4a**, sections were imaged with a Leica TCS SP8 confocal microscope. To confirm implant locations (**Supplementary Figure 6 and 12**), the section with the center of the lesion was identified. The dorsoventral and mediolateral position of the lesion as well as the distance between landmarks to estimate a scaling factor (distance from corpus callosum to cortical surface and midline to corpus callosum) was measured using NDPview (Hamamatsu) and aligned with the mouse atlas^131^.

### Behavior analysis

#### Data selection

Analyses of sex differences in performance of the task (**Figure 1**, **Supplementary** **Figures 1**, **Supplementary Figure 2, Supplementary Figure 3a**) used data from the post-surgery, no-laser, 2-hour sessions of mice from the optogenetic inactivation experiment (**Figure 2**) as well as mice with injections and implants in the DMS from a separate study. To select data from sessions after the mice had habituated to the tether, only sessions with at least 200 trials and a 10% difference in return probability following reward and no reward were included. Additionally, to account for differences in the time individuals became sated within a session, we only included trials until mice had not initiated a new trial for at least 5 minutes. Quantiles of relative side value, relative chosen value and total value were calculated across all sessions for each mouse.

#### Fitting trial initiation latency distributions

To analyze trial initiation latency distributions (**Supplementary Figure 1a-b**, **Figure 2c-d**, **Supplementary Figure 7a, Supplementary Figure 8**), we fit each animal’s trial initiation latencies with a shifted inverse gaussian distribution^137^ using the Matlab function *mle*. For optogenetics experiments we fit the laser and non-laser trials separately. Comparisons between males and females were performed with two-sided Wilcoxon rank sum tests and with two-sided Wilcoxon signed rank tests to compare laser and control parameter estimates.

*Q-learning model*:

We fit a trial-by-trial Q-learning model to choice data from all mice with a hierarchical model^47^. The models were initialized with *Q*-values of 0 for each action and updated on each trial according to:

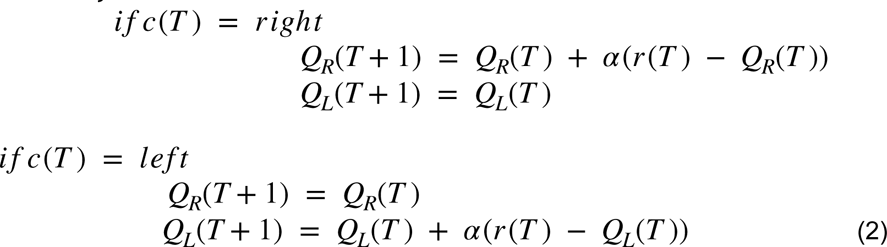

where *c*(*t*) is the choice on trial, *T*, *Q_R_* is the action value for the right choice, *Q_L_* is the action value for the left choice, *r* (*T*)is 0 or 1 to indicate if reward was received on trial, *T*, and *α* is a free learning rate parameter that is between 0 and 1. The probability of choosing choice *c*(*T*) was:

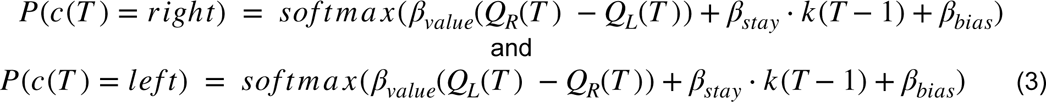

where *β_value_* is a free inverse temperature parameter, *β_stay_* is a free parameter to capture the tendency of mice to repeat their previous choice, *k* (*t* − 1) indicates the previous choice (-1 if right, 1 if left), and *β_bias_* is a free parameter to account for side bias.

The four free parameters were estimated separately for each session for each subject, and across sessions for each subject and jointly for the entire population (in a hierarchical random effects model), with group-level means and variance reflecting the distribution over sessions and over the population of subject-level parameters respectively. The parameters were estimated with Hamiltonian Monte Carlo implemented in the Stan programming language and the Python package PyStan (version 2.17.1.0^138^). We ran the model with 4 chains of 1000 iterations each (the first 250 were discarded for burn-in). The adapt_delta parameter was set to 0.99. We verified convergence by visual inspection and confirming that the potential scale reduction statistic Rhat^139^ was close to 1.0 for all parameters. Averages of the session-level parameters for all subjects are shown in **Supplementary Table 3**.

We then used the estimated parameters from each session to compute trial-by-trial estimates of *Q*-values for each action. We then calculated relative side value as the difference between *Q*-values for right and left choices, relative chosen value as the difference between *Q*-values for chosen and unchosen actions and total value as the sum of the *Q*-values for the two actions, to correlate with choice, trial initiation latencies and neural activity.

#### Statistical models of choice

Modulation of return probability by previous outcome was analyzed with a mixed-effects logistic regression (**Figure 1b** and **Supplementary Table 1**). Returning to the previously selected lever (return = 1, switch = 0) was predicted with fixed effects of sex, trial number, previous outcome and sex*previous outcome and random effects of subject (intercept and previous outcome).

To analyze the influence of value on choice (**Figure 1g**; **Supplementary Table 4**), for each subject, trials were divided into 9 quantile bins of relative side value. We fit a logistic mixed-effects model of choice (right = 1, left = 0), with sex, relative side value quantile bin, trial and sex*relative side value as fixed effects and random effects for subject (intercept and relative side value). Trial was z-scored and sex and value were categorical variables.

To analyze the effect of relative side value on choice in human subjects (**Supplementary Figure 5c-d**), for each subject, trials were divided into 9 quantile bins of relative side value. We fit a logistic mixed-effects regression of choice (right = 1, left = 0), with sex, relative side value, trial, age, sex*relative side value, age*relative side value, sex*age, and sex*relative side value*age as fixed effects and random effects for subject (intercept and relative side). Trial and age were z-scored and sex and value were categorical variables.

Models were fitted with the Matlab function, *fitglme,* using “effects” coding for categorical variables. Significant main effects and interactions were assessed with F-tests. Overall significance of relative side value and relative side value*sex was assessed with ssignificance tests on F-statistics calculated by comparing the full model with a model without all the value terms or without the value*sex terms.

#### Statistical models of trial initiation latency

Modulation of trial initiation latency by previous outcome was analyzed with linear mixed-effects models (**Figure 1c**; **Supplementary Table 2**). Trial initiation latency was averaged by subject based on whether the previous trial was rewarded for each mouse and predicted with sex, previous outcome and their interaction as fixed effects and random intercepts for subject.

To estimate the effect of relative chosen value on trial initiation latency (**Figure 1h**; **Supplementary Table 5**), trials were divided into 4 quantile bins of relative chosen value for each animal. Trial initiation latency was predicted using a linear mixed-effects model with sex, relative chosen value quantile bin, session weight, trial number, sex*relative chosen value quantile bin, weight*relative chosen value bin, trial*relative chosen value bin, sex*weight*relative chosen value bin and trial*weight*relative chosen value bin as fixed effects and random effects for subject (intercept, relative chosen value, trial, weight, relative chosen value*weight and relative chosen value*trial) and session (intercept, relative chosen value, trial, relative chosen value*weight).

To estimate the effect of total value on trial initiation latency (**Supplementary Figure 3a**), we fit the same model as **Figure 1h** and **Supplementary Table 5**, except with total value quantile bin instead of relative chosen value.

To estimate the effect of relative chosen value on trial initiation latency in human subjects, for each subject, we divided trials into 5 quantile bins of relative chosen value and averaged trial initiation latencies for each bin. We then fit a linear mixed effects regression on trial initiation latency using sex, relative chosen value bin, age, sex*relative chosen value bin, age*relative chosen value bin, sex*age and sex*age*relative chosen value bin as fixed effects and random intercepts for subject. Age was Z-scored across the population and value bin and gender were categorical variables (female = -1,male = 1). To perform post-hoc comparisons between males and females aged between 19 and 39 years old or 40 and 70 years old, we performed two-sided Wilcoxon rank sum tests.

To quantify the effect of both total value and relative chosen value on trial initiation latency (**Supplementary Table 6**), we fit a linear mixed-effects regression with total value, relative chosen value, sex, total value*sex and relative chosen value*sex as fixed effects and random effects for subject (intercept, relative chosen value and total value). Relative total value and relative chosen value were Z-scored and sex was a categorical variable (female = -1, male = 1).

All models were fitted with the *fitlme* function in Matlab using “effects” coding for categorical variables. Significant main effects and interactions were assessed with F-tests using the Satterthwaite method for estimating degrees of freedom. Post-hoc F-tests on contrasts between males and females were performed with the Matlab function, *coefTest,* using the Satterthwaite method to estimate degrees of freedom. Overall significance of relative chosen value and relative chosen value*sex was assessed with significance tests on F-statistics calculated by comparing the full model with a model without the value terms or without the value*sex terms, using the Satterthwaite method to estimate degrees of freedom.

#### Statistical models of the effect of optogenetic inhibition on choice and trial initiation latencies

Tso analyze sex differences in the effect of inhibition on value-dependent trial initiation latency (**Figure 2e**, **Supplementary Figure 7b**), we fit a linear mixed-effects regression to the difference between trial initiation latencies in quantile bins on laser and non-laser trials with relative chosen value bin, sex, opsin, relative chosen value*sex, relative chosen value*opsin, sex*opsin and sex*relative chosen value*opsin as fixed effects and random intercepts for subject. To analyze sex differences in the effect of inhibition on the modulation of trial inititation latency by total value (**Supplementary Figure 3b**), we fit a mixed-effects regression to the difference between trial initiation latencies in quantile bins on laser and non-laser trials with total value bin, sex, opsin, relative total value*sex, relative total value*opsin, sex*opsin and sex*relative total value*opsin fixed effects and random intercepts for subject. Models were fitted with the *fitlme* function in Matlab with “effects” dummy coding. Significant main effects and interactions were assessed with F-tests using the Satterthwaite method for estimating degrees of freedom. Overall effects of value, sex*value, sex*opsin, opsin*value and sex*value*opsin were assessed with significance tests on F-statistics calculated by comparing the full model with a model without the value, sex*value, sex*opsin, opsin*value or sex*value*opsin terms, using the Satterthwaite method to estimate degrees of freedom. Post-hoc comparisons between laser and non-laser trials were performed with two-sided Wilcoxon signed rank tests for each group.

To determine if there was an effect of inhibition on value-dependent choice (**Figure 2f**, **Supplementary Figure 7c**), we divided trials into quantiles of relative side value. We fit a logistic mixed-effects regression on choice (right=1,left =0) with sex, opsin, relative side value, laser, and all 2- and 3-way interactions as fixed effects and random effects for subject (intercept and relative side value). Models were fitted with the Matlab function, *fitglme,* using “effects” coding for categorical variables. Significant main effects and interactions were assessed with F-tests.

### Analysis of Ca^2+^ imaging data

#### Image processing

Images were spatially downsampled by 4-pixel bins and motion-corrected using Mosaic or the Inscopix Data Processing software (Inscopix). Following motion correction, we used the CNMFe algorithm^60^ to extract individual neurons and their fluorescence traces. Only neurons with an estimated event rate of four spikes per minute or higher were analyzed (307/314 neurons in females, 449/458 neurons in males). Fluorescence traces were Z-scored using the mean and standard deviation across the entire recording session.

#### Encoding models of calcium fluorescence

In order to determine how neural activity was modulated during the task (**>Figures 4-5** and **Supplementary Figure 13**), we built a linear encoding model to relate neural activity to each event^46, 140–145^. We used multiple linear regression analyses of each neuron with the fluorescence trace as the dependent variable and behavioral events as the independent variables. In order to account for delays in the relationship between changes in fluorescence and behavioral events, independent variables were generated by convolving a 25 degrees-of-freedom spline basis set with an 8 second duration with a binary vector of event times (1 at the time of the behavioral event and 0 otherwise). The spline basis set was generated with the Matlab package fdaM (https://www.psych.mcgill.ca/misc/fda/downloads/FDAfuns/R/inst/Matlab/fdaM/). The full model is

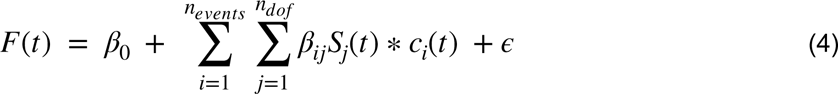

where *F*(*t*) is the Z-scored fluorescence at time, *t, β*_0_ is the intercept, ε is the error, *n_events_* is the number of behavioral events, *β_ij_* is the regression coefficient for the *j*th spline basis function and *i*th event, *S_j_* is the *j*th spline basis function, *n_dof_* is the degrees-of-freedom for the spline basis set (25) and *c_i_* is a binary vector the same length as *F* that is 1 at the times of event *i* and 0 otherwise. Thus, fluorescence is modeled as the convolution of the event time vector, *c_i_*(*t*), with a temporal kernel, *K_i_*(*t*), summed across events:

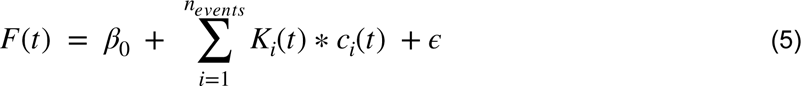

Where the temporal kernel for event is defined as:

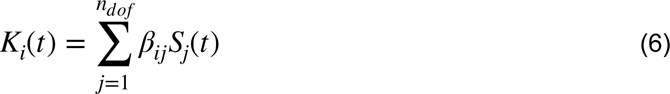

Temporal kernels for nose poke entry and lever press events spanned from 2 seconds before until 6 seconds after the event, and lever presentation and outcome-related kernels spanned 8 seconds from the time of the event. We estimated these temporal kernels with a linear regression using the Matlab function, *fitglm*.

To identify significant modulation of neural activity by a particular event (**Figures 4e-h, 5c-d**; details for specific models are below), we calculated an F-statistic comparing the fit of the full model to the fit of a reduced model without the predictors for that event. We then calculated the same F-statistic for the same comparison using 500 instances of shifted data generated by circularly shifting the fluorescence trace by a random integer. We then compared the F-statistic of the real data to the distribution generated from the shifted data to calculate a p-value. Neurons with p-values less than 0.01 were considered to significantly encode the event.

To determine whether a neuron significantly encoded outcome (**Figure 4e,f**), we estimated calcium fluorescence using “nose poke entry”, “lever presentation”, “ipsilateral lever press”, “contralateral lever press”, “outcome” and “CS-”. A neuron was considered to encode outcome if removal of the “CS-” event significantly worsened model fit. This allowed us to distinguish between neurons with significant changes in activity in response to both rewarded and unrewarded outcomes from those with significantly different activity during the two trial types. Neurons were then classified as “reward-preferring” or “no-reward preferring” based on the sign of the approximate integral of the “CS-” temporal kernel calculated with the *trapz* function in Matlab (**Supplementary Figure 13b**). Neurons with a negative integral were classified as “reward-preferring” and neurons with a positive integral were “no-reward preferring.”

Similarly, to identify neurons with outcome responses that were significantly different based on whether the upcoming choice was to stay with the previously selected lever, we estimated calcium fluorescence with “nose poke entry”, “lever presentation”, “ipsilateral lever press”, “contralateral lever press”, “reward delivery”, “reward delivery and stay”, “CS- and “CS- and stay” events. Significant modulation of reward and no reward responses were assessed based on removal of the “reward delivery and stay” and “CS- and stay” events, respectively (**Figure 4g,h**) and classification of “stay-preferring” versus “switch-preferring was based on the sign of the approximate integral of these temporal kernels (**Supplementary Figure 13d**, positive integrals indicated “stay-preferring” and negative “switch-preferring”). To determine significant choice encoding (**Figure 5c,d**) we fit a model with “nose poke entry”, “lever presentation”, “lever press”, “ipsilateral lever press”, “reward delivery” and “CS-” and identified neurons for which removal of the “ipsilateral lever press” event significantly affected fit. Neurons with a positive integral of the “ipsilateral lever press” temporal kernel were classified as “ipsi-preferring” and those with a negative integral were “contra-preferring” (**Supplementary Figure 13c**).

To determine how the transient, event-related responses of ACC-DMS neurons were modulated by trial-by-trial fluctuations in relative chosen value and relative side value, we fit bilinear models, which enabled multiplicative gain coefficients based on the trial-by-trial values to scale the temporal kernels for each event (**Figure 6** and **Supplementary Figure 14**)^146, 147^.

The trial-by-trial gain for event *i* is

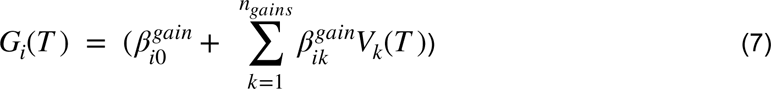

where 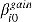 is the gain offset for event *i*, *n_gains_* is the number of trial-by-trial variables, 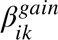 is the coefficient for trial-by-trial variable *k* and event *i*, *V_k_*(*T*) is the value of trial-by-trial variable *k* (e.g. relative chosen value) on trial *T*. Thus, the fluorescence on trial *T* is modeled as

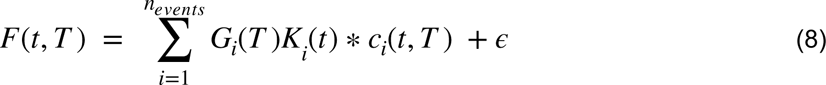

where *c_i_*(*t*, *T*) is a binary indicator of event times for event *i* on *T* trial and *K_i_*(*T*) is the temporal kernel as defined in equation 6.

We fit 3 separate models to evaluate the influence of relative chosen value (**Figure 6b,c**), relative side value (**Figure 6e,f**) or total value (**Supplementary Figure 3c,d**) on event-related neural activity. First, to estimate the effect of relative chosen value on event responses (**Figure 6b,c**), we considered 5 events: “nose poke entry”, “lever presentation”, “ipsilateral lever press”, “contralateral lever press” and “CS” (CS+ or CS-). For all events except CS, we fit 1 gain coefficient for relative chosen value on the current trial. For the CS event we fit 2 gain coefficients: current trial and next trial relative chosen value. To estimate the effect of relative side value on event responses (**Figure 6e,f**), we considered 5 events: “nose poke entry”, “lever presentation”, “lever press”, “reward delivery” and “CS-”. For all events except “reward delivery” and “CS-”, we fit 1 gain coefficient for relative side value on the current trial. For the outcome events we fit 2 gain coefficients: current trial and next trial relative side value. For the third model, to estimate the effect of total value on event responses (**Supplementary Figure 3c,d**), we considered 5 events: “nose poke entry”, “lever presentation”, “ipsilateral lever press”, “contralateral lever press” and “CS”. For all events except CS, we fit 1 gain coefficient for total value on the current trial. For the CS event we fit 2 gain coefficients: current trial and next trial total value.

To fit the model defined in equation 8, we iteratively estimated the temporal kernels, *K_i_*(*t*), and the value-dependent gains, *G_i_*(*T*), similar to^147^. On each iteration, we first kept the value-dependent gains fixed and estimated the temporal kernel, *K_i_*(*t*), with a linear regression using the Matlab function *fitlm*. On the first iteration, *G_i_*(*T*) was set to 1 and on subsequent iterations it was based on the gain coefficients estimated in the previous iteration. We next kept the temporal kernels, *K_i_*(*t*), fixed and estimated the value-dependent gains. When estimating the value-dependent gains, the temporal kernels were normalized by dividing the coefficients for the spline basis set, *β_ij_*, by the L2 norm of the kernel. This process was repeated 10 times.

In order to determine which neurons significantly encoded relative side value or relative chosen value (**Figure 6**), we fit the model to 1000 instances of shifted data generated by shifting the fluorescence trace by a random integer. We then compared the t-statistic for the gain coefficients estimated from the real data to a distribution of t-statistics generated with fits of the shifted data to determine if the contribution of that variable to the fit was significantly greater than chance. A significance level of 0.01 was used to classify neurons as significant.

## Acknowledgements

We would like to thank the Witten lab as well and N.D. Daw for comments and advice on this work. This research was funded by NYSCF and SCGB grant to IBW, and also ARO W911NF1710554 to IBW and the following NIH grants: U19 NS104648-01, R01 DA047869 and 5R01MH106689-02 to IBW and F32 MH112320 to J.C. IBW is a New York Stem Cell Foundation—Robertson Investigator

## Contributions

J.C. and I.B.W conceived the project and designed the experiments. J.C., A.R.M, W.T.F., C.H., A.B., S.O., B.M. and S.Z. collected the data. J.C. analyzed the data with support from C.A.Z, N.F.P and S.Z. J.C. and I.B.W. wrote the manuscript.

